# Tissue Reconstruction with Generative AI

**DOI:** 10.1101/2025.02.13.638045

**Authors:** Tingyang Yu, Chanakya Ekbote, Nikita Morozov, Antonio Herrera, Jiashuo Fan, Albert Dominguez Mantes, Salvatore Novello, Hilal Lashuel, Pascal Frossard, Stéphane d’Ascoli, Gioele La Manno, Maria Brbić

## Abstract

The spatial arrangement of cells is fundamental to their function, but single-cell RNA sequencing dissociates cells from their native spatial context. We present LUNA, a generative AI model that reconstructs tissue architecture from dissociated gene expression profiles by learning spatial priors from spatially resolved transcriptomics datasets. We validate LUNA by reconstructing the MERFISH mouse brain atlas comprising over 1.2 million cells, inferring the spatial locations of dissociated cells in a mouse central nervous system single-cell RNA-sequencing atlas, and enriching the Slide-tags data by imputing the locations of nuclei lost during profiling. LUNA generalizes to previously unseen cell types in the MERFISH mouse brain atlas and to pathological samples, identifying regions of disease-associated alterations in a mouse model of Parkinson’s disease. Our results establish generative tissue reconstruction as an approach for overcoming current technological limitations in spatial biology and advancing our understanding of tissue organization and function.

## Introduction

Single-cell RNA sequencing (scRNA-seq) technologies have enabled the high-throughput profiling of cellular transcriptomes, making it possible to identify and explore unique transcriptional profiles within individual cells [1, 2]. These technologies have revolutionized our understanding of cellular diversity, providing unprecedented insights into the complexity of cells and tissues [3–8]. However, a major limitation of scRNA-seq technologies is the loss of spatial context, as cells are dissociated from their native microenvironments during the sequencing process. The spatial context is crucial for understanding how cells interact with their neighbors and the surrounding microenvironment, influencing their roles in healthy tissue function and disease development. Although spatially resolved technologies aim to address this limitation [9–11], they are restricted in the number of genes that can be measured and are not always an option.

To address the limitations of individual technologies, computational methods have been developed to infer spatial context from gene expression of single-cells. Early approaches relied on in situ hybridization (ISH)-based reference atlases of landmark gene expression and leveraged them to infer the spatial location of cells in scRNA-seq datasets [12, 13]. More recently, reference mapping methods such as Tangram [14], CytoSPACE [15], STEM [16], scSpace [17], STALocator [18] and CeLEry [19], proposed mapping dissociated tissue data to reference spatial datasets. Although these methods have shown promising results, they rely on an exact spatial reference for mapping scRNA-seq datasets, thus primarily functioning as data integration tools. On the other hand, novoSpaRc [20, 21] enabled the spatial reconstruction of single-cell gene expression. However, novoSpaRc assumes that cells in close physical proximity exhibit similar gene expression profiles — an assumption that is often violated in complex tissue architectures.

Here, we present LUNA (Location reconstrUction using geNerative Ai), a generative AI model that reconstructs complex tissue structures from gene expression of cells by learning spatial priors over spatial transcriptomics datasets. LUNA learns cell representations that capture cellular interactions locally and globally across the entire tissue slice, enabled by an attention mechanism [22, 23] that takes into account interactions across all cells. LUNA operates as a diffusion model [24–26]: during training it learns to denoise corrupted cell coordinates, while during inference it starts from random noise and reconstructs the physical locations of cells solely from their gene expression.

We apply LUNA to reconstruct the MERFISH whole mouse brain atlas, which consists of 1.23 million cells spanning all brain regions [27], as well as the scRNA-seq mouse central nervous system atlas of 1.08 million cells [28]. LUNA effectively reconstructs mouse brain tissue across a wide range of brain regions and captures spatial gene expression patterns, including cells from previously unseen cell types without explicit training on those specific cell types. Beyond prediction, we use LUNA to identify genes that are the most predictive of the spatial structure, offering a means to link tissue architecture to its underlying transcriptional programs. Furthermore, LUNA enables the enrichment of Slide-tags datasets [29] by inferring the tissue locations of nuclei lost during cell profiling. LUNA correctly placed different subtypes of neuronal cells in mouse embryonic tissue, enriching the sparse Slide-tags data by up to 95.5%, as well as tumor cells belonging to spatially distinct compartments in a human metastatic melanoma sample. Finally, we generated a Xenium spatial transcriptomics dataset from the brains of wild-type control mice and *α*-synuclein-induced Parkinson’s disease model mice. LUNA successfully positioned clusters of cells with pathological *α*-synuclein, even though it was trained only on control samples, as confirmed by immunostaining. LUNA is highly scalable, with linear time and memory complexity relative to the number of input cells during the inference, and it can infer the spatial coordinates of tens of thousands of cells in less than a few minutes on a single GPU.

## Results

### Overview of LUNA

Given gene expressions of dissociated single cells, we aim to infer their spatial locations and reconstruct tissue structure (Fig. 1a). To achieve that, we designed a diffusion model [24–26], named LUNA, that generates complex tissue structures by predicting the physical locations of individual cells based on their gene expression. The key challenge in generating tissue structures arises from the inherent complexity of tissues, which are composed of a vast array of heterogeneous cells engaged in intricate cell-cell communication. LUNA overcomes these challenges by enabling each cell to “communicate” with all other cells guided by the learned weights, *i.e*., attention scores (Methods). In this way, LUNA learns cell embeddings that capture cellular interactions on both the global and local levels, allowing LUNA to directly map cell embeddings to spatial locations. During training, LUNA learns spatial priors on existing spatial transcriptomics data. In the inference stage, LUNA generates complex tissue structures solely from gene expression of cells.

**Figure 1.**
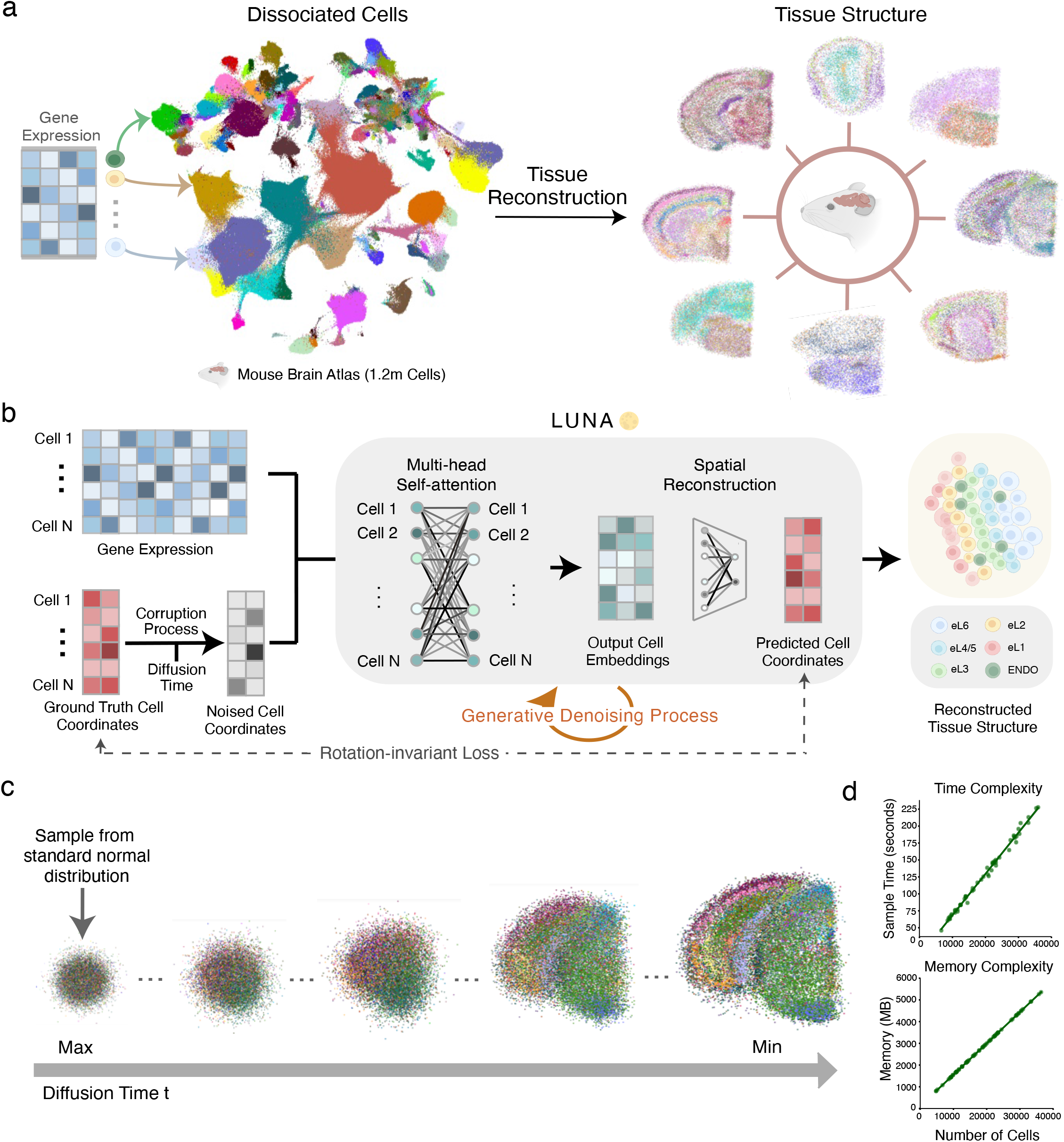
*(preceding page)*: LUNA is a generative AI model that positions cells into complex tissue structures from their gene expressions by learning spatial priors over existing spatially resolved datasets. **(a)** Given gene expressions of cells, the tissue reconstruction process involves predicting the complex tissue architecture from gene expression profiles of cells. We visualize the UMAP projection of cells based on their gene expression profiles (1, 122 genes) from the MERFISH whole mouse Allen Brain Cell (ABC) Atlas [27] (left). Each cell is colored according to one of 338 distinct subclasses. LUNA’s predictions of the spatial tissue structure is displayed on the right with eight slices randomly chosen from a total of 66 slices. LUNA was trained on spatial transcriptomics dataset of another mouse. **(b)** Overview of the training stage of LUNA. LUNA takes as input spatial transcriptomics data and corrupts the spatial locations of cells by adding noise. Using the attention mechanism, LUNA learns how to position each individual cell with respect to other cells based on their gene expressions. The loss function in LUNA ensures invariance to the rotations and translations of the predicted slice. **(c)** Visualization of the denoising during the inference stage in LUNA across different diffusion steps on an example slice (29, 079 cells) from the ABC atlas. The noise state is initialized from a standard normal distribution. Conditioned on the gene expressions of the cells, LUNA progressively removes noise at each diffusion step. **(d)** LUNA achieves linear time and memory complexity during inference relative to the number of input cells. Correlations between inference time and the number of input cells (top) and between reserved memory of the GPU and the number of input cells (bottom) across all 66 slices from the ABC atlas. Each point corresponds to a slice, correlating the cell count with either the time or memory used during the inference stage by LUNA. Icons in **(a-b)** were created using BioRender.

The neural network in LUNA consists of a multi-head self-attention mechanism [22, 23] that allows each cell to focus on gene expression of specific cells when making predictions (Fig. 1b). The cell embeddings are then mapped from the latent space to the physical space via a fully connected layer. To ensure robustness to rotations and translations of input slices, the loss function in LUNA is specially designed to be invariant to these transformations and preserve the relative spatial relationships of the cells (Methods).

The training stage of LUNA comprises two processes: corruption and denoising, both of which are repeated over a range of diffusion times (Methods). During the corruption process, a controlled amount of Gaussian noise is added to the ground truth cell coordinates. Using the noised cell coordinates generated from this process and conditioned on the gene expression data, LUNA then learns to recover the original cell coordinates through the denoising process. During the inference stage, LUNA takes gene expression profiles as model input and reconstructs tissue structures by denoising cell coordinates in an autoregressive manner. Initially, cell coordinates are generated from random Gaussian noise, and LUNA iteratively removes noise from these coordinates, predicting spatial locations of cells (Fig. 1c, Extended Data Fig. 1). LUNA is highly scalable, with linear time and memory complexity relative to the number of cells during the inference. It infers physical locations of tens of thousands of cells in just a few minutes on a single GPU. For example, LUNA inferred locations of 40, 000 cells from the Allen Brain Cell (ABC) MERFISH mouse brain atlas [27] in approximately four minutes using around 6 GB of GPU memory (Fig. 1d).

### LUNA accurately reconstructs the whole mouse brain atlas

LUNA effectively scales to atlas-scale large datasets and accurately reconstructed a whole mouse brain of the Allen Brain Cell (ABC) MERFISH mouse brain atlas [27]. We trained LUNA on 2.85 million cells across 147 slices from one mouse (Supplementary Note 1). We then applied it to infer locations of cells from the whole brain of another mouse consisting of 1.23 million cells and 338 identified subclasses in 66 slices (Fig. 2a).

**Figure 2.**
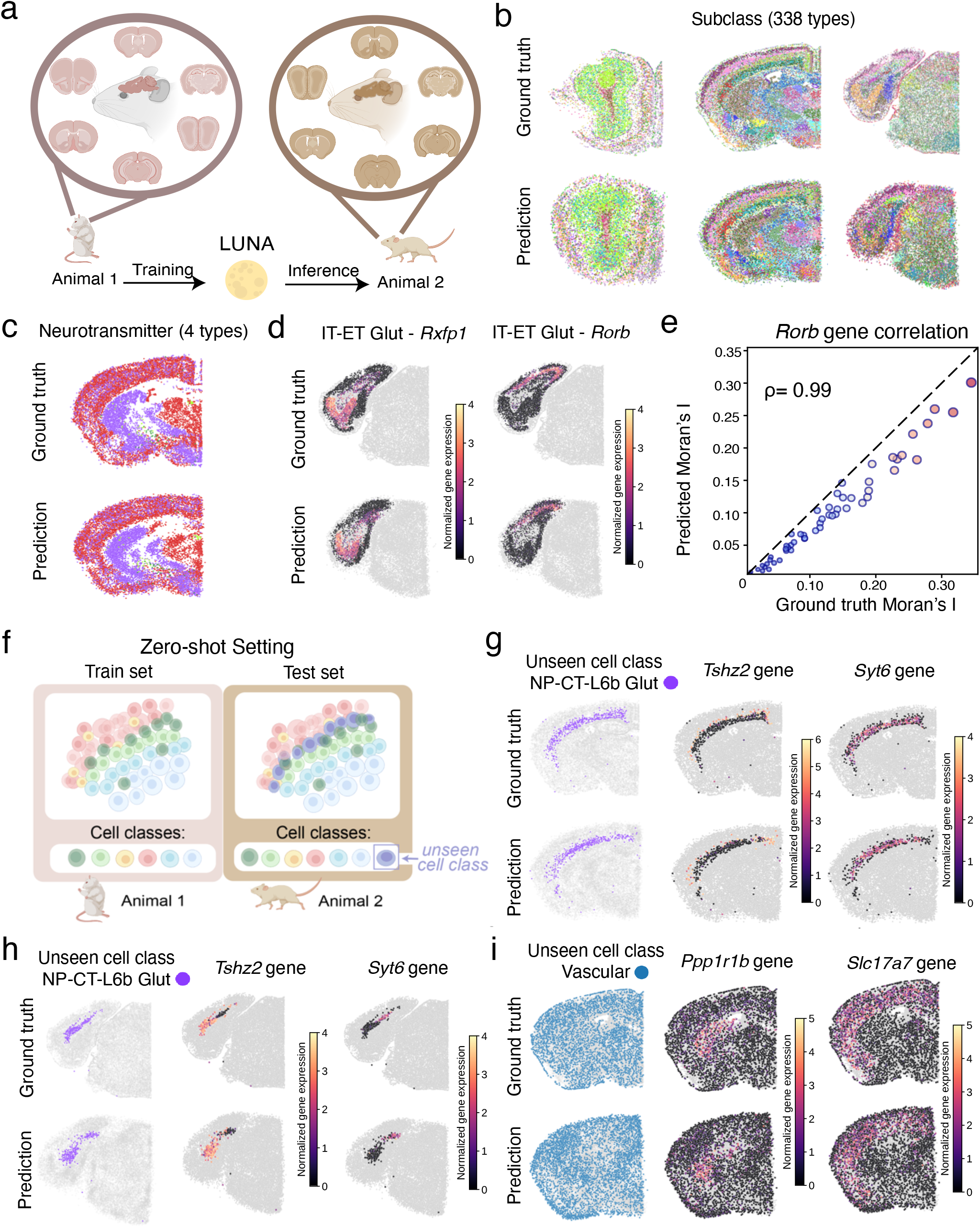
*(preceding page)*: LUNA accurately reconstructs the mouse brain atlas of 1.23 million cells. **(a)** LUNA was trained on all slices from the Animal 1 in the ABC atlas (2.85 million cells) and then applied to predict the tissue structure of cells from the Animal 2 unseen during training (1.23 million cells). **(b)** Examples of tissue reconstruction results using LUNA on three representative slices from distinct major brain regions of the MERFISH whole mouse Allen Brain Cell (ABC) atlas [27], comprising 12, 610 cells, 30, 931 cells and 22, 010 cells, respectively. For visualization purposes, three slices were randomly selected from the ABC atlas. Cells are coloured according to their cell type annotations with 338 distinct cell subclasses, showing ground truth (top) and predicted (bottom) locations. **(c)** LUNA’s predictions across 4 neurotransmitter types. Ground truth locations (top) and LUNA’s predictions (bottom) for a slice with 35, 738 cells . **(d)** Spatial expression patterns of two spatially variable genes in the IT-ET Glut cell type. To select spatially variable genes, we calculated Moran’s I values for all genes (1, 122 in total) based on ground truth locations and identified those with the highest Moran’s I value. Representative genes include *Rxfp1* and *Rorb* for IT-ET Glut cells. Cells from other cell types are shown in gray color. **(e)** Spatial autocorrelation for the *Rorb* gene computed using the Moran’s I on the ground-truth locations and the locations predicted by LUNA. Each point represents a single slice, with a total of 66 slices. The slices are color-coded from blue (low) to red (high) based on the true Moran’s I values for *Rorb*. Points near the diagonal indicate better spatial pattern preservation by LUNA. Pearson’s correlation coefficient (*ρ*) is displayed in the top left. **(f)** Zero-shot setting evaluation of LUNA. We exclude all cells from a randomly chosen cell class (out of 34 total classes) during model training and evaluate LUNA’s predictions on the cells from a held-out cell class, without any further training or refinement of the model. **(g, h)** LUNA effectively generalizes to unseen cells from NP-CT-L6b Glut cell class in the zero-shot setting. NP-CT-L6b Glut cells (*n* = 69, 641 in Animal 1) were excluded during training. The model was evaluated on Animal 2, including NP-CT-L6b Glut cells (*n* = 29, 262 in Animal 2). The spatial distribution of the excluded class (left), and the spatial expression patterns of the genes *Tshz2* and *Syt6* (middle and right) in the NP-CT-L6b Glut cell type across two representative slices (g, h). Genes *Tshz2* and *Syt6* were identified as the most spatially variable among 1, 122 genes, selected based on the highest Moran’s I values calculated from ground-truth cell locations. **(i)** Zero-shot generalization to unseen cells from the vascular cell class. Vascular cells (*n* = 447, 941 in Animal 1) were excluded during training. The model was evaluated on Animal 2, including vascular cells (*n* = 198, 550 in Animal 2). The spatial distribution of the excluded class (left), and the spatial expression patterns of the genes *Ppp1r1b* and *Slc17a7* (middle and right) in vascular cells for a representative slice. Icons in **(a)** and **(f)** were created using BioRender.

We evaluated LUNA’s predictions using the ground-truth cell locations. We found that LUNA’s predictions agree well with the ground-truth cell locations across 11 major regions identified in the ABC atlas, despite their distinct structural characteristics (Supplementary Fig. 1-3). For example, LUNA accurately captured the circular structure of the olfactory bulb region, the layered organization of the isocortex region, and the anatomical separation between the brain stem and cerebrum (Fig. 2b). In the slice from the olfactory bulb region affected by sequencing artifacts, LUNA inferred the spatial distribution of cells in areas lacking ground truth spatial information.

To further evaluate the generation results, we compared LUNA’s predictions across various levels of categorization, ranging from neurotransmitter level (4 types) to subclass level (338 types), and cell supertype level (1, 201 types). LUNA accurately positioned cells at different levels of categorization, from coarser to finer (Fig. 2c and Supplementary Fig. 4).

We next assessed LUNA’s ability to correctly position cells within a cell type based on their gene expression profiles. We identified spatially variable genes by computing Moran’s I statistic [30] and visualized the expression values of genes with the highest Moran’s I values (Supplementary Note 3). LUNA successfully captured the spatial expression patterns of genes within individual cell types, demonstrating its ability to preserve biological information. For example, LUNA accurately positioned cells expressing *Rxfp1* and *Rorb* in extratelencephalic and intratelencephalic neurons containing the neurotransmitter Glut (Fig. 2d). We also observed a high reconstruction quality across non-neuronal cells (Supplementary Fig. 5). For example, LUNA correctly positions the physical locations of a small group of 85 cells out of 3, 725 astrocyte-ependymal cells expressing the *Cfap206* gene (Extended Data Fig. 2).

To quantitatively evaluate the conservation of spatial gene expression patterns, we computed spatial autocorrelation using the Moran’s I statistic for all 1, 122 genes across the 66 slices. We compared Moran’s I values derived from LUNA’s predicted locations with those from the ground-truth cell locations. This analysis revealed a strong correlation between the Moran’s I values based on LUNA’s predictions and those based on the true locations, with an average Pearson correlation coefficient of 0.95 across all genes (Supplementary Fig. 6). For instance, the spatial autocorrelation for the spatially variable *Rorb* gene was consistently well-preserved across all slices, with a near perfect Pearson’s correlation coefficient (Fig. 2e). These results demonstrate that LUNA effectively preserves spatial gene expression patterns throughout the mouse brain.

We further validated LUNA’s capability to predict spatial information of missing slices. We trained LUNA on a subset of spatially measured slices from the ABC dataset and evaluated on 41 randomly held-out slices. LUNA’s predictions agree well with ground-truth cell locations on this cross-slice generalization task (Supplementary Fig. 7-8).

### LUNA generalizes to unseen cell classes

We next evaluated the capability of LUNA to predict the spatial locations of cells from classes unseen during training without any additional training or fine-tuning of the model (Fig. 2f). We refer to it a zero-shot capability, since the model was never trained on any samples from the unseen cell class [31, 32]. We excluded each of the 31 broad cell class categories that have sufficient number of cells (more than 1000 cells across the entire atlas) identified in [27] and then trained LUNA on slices from the Animal 1 in the ABC atlas without the held-out cell class (Supplementary Note 1). We then applied LUNA to predict tissue structures for all cell classes in Animal 2, including the held-out cell class. Across different cell classes, LUNA correctly positioned cells within the spatial architecture of the tissue (Extended Data Fig. 3-5). For example, LUNA correctly positioned NP-CT-L6b Glut cells in the isocortex, cortical subplate and hippocampal formation regions (Fig. 2g, h). We additionally examined the spatial expression patterns of spatially variable genes in NP-CT-L6b Glut cells. We identified genes *Tshz2* and *Syt6* as genes with the highest Moran’s I values among 1, 122 genes based on the ground-truth cell locations. We found that within the unseen NP-CT-L6b Glut class, LUNA correctly positioned cells according to the *Tshz2* and *Syt6* expression patterns (Fig. 2g,h). To analyze the similarity of the unseen NP-CT-L6b Glut cells with seen classes, we examined the gene expression similarity between this class and other seen classes. NP-CT-L6b Glut cells partially share gene expression patterns with several seen classes (Supplementary Fig. 9). This suggests that LUNA captures shared spatial gene expression patterns and models continuous spatial variation in gene expression, enabling it to infer spatial coordinates for previously unseen cell types. We further analyzed other randomly excluded cell classes, such as the vascular cell class that is spread across the entire slice (Fig. 2i, Supplementary Fig. 10) and the HY GABA cell class (Supplementary Fig. 11), confirming that LUNA preserves key spatial gene expression patterns of unseen cell types, demonstrating strong zero-shot capabilities.

### Interpreting gene contributions to spatial structure prediction

LUNA can enable identification of genes whose exclusion affects the prediction of spatial structure the most. Specifically, we measured the sensitivity of spatial predictions to exclusions of each individual gene by masking its expression to zero during inference, and quantified the resulting changes in predicted cell positions. We computed discrepancies between spatial predictions obtained with the full gene panel and those obtained with one gene excluded (Fig. 3a).

**Figure 3.**
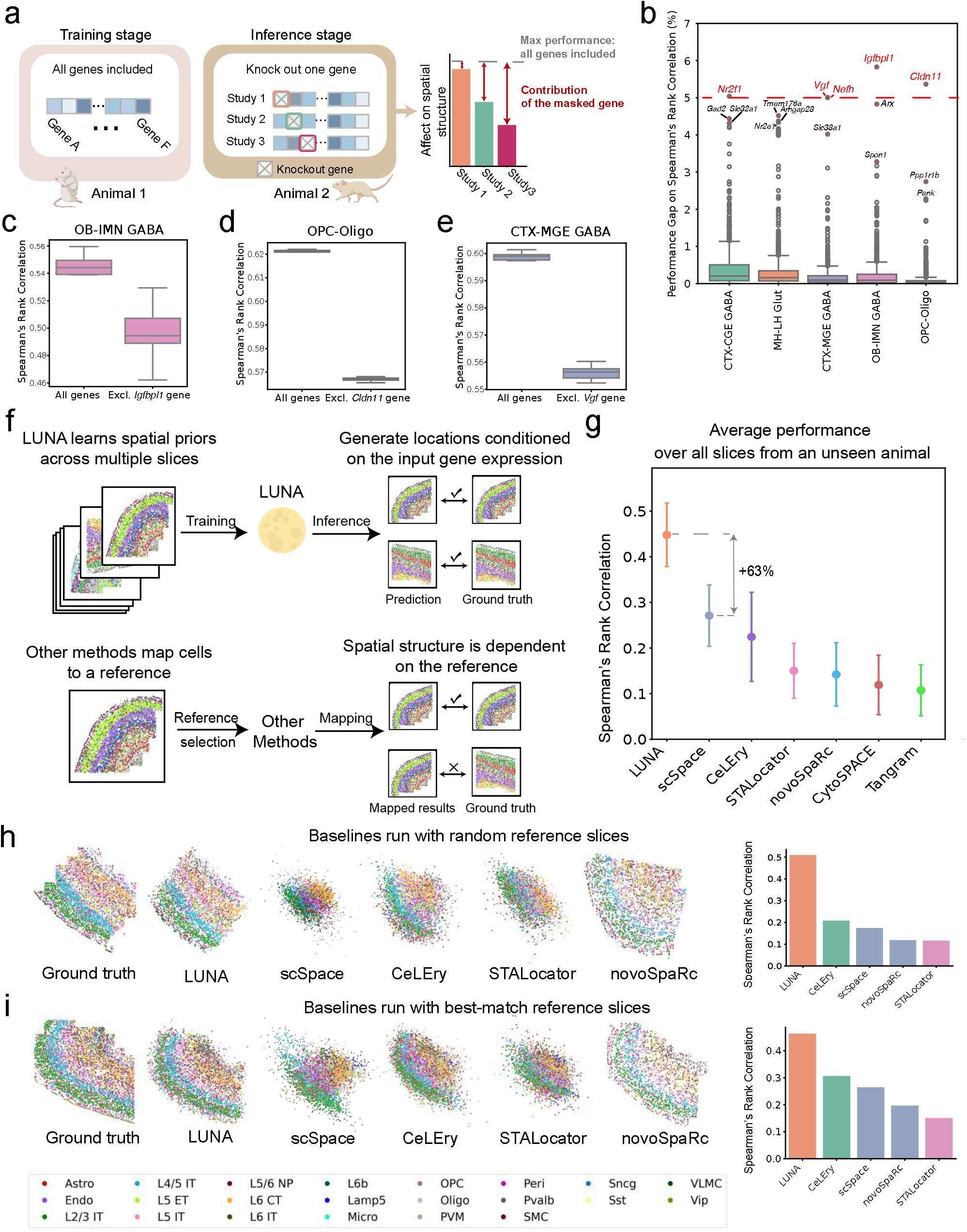
*(preceding page)*: LUNA identifies genes that have a strong impact on predicting spatial organization and outperforms alternative baselines on predicting tissue structure. **(a)** We train LUNA on all slices from the Animal 1 in the ABC atlas (2.85 million cells) with the whole gene panel. During inference, we mask expressions of individual genes, repeating this process for each gene in the panel (1, 122 genes). We then assess the contribution of individual genes for predicting spatial organization of Animal 2 with LUNA. To quantify the contribution of the masked gene, we compare the difference in the physical spaces between the full-gene model and the gene-excluded model. **(b)** Performance difference caused by excluding each gene for different cell classes. The performance is measured using the Spearman’s Rank Correlation (SRC) coefficient computed within each cell class. Each box summarizes *n* = 1, 122 single-gene exclusion experiments (one per gene) for one cell class; box plots show the median (center), 25th–75th percentiles (box bounds), 1.5*×* the interquartile range (whiskers) and outliers. The five cell classes were selected as the classes whose spatial structure is most affected by excluding specific genes across all cell classes. Genes causing more than a 5% performance change are marked in red. **(c-e)** The effect of different initializations of the initial noise during inference. We conduct 10 experiments with different random seeds, focusing on genes that significantly influence specific cell classes. We compare SRC when using all genes with scenarios excluding **(c)** *Igfbpl1* in OB-IMN GABA, **(d)** *Cldn11* in OPC-Oligo, and **(e)***Vgf* in CTX-MGE GABA. Box plots show the median (center), 25th–75th percentiles (box bounds), and whiskers extend to the most extreme value within 1.5*×* the IQR of the box edges. **(f)** LUNA fundamentally differs from the existing methods. LUNA takes as input different slices and learns spatial priors across all slices during training. In the inference stage, LUNA generates cell locations conditioned on their gene expressions by leveraging learnt spatial priors. In contrast, other methods require the pre-selection of a single reference slice and then map cells to the reference slice. Consequently, the spatial structure of their results heavily depends on the chosen reference slice. **(g)** Performance comparison on the cross-animal generalization task between LUNA and alternative baselines on the MERFISH mouse primary motor cortex atlas [43]. The performance is measured as the median Spearman’s rank correlation between the ground truth spatial coordinates and the predicted spatial coordinates over all slices from an unseen mice (31 slices, 118, 036 cells). Higher correlation indicate better prediction performance. Error bars represent standard deviation across 31 slices. **(h)** Visualization of ground-truth locations and predictions using LUNA and alternative baselines for one example slice (5, 235 cells) using randomly selected reference slice. Colors denote cell class labels. Performance comparison on that slice using Spearman’s rank correlation between the ground truth spatial coordinates and the predicted spatial coordinates as a metric (far right). **(i)** Visualization of ground-truth locations and predictions using LUNA and alternative baselines for one example slice (5, 024 cells) using the best matched slice as a reference for alternative baselines. The best matched slice was found using prior information about distribution of cell classes which LUNA does not use. Colors denote cell class labels. Performance comparison on that slice using Spearman’s rank correlation between the ground truth spatial coordinates and the predicted spatial coordinates as a metric (far right). Icons in **(a)** were created using BioRender.

We investigated how individual genes influence the predicted spatial locations of major cell classes in the ABC atlas. We trained LUNA on all slices from Animal 1 with the whole gene panel and, during inference, masked individual genes from Animal 2, quantifying each gene’s cell-class-specific contribution as the Spearman rank correlation (SRC) within each class (Fig. 3b, Extended Data Fig. 6, Supplementary Table 1). Certain genes had a particularly strong effect on prediction accuracy, including *Igfbpl1, Cldn11, Vgf, Nefh*, and *Nr2f1* (Fig. 3b), with the highest impact in the olfactory bulb–intracerebral migrating stream GABA neurons (OB-IMN GABA), oligodendrocyte precursor cells and oligodendrocytes (OPC-Oligo), cortex–medial ganglionic eminence GABA neurons (CTX-MGE GABA), and cortex–caudal ganglionic eminence GABA neurons (CTX-CGE GABA), respectively. Functionally, these genes are closely tied to cell identity and function: *Igf-bpl1* is involved in regulating cell growth and GABAergic transmission in mice [33, 34]; *Cldn11* encodes for oligodendrocyte-specific protein essential for myelin sheath formation [35] and previous work has shown that myelin mutant mice lacking expression of the *Cldn11* gene in oligodendrocytes exhibit central auditory deficits, reduced anxiety-like behavior and neurotransmitter imbalances [36]; *Vgf* and *Nefh* genes are biomarkers related to neurodegenerative diseases and neuronal damage [37, 38]; *Nr2f1* plays a crucial role in neurodevelopment, significantly influencing cortical development and network activity maturation [39, 40]. We further evaluated the performance variation with different initializations of the noise in the inference phase, confirming the robustness of our findings and systematic influence of *Igfbpl1, Cldn11, Vgf, Nefh* and *Nr2f1* genes on specific cell classes (Fig. 3c-e, Supplementary Fig. 12).

LUNA can also assess the joint contribution of gene pairs on spatial organization. Focusing on the 332 ligand-receptor pairs from OmniPath [41] that overlap the ABC atlas gene library, we jointly masked each pair during inference and measured its effect on the five cell classes most affected in the single-gene analysis (Extended Data Fig. 7). The strongest interactions involved vesicular neurotransmitter transporters: *Slc32a1* (a GABA transporter) in CTX-CGE GABA neurons, and *Slc17a7* (a glutamate transporter) in MH-LH Glut neurons, with *Slc17a7* also affecting CTX-CGE GABA neurons and oligodendrocytes. Interactions with the *Thbs4* ligand, whose loss causes a dystrophic phenotype in mice [42], most affected OB-IMN GABA neurons.

### LUNA outperforms other methods by a large margin

LUNA is a unique method in its capability to reconstruct complex tissue structures. Existing methods are reference mapping methods and require the pre-selection of a single reference slice to which then new cells are mapped based on the similarity of the gene expression between the cells in the reference slice and the input gene expression profiles (Fig. 3f). This mapping is performed either by explicitly learning a mapping matrix, as in novoSpaRc [20], Tangram [14], STEM [16], and CytoSPACE [15], or by implicitly regressing from gene expression space to the physical space, as in scSpace [17], STALocator [18], CeLEry [19] (Supplementary Note 2).

We compared the performance of LUNA to alternative methods on the spatially resolved cell atlas of the mouse primary motor cortex sequenced by MERFISH [43] (Supplementary Note 1). We used 31 slices (118, 036 cells) from one animal as the test set and trained LUNA on 33 slices (158, 379 cells) from another mouse to learn spatial priors across multiple slices. During inference, all methods were conditioned only on the gene expression. Since existing methods can not accept multiple slices as input, we first randomly selected a single slice from the training mouse to serve as the reference, repeating the procedure for each slice of the testing mouse.

We evaluated the results using the Spearman’s rank correlation between the predicted and ground truth locations, averaging the performance across all slices from the unseen mouse (Supplementary Note 3). LUNA achieved an average correlation coefficient of 0.448, outperforming the best alternative baseline scSpace by 63% (Fig. 3g). Performance gains of LUNA were retained using other evaluation metrics, such as precision and RSSD (Supplementary Note 3, Supplementary Fig. 13). CeLEry is the only other method besides LUNA capable of potentially leveraging information from multiple slices. We reran CeLEry using all slices from the training mouse as the training dataset. However, since CeLEry is a regression model that does not account for geometric constraints, it is not robust to variations in slice orientation or cutting shapes within the training set. Consequently, we observed a significant performance decline when CeLEry was trained with multiple slices compared to using a single slice (Supplementary Fig. 13). We visualized the spatial locations predicted by different methods on a sample slice from the unseen test mouse (Fig. 3h, Supplementary Fig. 14). LUNA accurately predicted the layer-wise structure of mouse primary motor cortex tissues and correctly inferred the spatial priors of the tissue, such as the contour of the tissue. We analyzed the spatial gene expression patterns of cell type specific marker genes. LUNA distinguished cells from different cell classes, accurately reflecting the spatial gene expression patterns (Supplementary Fig. 15). In contrast, other methods struggled due to the dissimilarity between the reference and test slices.

The limitations of other methods stem from their reliance on pre-selected reference slices, which constrains their generalization capabilities. To systematically evaluate the sensitivity of existing methods on the reference slice, we selected different slices as a reference training slice and evaluated performance on a fixed test slice. We computed the average performance of these methods across different reference slices. LUNA achieved a 121% higher Spearman’s rank correlation coefficient compared to the average performance of the best alternative method, scSpace, where the average is calculated over different reference slices (Supplementary Fig. 16). Given the sensitivity of reference mapping methods to the choice of reference slices, we further explored alternative strategies by selecting the most similar slice from another animal as the reference by using cell class information as a prior (Supplementary Note 2). We then reassessed the performance of alternative methods using this prior information. This prior information improved the performance of alternative baselines (Fig. 3i); however, they still lagged significantly behind LUNA’s performance. Even without using any prior information, LUNA outperformed all baselines, achieving 37.5% better performance on SRC than the best alternative method, CeLEry (Supplementary Fig. 17). We further compared LUNA to other alternative methods on the MERFISH ABC Atlas in the setting that selects the best reference slice for alternative methods as a prior (Supplementary Note 2). LUNA again significantly outperformed all alternative methods in all evaluation metrics even without using any prior information (Supplementary Fig. 18). Moreover, alternative methods fail to generalize to cell classes unseen in the reference section, whereas LUNA successfully reconstructs these patterns (Supplementary Note 2; Extended Data Fig. 8). This gap reflects a difference in inductive bias: many of the benchmarked approaches depend heavily on relationships learned from the reference cell types seen during training, with limited capacity to capture gene-level spatial localization or inter-gene co-localization. Their performance therefore tends to degrade on cell types whose transcriptional profiles are substantially underrepresented in, or absent from, the training data.

### Reconstruction of scRNA-seq atlas

We next applied LUNA for reconstruction of tissue structures of 1.08 million dissociated single cells across 13 coronal slices from the mouse central nervous system (CNS) scRNA-seq atlas [28] (Fig. 4a). We used the model trained on the ABC MERFISH mouse brain atlas to predict spatial locations for cells in the scRNA-seq mouse CNS atlas integrated with the MERFISH atlas using Harmony [44] (Supplementary Note 1). Since the scRNA-seq atlas lacks spatial information, we validated LUNA’s performance using estimated cell locations derived from the integration of the scRNA-seq atlas with the STARmap PLUS CNS spatial atlas [45] (Supplementary Note 1). In [45], the mouse CNS spatial atlas was created using the in situ sequencing STARmap PLUS, which was then integrated with the scRNA-seq data using Harmony [44]. The integrated dataset underwent careful quality control by human experts, ultimately providing estimated cell locations of the scRNA-seq atlas [45] (Fig. 4b). To validate LUNA’s predictions, we used these estimated cell locations obtained after the integration. Notably, LUNA never used spatial information from the STARmap PLUS atlas.

**Figure 4.**
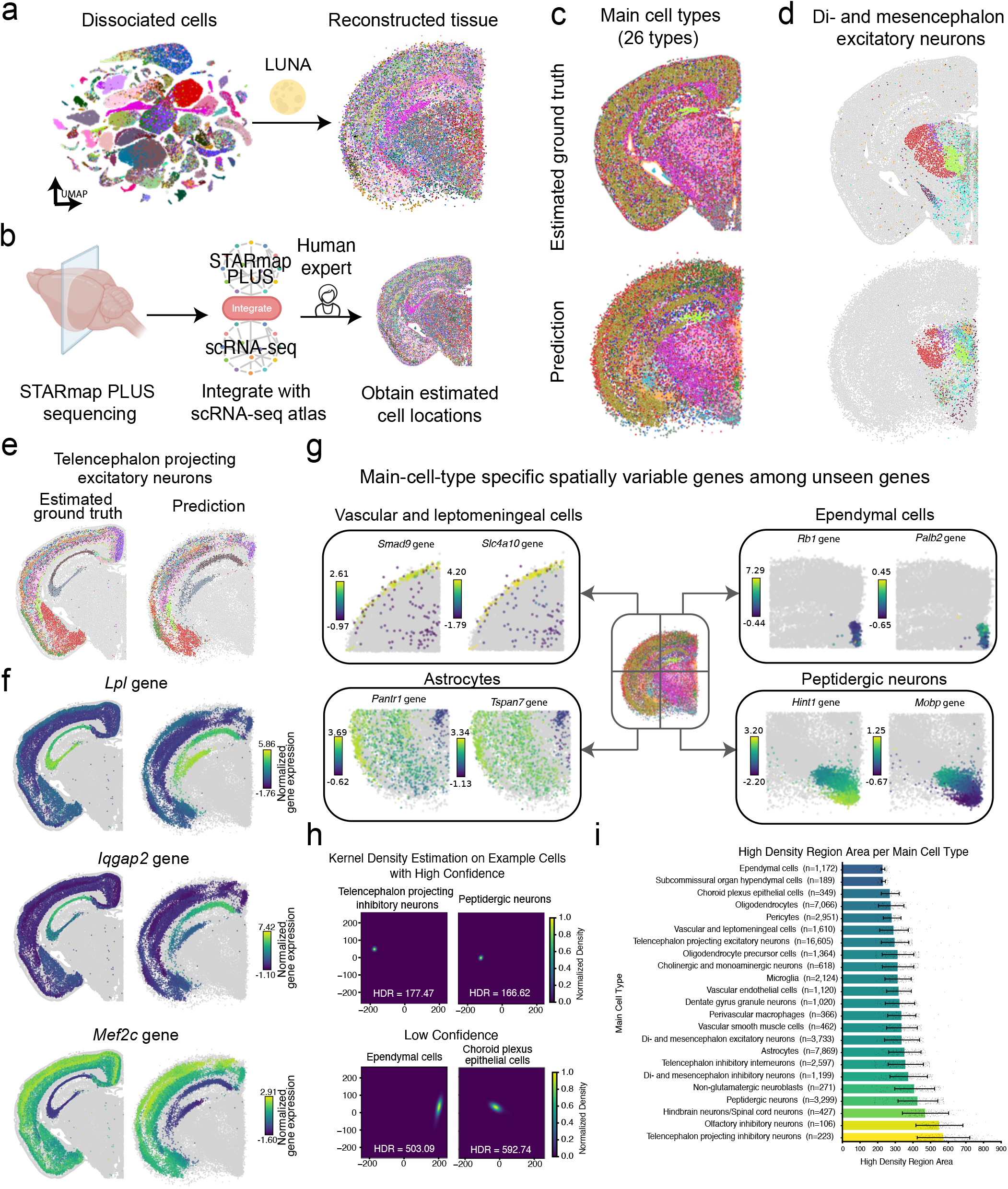
*(preceding page)*: LUNA predicts the tissue structure of 1.08 million cells from the scRNA-seq mouse central nervous system atlas. **(a)** Given the dissociated cells from the scRNA-seq atlas of the mouse central nervous system (CNS) [28], LUNA infers the tissue architecture. **(b)** Estimated ground truth spatial locations are obtained by sequencing mouse CNS using STARmap PLUS technology and integrating the CNS scRNA-seq dataset with the STARmap PLUS CNS dataset, followed by the manual evaluation of human experts. [45]. We use these estimated locations to validate LUNA’s predictions. **(c)** Tissue reconstruction of the scRNA-seq mouse central nervous system atlas using LUNA for one example slice (58, 117 cells). Cells are colored based on the main cell types (26 types). LUNA was trained on all the cells from Animal 1 of the ABC atlas (2.85 million cells) and applied to generate cell locations for a scRNA-seq atlas from the mouse central nervous system (1.08 million cells, 13 coronal slices). **(d-e)** Tissue reconstruction of the scRNA-seq mouse central nervous system atlas using LUNA for **(d)** di- and mesencephalon excitatory neurons and **(e)** telencephalon projecting excitatory neurons. The top plot shows estimated ground truth locations obtained by aligning scRNA-seq atlas with the STARmap atlas and the bottom plot shows LUNA’s predictions. Cells from each main cell type are colored by their sub-molecular types. Cells from other main cell types are shown in gray color. **(f)** Spatial expression patterns of three spatially variable genes across telencephalon-projecting excitatory neurons. To identify these genes, we calculated Moran’s I values for all 11, 844 genes based on LUNA’s predictions, selecting three genes with distinct spatial patterns from the top 10 genes with the highest Moran’s I values. The representative genes include *Lpl, Iqgap2*, and *Mef2c*. Cells from other main cell types are depicted in gray. **(g)** Main-cell-type specific spatially variable genes (SVGs) detected using LUNA’s predictions among genes that are unseen during training and unique to the scRNA-seq atlas. Four example main cell types are displayed, including both non-neuronal and neuronal cell types. Cells from other main cell types are depicted in gray. Genes are randomly selected among their top five SVGs with the highest Moran’s I values. **(h)** Kernel density estimation visualizations of example cells with high (top row) and low (bottom row) confidence predictions. We generate 150 samples with different random seeds and compute the area encompassing 80% of predictions, defining this as the high density region (HDR) value for each case cell. Smaller HDR region implies lower uncertainty of a cell’s location. Cells with high confidence display sharply peaked distributions (low HDR values), while low-confidence cells exhibit more dispersed distributions. **(i)** Average area of high-density regions computed from kernel density estimation of predicted spatial coordinates for each main cell type. Each bar represents a main cell type, with its height corresponding to the mean HDR area for cells of that type, and error bars indicating standard deviations. Smaller HDR areas reflect smaller uncertainty in predicting locations of cells. Number of cells (*n*) for each main cell type is annotated in the panel. We overlay maximum 200 randomly sampled points as the grey dots for each type. Icons in **(b)** were created using BioRender.

Applying LUNA to the scRNA-seq CNS atlas, its predictions aligned closely with cell locations estimated through the integration data with the STARmap PLUS atlas at both the main cell type level (26 types; Fig. 4c; Supplementary Fig. 19) and the finer sub-molecular type level, which includes 216 distinct types (Supplementary Fig. 20). The agreement is particularly pronounced in the isocortex, hippocampal formation, thalamus, and hypothalamus regions. We further evaluated LUNA’s performance in reconstructing mouse CNS tissue architecture across various main cell types (Supplementary Fig. 21). Additionally, we examined LUNA’s predictions for specific neuronal main cell types including di- and mesencephalon excitatory neurons (Fig. 4d), and telencephalon projecting excitatory neurons (Fig. 4e), along with their respective sub-molecular types. The results indicate that LUNA not only accurately placed cells across main cell types but also precisely predicted the spatial relationships of complex sub-molecular types within these neuronal cells. This underscores LUNA’s capability to capture both broad and intricate cellular architectures within the CNS.

To evaluate the conservation of spatial gene expression patterns, we analyzed the spatial distribution of three highly variable genes—*Lpl, Iqgap2*, and *Mef2c*, within telencephalon-projecting excitatory neurons (Fig. 4f). These genes were selected based on their high Moran’s I values, calculated from a dataset of 11, 844 genes using LUNA’s predictions. This analysis revealed that the spatial gene expression patterns predicted by LUNA closely align with the estimated ground truth, demonstrating LUNA’s effectiveness in capturing gene expression gradients at the cellular level. Specifically, *Iqgap2* gene shows a gradual downregulation across the Ammon’s horn region (CA2) of the hippocampus, while *Lpl* displays an opposite trend with gradual upregulation in this region. For *Mef2c* gene, expression is minimal in telencephalon-projecting excitatory neurons within the CA2 region but substantially upregulated in the isocortex and olfactory areas. Overall, these results confirm that LUNA effectively reconstructs the complex spatial architecture of the mouse CNS scRNA-seq atlas and captures gene expression spatial changes. This underscores LUNA’s applicability in reconstructing cell locations for scRNA-seq atlases. We evaluate these predictions quantitatively by computing spatial autocorrelation using Moran’s I statistics across all 11, 844 genes from the scRNA-seq CNS atlas and observe high agreement between LUNA’s predictions and estimated groundtruth locations (Supplementary Fig. 22). To further assess LUNA’s performance in the presence of batch effects, we evaluated LUNA on both non-integrated and integrated versions of the scRNA-seq data. LUNA maintains good performance despite batch effects, demonstrating robustness to technological differences. However, it still benefits from batch correction to further enhance overall performance. (Supplementary Fig. 23).

By predicting the locations of dissociated scRNA-seq data, LUNA can be used to detect spatially variable genes that are absent from the spatial dataset used for model training. To demonstrate this, we detected main-cell-type specific spatially variable genes among genes that are unseen during training (*i.e*., specific to scRNA-seq atlas) using LUNA’s predicted cell locations (Supplementary Note 5). Using predicted locations, we identified genes with distinct spatial patterns in both neuronal and non-neuronal cells (Fig. 4g). Additionally, we report the top five (among the total of 11, 844 genes) unseen genes with the highest Moran’s I value for different main cell types (Supplementary Table 2).

As a generative model, LUNA allows capturing multiple plausible spatial arrangements, enabling the estimation of uncertainty in predictions. To estimate uncertainty in locations of scRNA-seq atlas, we generated 150 samples from the LUNA model using different random seeds. We then applied kernel density estimation to the predicted locations of each cell to analyze spatial uncertainty (Supplementary Note 6). For each individual cell, LUNA can estimate high density regions (HDR) and thus model uncertainty in predictions (Fig. 4h). Extending this modeling on the level of main cell types, LUNA can be used to model uncertainty of main cell types and identification of regions in which locations of cells are interchangeable. For example, we found that telencephalon projecting inhibitory neurons on average have smaller HDR regions and thus lower uncertainty in the spatial locations of cells belonging to these main cell types, while ependymal cells have the highest uncertainty (Fig. 4i).

### LUNA infers locations of spatially unmapped nuclei in the Slide-tags data

LUNA can be used to predict the tissue locations of nuclei lost during cell profiling with Slide-tags technology [29]. Slide-tags enables profiling single-cell and spatially resolved transcriptome by tagging nuclei with spatial barcode oligonucleotides derived from DNA-barcoded beads with known positions and then using tagged nuclei as an input to single-nucleus profiling assays [29]. However, many nuclei are lost during the barcoding process due to a combination of dissociation and microfluidic losses. To compensate for the sparsity in Slide-tags data, we applied LUNA to assign spatial locations to the spatially unmapped nuclei in the Slide-tags data.

We first applied LUNA to the sagittal section of the embryonic mouse brain at embryonic day 14 (E14) dataset consisting of 4, 623 spatially mapped nuclei with Slide-tags (Fig. 5a) (Supplementary Note 1). To evaluate LUNA’s performance using ground truth nuclei locations, we used 4, 160 spatially mapped nuclei as the train set and evaluated LUNA’s performance on remaining 463 cells. LUNA’s predictions agree well with the ground truth locations of cells obtained with Slide-tags (Fig. 5b). Without using any cell class information, LUNA accurately predicted cell locations, even for different neuronal subclasses. We further validated these predictions by computing spatial autocorrelation using Moran’s I index across all genes used for training (Fig. 5c). The relationship exhibits an almost perfect correlation, with a slope slightly below one (slope=0.864), indicating a modest systematic underestimation of spatial autocorrelation values alongside a near-perfect rank agreement.

Encouraged by these results, we applied LUNA to infer locations of spatially unmapped nuclei due to nuclei losses with Slide-tags. We trained LUNA on spatially mapped cells of mouse E14 tissue and applied the model to infer locations of 4, 414 spatially unmapped nuclei (Fig. 5d). To validate predictions of LUNA on the cell class level, we ran linear support vector classifier to infer cell classes of spatially unmapped nuclei since annotations for spatially unmapped nuclei were missing in the original dataset (Supplementary Note 1). We found that LUNA placed spatially unmapped nuclei in proximity of cells with the same class, enriching sparse Slide-tags dataset. We further examined the spatial distribution of specific cell types, additionally confirming that LUNA placed nuclei from both neuronal and non-neuronal cells to the correct spatial regions (Fig. 5e, Supplementary Fig. 24).

**Figure 5.**
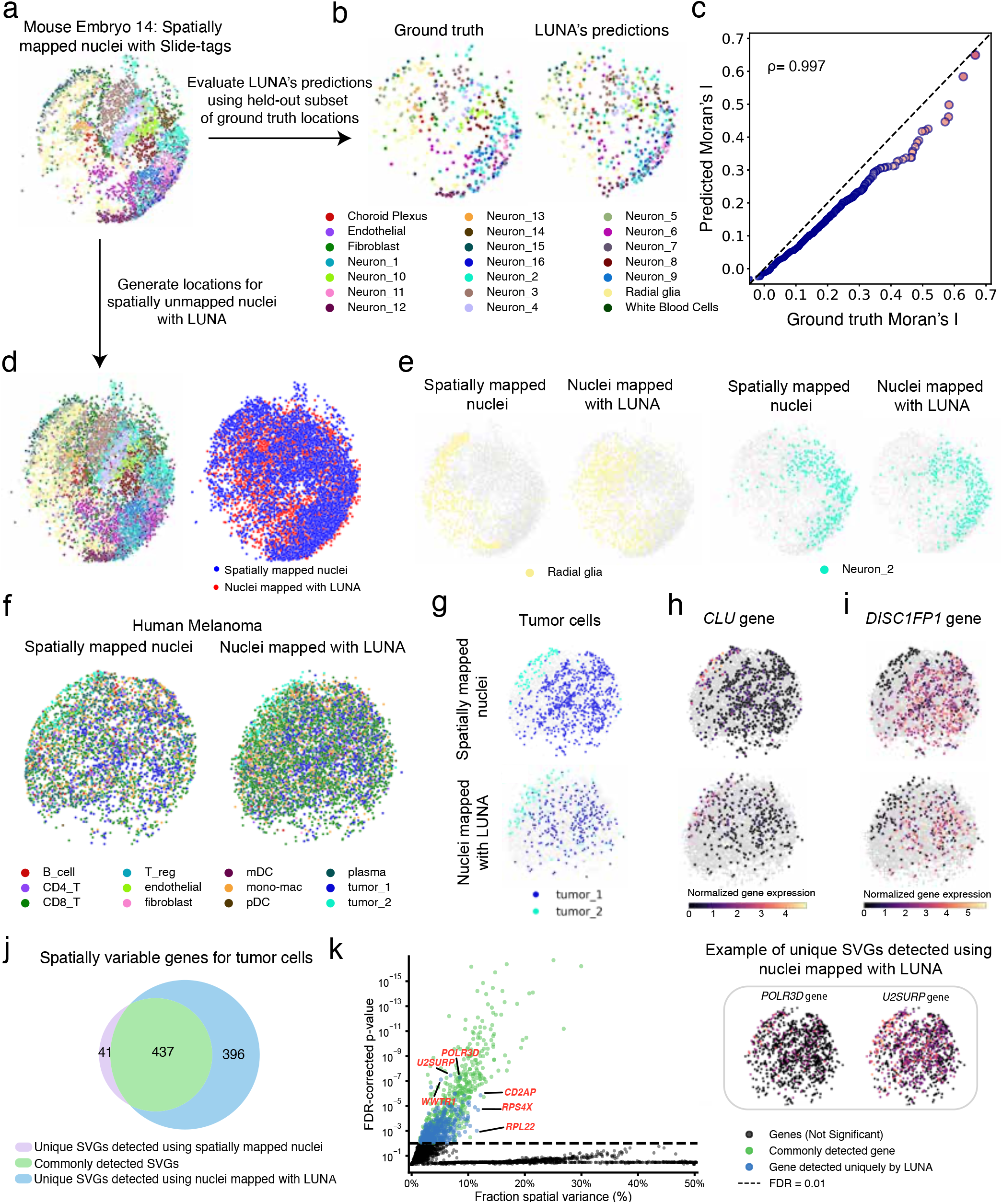
*(preceding page)*: LUNA infers locations of spatially unmapped nuclei in the Slide-tags data. **(a)** Spatially mapped nuclei using Slide-tags from the embryonic mouse brain at embryonic day 14 (E14) containing 4, 623 cells [29]. Cells are coloured according to their cell type annotation. **(b)** LUNA’s predictions of nuclei locations agree well with the ground truth locations from Slide-tags. Ground truth locations (left) and LUNA’s predictions (right) on the subset of the held-out dataset of the mouse E14 embryonic brain. LUNA was trained on 4, 160 cells with ground truth locations from spatially mapped nuclei and we evaluated its predictions on the held-test set of 463 cells. **(c)** Spatial autocorrelation computed using the Moran’s I on the ground truth locations and locations predicted by LUNA on the held-out test set of mouse E14 embryonic brain. Each point represents a gene and 690 highly differentially expressed genes are visualized. Genes are colored on a gradient from blue (low) to red (high) based on their ground truth Moran’s I value. Points closer to the diagonal indicate better preservation of spatial patterns by LUNA. The Pearson’s correlation coefficient (*ρ*) is reported on the top left. **(d)** LUNA generated locations of 4, 414 initially unmapped nuclei of the mouse E14 embryonic brain. LUNA was trained on spatially mapped nuclei and then applied to infer locations for the unmapped nuclei. The combined locations of nuclei mapped by LUNA and those spatially mapped by Slide-tags with cells coloured according to their cell type annotation (left). Cells colored based on the source of cell locations (right). **(e)** Nuclei mapped by Slide-tags (left) and nuclei mapped by LUNA for Neuron 2 (cyan color) and Radial glia (yellow color) cell types. All other cells are shown in gray color. **(f)** LUNA enriched the Slide-tags human metastatic melanoma sample from 4, 804 cells to 6, 466 cells by generating locations for spatially unmapped nuclei. The left panel shows the ground truth locations of spatially mapped nuclei used as the training dataset, while the right panel combines these with the nuclei locations predicted by LUNA. **(g)** LUNA correctly inferred the spatial compartment of two tumor clusters segregated into spatially distinct compartments. Spatially mapped nuclei by Slide-tags (top) and locations of unmapped nuclei generated by LUNA (bottom). Tumor cells are colored according to their type, while other cells are shown in gray color. **(h, i)** Spatial expression patterns of **(h)** the *CLU* gene and **(i)** the *DISC1FP1* gene across spatial compartments of tumor cells. We selected these genes as spatially variable genes (SVGs) for each tumor type. Spatially mapped nuclei by Slide-tags (top) and locations of unmapped nuclei generated by LUNA (bottom). Tumor cells are colored according to expression levels of these genes. Other cells are shown in gray color. **(j)** Number of SVGs uniquely detected by LUNA. SVGs were identified with SpatialDE [46] (likelihood-ratio test, one-sided) in tumor cells using spatially mapped nuclei (*n* = 899) and nuclei mapped with LUNA (*n* = 1,250). FDR-corrected [47] *p*-value *<* 0.01. **(k)** Fraction of variance explained by spatial variation (FSV) versus significance of spatial variation (FDR-corrected *p*-value) for SVGs. Each dot represents a gene (*n* = 9,413). The dashed horizontal line indicates the significance cutoff (FDR *<* 0.01). Genes are categorized as non-significant, significant genes commonly detected, and significant genes uniquely detected using nuclei mapped with LUNA. Among genes detected uniquely by LUNA, the top 3 genes that have the lowest FDR-corrected *p*-value (*POLR3D, U2SURP* and *WWTR1* genes) and the top 3 genes that have the highest FSV (*CD2AP, RPL22* and *RPS4X* genes) are highlighted in red. The spatial gene expression patterns of *POLR3D* and *U2SURP* genes are visualized (top right).

We next applied LUNA to a human metastatic melanoma sample obtained with Slide-tags technology [29]. This sample includes a diverse array of cells, including immune cells and tumor cells. We trained LUNA on 4, 804 spatially mapped nuclei and applied it to generate locations for the spatially unmapped nuclei. We again transferred cell type annotations of spatially unmapped nuclei with the linear support vector classifier to validate LUNA’s predictions (Supplementary Note 1). We found that LUNA successfully enriched the dataset to the total of 6, 466 spatially mapped cells (Fig. 5f). LUNA correctly placed two tumor subpopulations into spatially distinct compartments according to predicted cell class annotations (Fig. 5g), as well as non-tumor cells (Supplementary Fig. 25). To further validate LUNA’s predictions, we investigated the heterogeneity of these two tumor populations by identifying highly differential genes between the two cell classes. Cells expressing *CLU* gene are placed by LUNA in different spatial compartments than cells expressing *DISC1FP1* gene, which is also the case in the original dataset of spatially mapped nuclei of the melanoma sample (Fig. 5h, i).

We investigated whether LUNA-enriched Slide-tags data can increase the power to detect spatially variable genes in the human metastatic melanoma sample. We detected SVGs using SpatialDE [46], focusing exclusively on tumor cells in the sample (Supplementary Note 5). The LUNA-enriched dataset revealed 396 additional SVGs (FDR-corrected [47] p-value *<* 0.01) not detectable in the original sample, an 83% increase (Fig. 5j). Gene Ontology enrichment (Supplementary Fig. 26) was strongest for “Myc Targets V1” (MSigDB [48]) which includes genes regulated by the MYC oncogene known to promote cell proliferation and closely linked to tumor aggressiveness and survival outcomes [49], and the Ribosome pathway (KEGG [50]), with known causal associations with increasing cancer risk [51].

Among genes uniquely detected using LUNA-enriched sample, the genes with the lowest *p*-value include *U2SURP* (*p*-value = 3.4*e −* 8), *POLR3D* (*p*-value = 5.2*e −* 8), and *WWTR1* (*p*-value = 7.6*e −* 8) (Fig. 5k), whose abnormal activity is known to be associated with cancer [52–55]. We use the Fraction of Spatial Variance (FSV) [46] to measure the extent to which spatial patterns contribute to the total variance observed in gene expression data. Of all significant genes detected using the LUNA-enriched sample, 50 genes exceeded the average (FSV = 7.3%, over 833 identified SVGs), including *CD2AP* (FSV=12.2%), *RPL22* (FSV=11.7%) and *RPS4X* (FSV=11.4%) genes. *RPL22* is known to act as a tumor suppressor gene [56], while previous studies have suggested that *CD2AP* and *RPS4X* are associated with certain malignant tumors [57– 60]. We applied LUNA to enrich other Slide-tags datasets, including samples from human tonsil (Supplementary Fig. 27) and human cortex (Supplementary Fig. 28).

### LUNA generalizes to pathological conditions

To study the ability of LUNA to predict spatial locations of cells in tissues with undergoing pathological processes, we generated Xenium data from an *α*-synuclein-induced Parkinson’s disease (PD) mouse model [61, 62] (Methods). Mice were unilaterally injected with 2*µl* of pre-formed fibrils (PFFs) into the dorsal striatum at eight weeks of age (Fig. 6a). In this model, misfolded proteins propagate to neighboring neurons, seeding further aggregation in anatomically connected regions. Three months post-injection (3MPI), brains were sectioned and profiled. The same sections were subsequently immunostained to detect misfolded *α*-synuclein, providing an independent experimental validation of the predicted pathological changes (Fig. 6b).

**Figure 6.**
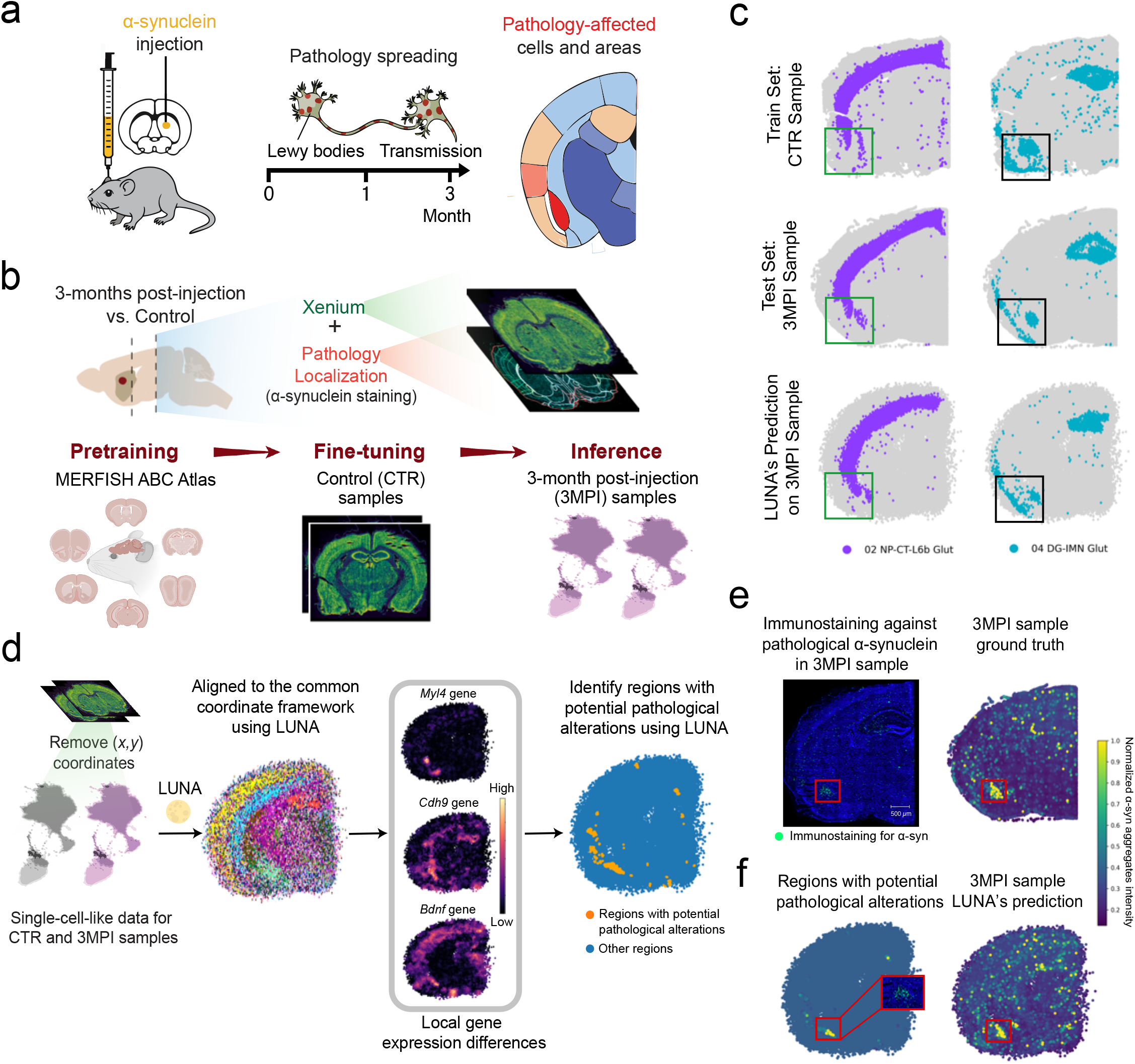
*(preceding page)*: LUNA enables robust spatial reconstruction of diseased samples using a model fine-tuned on wild-type control tissues. **(a)** Illustration of the *α*-synuclein (*α*-syn) pre-formed fibrils (PFF) injection model in the Parkinson’s disease mouse model. Misfolded *α*-syn is transmitted to neighboring neurons, seeding further aggregation and pathology spread in connected regions. To model this pathology, wild-type (WT) mice were unilaterally injected with 2 *µ*l of pre-formed fibrils into the dorsal striatum. This schematic was adapted from ref. [61]. **(b)** Illustration of the experiment. Brains from control (CTR) mice and from mice at three months post-injection (3MPI) were extracted, sectioned, and processed using Xenium spatial transcriptomics. For each brain, two distinct anatomical positions were collected and analyzed. The same sections from 3MPI samples were subsequently subjected to immunostaining for misfolded *α*-syn, providing validation of pathology-affected regions. We use the LUNA model pre-trained with MERFISH ABC Atlas, and fine-tune it on CTR samples, and then apply the model to infer spatial structures of 3MPI samples. **(c)** Spatial distributions of neuronal subtypes for the posterior section. Shown are NP-CT-L6b glutamatergic neurons (02 NP-CT-L6b, purple) and DG-IMN glutamatergic neurons (04 DG-IMN Glut, blue) in a training sample (CTR, top), a 3MPI sample (middle), and the LUNA-predicted spatial coordinates for the 3MPI sample (bottom). Gray points denote all the other cells for reference. **(d)** Cells exhibiting differential transcriptomic profiles between CTR and 3MPI samples identified from LUNA’s predictions. We removed spatial coordinates from the Xenium dataset and input single-cell-like gene expression data from the CTR and 3MPI sections into LUNA. During inference, LUNA aligns these samples to a common coordinate framework. We visualized the genes *Myl4, Cdh9*, and *Bdnf*, which were selected for their local spatial differential expression in the amygdala and hippocampus regions. Cells were colored by their local expression differences, with lighter shades indicating greater local shifts. To identify regions with potential pathological alterations, we further computed a unified score across genes with high distributional shifts between the 3MPI and CTR samples. Cells with significant local transcriptomic differences were labeled in yellow, while all others were shown in blue. These highlighted cells indicate regions with localized spatial transcriptomic alterations, potentially associated with pathological changes. **(e)** Immunostaining for *α*-syn pathology in the 3MPI sample (left) shows *α*-syn in green, with each dot representing an individual protein aggregate. Scale bar is defined on the panel. Ground truth cell locations for the same sample are shown on the right, with each cell colored by the intensity of *α*-syn fibril staining. Immunostaining in (e) was performed on two 3MPI sections with similar results. **(f)** LUNA enables spatial data-free identification of brain regions with spatial gene expression disregulations that correlate with *α*-syn pathology. Cells that are identified with potential pathological alternations between 3MPI and CTR (left) and all cells (right) are positioned according to LUNA’s predicted locations and colored by *α*-syn fibril staining intensity. A close-up view highlights the immunostaining of the regions identified by LUNA that potentially associate with pathological alterations, overlaid above the left plot. Icons in **(b)** were created using BioRender.

We used a LUNA model pretrained on the ABC MERFISH Atlas and we fine-tuned it on generated Xenium wild-type sections. We then applied the model to infer the spatial locations of diseased 3MPI sections that the model has never seen (Fig. 6b, Supplementary Note 1). LUNA successfully reconstructed the spatial organization of the different coronal sections (Supplementary Fig. 30). We visualized individual cell classes and observed strong alignment between LUNA’s predictions and the ground truth spatial distributions in the 3MPI section (Supplementary Fig. 31). Notably, in the amygdala region, LUNA’s predictions more closely resembled the spatial organization of the diseased samples than the wild-type samples on which the model was fine-tuned (Fig. 6c). We further quantitatively measure the spatial distribution similarities using Wasserstein distance. Across all cell classes, the distance between the predictions and the test section is consistently lower than that between the predictions and the training section, indicating that the model predictions better reflect the spatial layout of the diseased samples across all cell classes than that of the wild-type samples (Extended Data Fig. 9).

Encouraged by these results, we decided to test whether LUNA can be used to identify spatial gene expression deregulation, such as those we expect to be induced by *α*-synuclein pathology spread. Specifically, we reasoned that the physical space learned by LUNA can be used for cell alignment and the identification of regions of local gene-expression changes, without ever measuring spatial data. To achieve this, we first run LUNA on slices from both wild-type and 3MPI samples, resulting in spatial predictions in a shared coordinate framework (Fig. 6d, Supplementary Fig. 32). Then, we use these cell coordinates to quantify 3MPI-vs-wild-type local gene expression changes (Supplementary Note 7). We found differentially expressed genes like *Myl4, Cdh9* and *Bdnf* in the amygdala, and the hippocampus, some of which have been associated with neuronal stress or senescence [63–65]. We reasoned that the regions where cumulative changes across all genes were maximal were also the most likely to be the most affected by pathology. To test this, we computed a unified score for all cells based on their local gene expression differences between the 3MPI and wild-type samples (Supplementary Fig. 33, Supplementary Note 7). We computed Jensen-Shannon Divergence across all genes to identify those with the most significant distributional shifts between wild-type and 3MPI samples. Local gene expression differences were then assessed by comparing spatially averaged expression across each cell’s neighborhood. We found certain cluster of cells that are consistently flagged across more than half of the significantly divergent genes, revealing regions with potential pathological transcriptomic alterations (Fig. 6d, Supplementary Fig. 32).

To validate these regions of interest, we performed immunostaining against pathological *α*-synuclein in the 3MPI sample (Fig. 6e). We found that cells identified as locally significantly different by LUNA exhibited higher levels of *α*-synuclein aggregates (Fig. 6f, Supplementary Fig. 34). Interestingly, LUNA accurately predicted the locations of clusters of cells with pathological *α*-synuclein. These results showcase how LUNA can generalize beyond wild-type conditions, and can be applied to different biological conditions providing an unique opportunity to identify spatial gene expression dysregulations without spatial data.

We further show that LUNA is applicable without additional fine-tuning on healthy Xenium samples. Applying the library-size normalization and log-transformation described in Supplementary Note 1 (Methods) to both datasets, we train the model using only the 133 intersecting genes between the MERFISH ABC Atlas and Xenium datasets and directly evaluated the Xenium pathological samples without any fine-tuning on the healthy Xenium samples. The model achieves a median Spearman’s rank correlation of 0.673, and over 80% of cells achieve a Spearman’s rank correlation above 0.5, indicating robust performance (Extended Data Fig. 10a). We further observe robust performance at the cell class level and in localizing alpha-synuclein fibril staining intensity (Extended Data Fig. 10b-c). However, fine-tuning is nonetheless beneficial and yields further performance gains(Extended Data Fig. 10).

## Discussion

LUNA is a generative AI model that reconstructs tissue architecture from gene expression profiles of dissociated single-cells by learning spatial priors over spatial transcriptomic data. The model learns cell representations that allow for accurate positioning of cells within the tissue context, even in the absence of direct spatial information. LUNA operates in a continuous physical space, unrestricted by a finite number of physical spots, thus enabling the generation of tissue structures with arbitrary spatial density. Moreover, LUNA can infer spatial priors of the tissue and adapt to the context of the test sample without requiring explicit slice alignment. This is achieved by conditioning on the global gene expression profile of all cells, whereas alternative methods rely on selecting a specific reference slice for spatial reconstruction.

LUNA has broad implications in the field of spatial biology and addresses existing technological limitations. LUNA can infer 2D spatial locations for dissociated single cells from scRNA-seq datasets, enabling reconstruction of complex tissue architectures. Reconstructing spatial structures from scRNA-seq data thus enables the recovery of both transcriptome-wide gene expression profiles and inferred spatial localization, thereby combining the strengths of both modalities. In addition, LUNA can generate locations for cells or spots missing spatial information in the existing spatial technologies such as Slide-tags [29]. LUNA could reduce the need for comprehensive imaging of all tissue slices. A smaller number of slices can be profiled spatially, and LUNA can be used to infer spatial information for the remaining slices given their gene expression profiles, lowering experimental costs. Finally, LUNA is a complementary tool for analyzing tissue spatial structures and can be used along with existing tools for spatial analysis such as Squidpy [66], Seurat V5 [67], STELLAR [68] and STalign [69], as well as with existing batch correction methods [44, 70], whose embeddings can be used to condition LUNA.

Currently, LUNA operates within a two-dimensional (2D) spatial framework, which does not fully capture the inherently three-dimensional (3D) nature of tissue architectures. This limitation stems primarily from the constraints of existing sequencing technologies, which are not yet capable of measuring tissue structures in full 3D detail. LUNA can be extended beyond 2D spatial reconstruction and with ongoing advancements in technology [71], datasets that provide rich molecular organization of tissues in 3D are likely to become more widely available in the near future.

Our current analysis has mainly focused on mouse brain cell atlases due to the abundance of available data. Applying LUNA to other tissues and model organisms, could reveal the extent to which generalizable spatial priors can be learned across different biological systems. A limitation of the current framework is its requirement for a shared gene panel between the reference and query datasets. Using existing foundation models [72, 73] for conditioning LUNA could help to map different gene panels into a common embedding space. While LUNA effectively generalizes to unseen cell types within the mouse brain atlas, its ability to generalize in a zero-shot setting is limited when the training and test slice distributions differ substantially. In such settings, LUNA would require additional fine-tuning.

We envision LUNA, as an AI-driven tissue reconstruction model, can contribute to our understanding of complex biological processes and generating the intricate architecture of tissues across diverse biological systems. As large-scale spatial datasets become available, generative tissue reconstruction models may lay the foundation for future virtual tissue models that capture the spatial and functional landscape of real tissues and simulate the effects of perturbations within the context of tissue architecture.

## Acknowledgements

We are grateful to Ramon Viñas Torné, Shuyang Fan, Clément Vignac, Yiming Qin, Ed Lein, Philippe Schwaller, Evan Macosko, Mor Nitzan and Aviv Regev for valuable discussions. Generative AI tools were used to assist with formatting references and to provide feedback on the manuscript. All suggestions were reviewed and verified by the authors, who made all changes themselves.

## Author Contributions Statement

T.Y. and M.B. designed the study, performed research, contributed new analytical tools and analyzed data. T.Y. and C.E. performed experiments and developed the software. N.M., J.F. and S.A. contributed to the codebase. P.F. contributed to the algorithmic framework. A.H., S.N., H.L. and G.L.M. performed the mouse experiments and generated the Xenium data. T.Y., A.D.M., A.H., G.L.M. and M.B. analyzed the Xenium data. T.Y. and M.B. wrote the manuscript with the input from other authors. M.B. supervised the research.

## Methods

### Overview of LUNA

LUNA is a diffusion-based generative model designed to generate spatial locations of cells from their gene expression. Using an attention-based mechanism, LUNA learns how each individual cell should attend to other cells according to their relevance to a cell’s own location and molecular features. During training, LUNA leverages spatial transcriptomics data with ground truth cell locations to learn to generate cell locations conditioned on gene expression. During inference, the input to LUNA is the gene expression of dissociated cells, and the model then generates cell locations from pure Gaussian noise conditioned on gene expression, enabling cell locations inference.

In the training phase, the input to LUNA is spatial transcriptomics data from single or multiple slices. LUNA learns cell embeddings that capture both the local and global tissue structure using a multi-head self-attention mechanism [22, 23] that enables cells to learn to attend to other cells. We designed the loss in LUNA as an SE(2)-invariant function [74–77], *i.e*., it is robust to arbitrary rotations and translations of the predicted locations, only enforcing the preservation of relative distances between cells within each slice.

Training LUNA involves two main steps: *(i)* a corruption process that iteratively adds noise to the ground truth cell locations, and *(ii)* a denoising process that generates the ground truth locations from the noise-corrupted locations. During the denoising process, LUNA learns to reverse the corruption process, gradually transitioning from noisier to less noisy cell locations. LUNA as a generative model is more robust to distribution shifts, compared to a regression model which is prone to overfitting to specific tissue distributions (Supplementary Fig. 35). Another benefit of using generative modeling lies in estimating uncertainty in predictions (Fig. 3 c-e, Fig. 4 h-i).

### Learning cell representations

LUNA learns cell representations that capture the relationship between the molecular features of individual cells and their spatial context within tissue. These embeddings form the foundation for predicting the locations of cells.

The model is built on the premise that a cell’s location is influenced not only by its own molecular features but also by those of other cells in the tissue. To effectively model these interactions, LUNA employs a multi-head self-attention mechanism [22, 23], which dynamically integrates information from all cells in the slice. This mechanism allows the model to assign attention scores to different cells based on their relevance to the prediction of each cell’s location, capturing both local and global cellular interactions. Once the cell embeddings are learned, LUNA projects each cell’s embedding into a 2-dimensional space to predict its locations.

Formally, LUNA takes as input *(i)* molecular features of cells **X**_*s*_ ∈ ℝ^*m×d*^ in a given slice *s* where *m* denotes the number of cells and *d* denotes the number of genes, *(ii)* time step *t* ∈ N uniformly sampled from 0 to the maximum diffusion time *T*, and *(iii)* corrupted cell locations 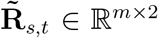 at time step *t*. The output of the model are the denoised cell locations 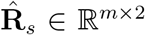 for a given slice *s*. We omit slice index *s* for the ease of notation.

LUNA is composed of multiple transformer layers, each integrating fully connected layers followed by the self-attention block. At each layer *l* given a fixed diffusion time *t*, LUNA learns *(i)* cell embeddings 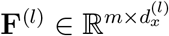, *(ii)* a diffusion time embedding 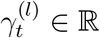 ;, and *(iii)* cell location embeddings 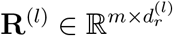 . The cell embeddings **F**^(0)^ in layer 0 are initialized via fully connected neural network layer that maps the molecular features of cells into the latent embedding space with dimensionality 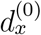 :

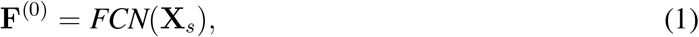

where *FCN* denotes a fully connected neural network layer. The diffusion time embedding 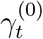 is initialized to the value of the diffusion time itself:

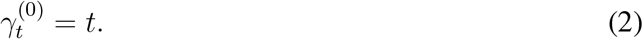

Finally, the cell location embeddings are initialized as the corrupted cell locations obtained from the corruption process at time step *t*:

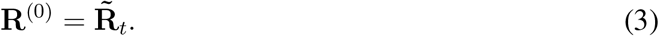

Then, at each layer *l* ∈ *{*1, …, *L}*, LUNA first transforms cell embeddings, a diffusion time embedding and cell location embeddings using fully connected layers:

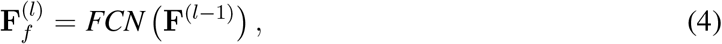

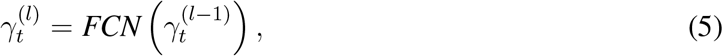

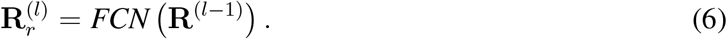

The transformed cell embeddings, diffusion time embedding and cell location embeddings are concatenated to form a unified representation for each cell:

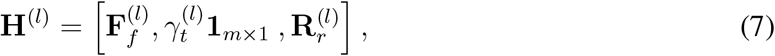

where **1**_*m×*1_ is a column vector of ones used to replicate the diffusion time embedding 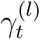 across all *m* cells. To obtain final cell embeddings **F**^(*l*)^ in layer *l*, the unified representation **H**^(*l*)^ is passed through a self-attention block which computes attention weights between all pairs of cells [23]:

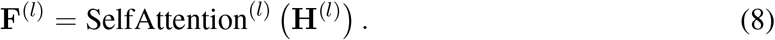

This self-attention operation enables the model to aggregate information from all cells in a slice, capturing how each cell’s location is influenced by others. To ensure scalability, we implemented the efficient attention [23] – an approximation of attention computation with a linear complexity. This approach reduces memory usage and computational complexity by circumventing the need for explicit pairwise interactions between every query-key pair (Supplementary Note 9).

Finally, the cell embeddings **F**^(*l*)^ are used to predict the locations of each cell. The cell locations are generated from the embeddings using a fully connected layer that projects the location of each cell from the latent embedding space to the two-dimensional physical space:

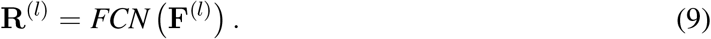

At each layer, the updated locations are centered by subtracting the mean, ensuring that the cell locations remain stable and consistent relative to the tissue structure. At the final layer, the output cell locations are the predicted clean cell locations 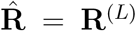. To better illustrate the scalability of our model components, especially the efficient attention mechanism and the FCN layer, we conducted kernel-level profiling of LUNA (Supplementary Note 10 and Supplementary Table 3).

### Loss objective

To optimize LUNA, we introduce a specific loss objective that is SE(2)-invariant. This geometric invariance is crucial because the spatial arrangement of cells in the tissue may undergo transformations due to experimental artifacts, while the underlying gene expression profiles remain unchanged.

To ensure SE(2)-invariance, LUNA introduces a pairwise loss function which focuses on preserving the relative distances between all cells in the slice defined as follows:

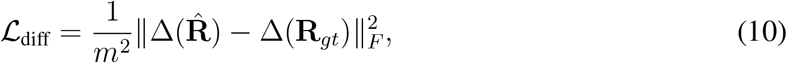

where Δ(*·*) : ℝ^*m×*2^ *→* ℝ^*m×m*^ computes the pairwise squared Euclidean distances between the locations of all cells and **R**_*gt*_ ∈ ℝ^*m×*2^ represents the ground true cell locations.

The function 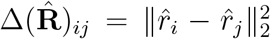 computes the squared distance between the predicted locations of cells *i* and *j*, and similarly, 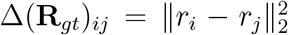 computes the squared distance between the true locations. The pairwise loss then becomes:

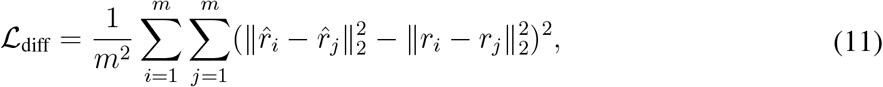

which penalizes deviations between the predicted and true pairwise distances, ensuring that the spatial relationships between cells are preserved.

The final loss is computed over all slices *s* ∈ {1, …*S*} where *S* denotes the total number of slices:

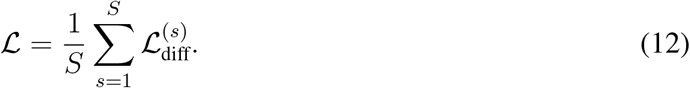

We further conducted a comprehensive ablation study on LUNA’s model components which confirms that LUNA’s attention-based architecture is crucial for capturing complex spatial dependencies between cells, while the pairwise loss function plays a key role in maintaining relational consistency (Supplementary Note 8, Supplementary Fig. 36).

### Corruption process

The diffusion model training begins with a corruption process, where LUNA progressively introduces noise to the true cell locations in an autoregressive manner. This step gradually transitions the true cell locations into pure noise, with the resulting corrupted locations serving as inputs for the subsequent denoising process.

We denote the initial diffusion time step, where no noise has been introduced to the cell locations, as time step 0, where ground-truth spatial information is fully retained. Conversely, the maximum diffusion time step, denoted as *T*, represents the state where no ground-truth information is preserved, and the locations are sampled from pure Gaussian noise. The time step *t* is randomly selected from *{*0, 1, 2, …, *T}*. The corrupted cell locations at time step *t* for the slice *s* are denoted as 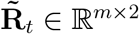 where again we omit the slice index *s* for the ease of notation. From time step *t –* 1 to *t*, Gaussian noise is added to the corrupted cell locations 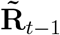, yielding noisier locations 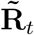 according to a predefined noise scheduler [24, 25, 78]. This noise scheduler is controled by the noise schedule exponent, which controls how much noise is injected or removed at each timestep of the reverse diffusion process and governs the randomness during sampling [79]. At time step *T*, the true cell locations 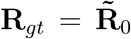 are transformed into corrupted cell locations 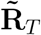 that have standard Gaussian distribution. The conditional distribution is defined as:

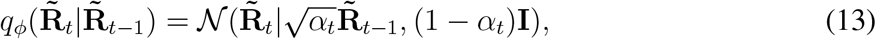

where *q*_*ϕ*_(*·*|*·*) represents the corruption process with a noise scheduler *ϕ* that conditionally transitions the cell location distribution from less corrupted to more corrupted states. *N* (*·*|*·, ·*) denotes a Gaussian distribution, while **I** is the identity matrix. The noise scheduler *ϕ*, regulated by hyperparameters *{α*_*t*_*}*, controls the balance between retaining the original cell locations and adding noise and governs the randomness during sampling.

To mitigate computational complexity, we compute the conditional distribution of corrupted cell locations given the true cell locations **R**_*gt*_, rather than from the previous diffusion step [25,80]:

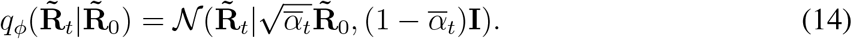

Sampling from this conditional distribution is equivalent to adding Gaussian noise to the true cell locations. As shown in [80], the corrupted cell locations are computed as:

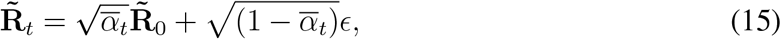

where 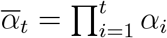, and *ϵ* is drawn from a Gaussian distribution with mean 0 and variance **I**, *i.e*., *ϵ* ∼ *N* (0, **I**). As *t* progresses from 0 to *T*, the cell locations become increasingly corrupted until they are indistinguishable from white noise. At *t* = *T*, the corrupted locations 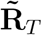 are sampled purely from Gaussian noise, with 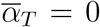. To ensure robustness to translations, we additionally subtract the center of mass from the noise *ϵ* and the data slices 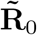, thus working with distributions defined in the subspace where the center of mass is fixed to 0 [81, 82].

### Denoising process

Given the corrupted locations, we train the LUNA model to recover the true cell locations by reversing the corruption process using gene expression profiles. We denote the model as *µ*_*θ*_(*·, ·, ·*) where *θ* represents all model parameters. LUNA progressively denoises the corrupted cell locations 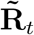 producing less corrupted locations 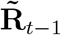 given time step *t*, more corrupted locations 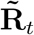 at the time step *t*, and the molecular features **X**.

Eventually, LUNA learns to generate clean cell locations from pure white noise, transitioning from *t* = *T* to *t* = 0. To achieve this, the model first predicts the clean cell locations 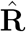 using the learned network *µ*_*θ*_(*·, ·, ·*). It then interpolates between the corrupted locations 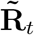 and the predicted clean locations 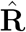, producing less corrupted locations for the next diffusion step:

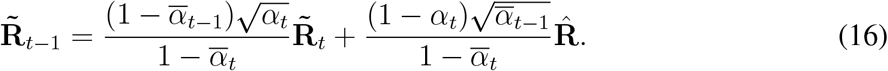

### Inference phase

Once the model *µ*_*θ*_(*·, ·, ·*) is trained, LUNA generates cell locations based on the molecular features of the cells in the test set 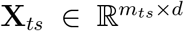 . The inference process starts by sampling locations from a normal distribution *ϵ* ∼ *N* (0, **I**), where 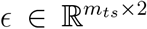 . The noisy locations, **R**_*ts,T*_ = *ϵ* are then refined through a sequence of diffusion steps.

At each step *t*, the model takes the noisy locations **R**_*ts,t*_, the molecular features **X**_*ts*_, and the time step *t* as inputs to predict the denoised locations for the next time step *t −* 1:

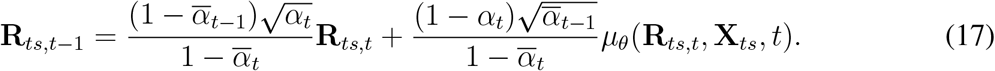

This process is repeated for each time step, iterating from *t* = *{T*, …, 0*}* until the model generates the final clean cell locations at *t* = 0.

### Hyperparameters

We use a consistent learning rate of 5 *×* 10^*−*4^ across all experiments, with a maximum diffusion time of 1000 steps and a cosine noise schedule characterized by *ν* = 2, which dictates the noise addition rate to the coordinates. A detailed analysis of hyperparameter selection and robustness to different values is provided in Supplementary Note 4 and Supplementary Fig. 37-38.

### Xenium Data Collection

We generate a new spatial transcriptomics dataset using Xenium. Animal husbandry and experimental procedures adhered to the Swiss Federal Veterinary Office guidelines and were authorized by the Cantonal Veterinary Office (cantonal animal license no. VD3499).

### Animal Procedures

C57BL/6J male mice were group-housed, with a maximum of four animals per cage, at 22 ^*°*^C and 50–60% relative humidity, under a 12-hour light–dark cycle. At eight weeks of age, mice were anesthetized with a Ketamine/Xylazine cocktail prepared in sterile saline. Once fully anesthetized, mice were placed in a stereotaxic frame. The head was shaved, Viscotears was applied to protect the eyes, and the scalp was disinfected with liquid betadine solution. After incision, connective tissue was removed to expose the skull and identify the bregma landmark for unilateral dorsal striatum injections.

Stereotaxic coordinates relative to bregma were: anterior-posterior +0.6 mm, medial-lateral +2.0 mm, and dorsal-ventral *−*2.6 mm. A small hole was drilled into the skull, preserving the dura mater, and 2 *µ*l of mouse wild-type pre-formed fibrils (PFFs) at a concentration of 2.5 *µ*g*/µ*l were injected into the right hemisphere at a flow rate of 0.4 *µ*l*/*min using a Hamilton syringe. The needle was left in place for 5 minutes post-injection before slow retraction to minimize backflow and was cleaned from blood to avoid clogging. The incision was sutured, betadine gel was applied, and animals were monitored in a heated recovery cage until fully awake. Post-operative well-being was assessed daily in accordance with animal license scoring sheets.

### Tissue Preparation for Xenium

Xenium was performed on four C57BL/6J mice: two wildtype controls and two *α*-synuclein PFF-injected animals harvested at three months post-injection (3MPI). One 3MPI animal was excluded prior to any downstream analysis because its coronal sections were not anatomically matched to the control sections, showing a systematic spatial offset in the sampled planes that precluded reliable cross-sample comparison. All analyses in this study, and the dataset deposited on Figshare, are based on the remaining three animals (two control and one 3MPI). Control and 3-months post-injected mice were euthanized via overdose of a Ketamine/Xylazine cocktail, and brains were rapidly removed. Brains were embedded in Tissue-Tek O.C.T. compound (catalog #4583) within 10 *×* 10 *×* 5 mm cryomolds (Tissue-Tek #4565) and frozen by submersion in dry ice-chilled (approximately *−*70^*°*^C) isopentane (Sigma #M32631). Frozen embedded brains were maintained on dry ice until transfer to *−*80^*°*^C storage.

Coronal sections (10 *µ*m) were cut on a Leica cryostat and mounted onto Xenium capture slides (10x Genomics, #10006559). For each brain, one coronal section targeting the amygdala (*−*1.70 mm relative to bregma) and one targeting the striatum (+0.6 to *−*1 mm relative to bregma) were collected. Two sections were placed per capture slide, which were returned to *−*80^*°*^C until processing. The predesigned 248-gene Xenium Mouse Brain Gene Expression panel was profiled across all sections on the Xenium *in situ* platform (10x Genomics).

### Post-Xenium Immunofluorescence

Following completion of the Xenium runs, slides were stored in PBS at 4^*°*^C. For immunofluorescence staining, tissues were permeabilized with PBS containing 0.5% bovine serum albumin (BSA) and 0.1% Triton X-100, blocked with 2% BSA in PBS, and then incubated overnight at 4^*°*^C with primary antibody against phosphorylated Ser129 *α*-synuclein (pSer129 *α*-synuclein, clone EP1536Y, Abcam ab51253, RRID:AB 869973) at 1/1000, followed by a secondary antibody (Invitrogen A32731, polyclonal, Alexa488) at 1/500.

Imaging was performed using a Leica DMi8 epifluorescence microscope equipped with a Lumencor SPECTRA X LED light source (nIR, 90-10172), a Leica DFC9000 GTC sCMOS camera (11547007), and a 20*×* HC PC APO objective (NA 0.8, air), yielding a pixel size of 0.34 *µ*m. Images were acquired with 10% tile overlap to cover the entire tissue section, collecting between 8 and 12 *z*-planes with 1 *µ*m spacing.

### Anatomical Registration

Anatomical registration was performed using Aligning Big Brains and Atlases (ABBA, https://go.epfl.ch/abba) [83], which combines affine transformations and nonlinear warping with anatomical border-guided feature registration. Images were automatically aligned to the Allen Brain Atlas CCFv3, and the alignment was subsequently manually refined to ensure precise anatomical correspondence.

### Immunofluorescence data processing

2D images were obtained from the acquired immunofluorescence stacks by performing maximum intensity projection channel-wise on the raw stacks. The immunofluorescence images were then stitched into a mosaic image using the *Ashlar* [84]. Only the DAPI channel was used to compute the corrected stitched coordinates. The immunofluorescence images were then aligned to the Xenium images by running the software *wsireg* [85], a Python wrapper of *elastix* [86, 87]. Each image transformation was computed using only the DAPI channel of both acquisitions, and the same obtained transformation was applied to the other channels of the immunofluorescence images (*i.e*. no warping was applied to the Xenium data in the process). To calculate the *α*-synuclein intensity level per cell, we first normalized the aligned *α*-synuclein channel by applying percentile-based normalization, in which we transformed the intensity values of the image so that the 1*st* percentile is mapped to 0 and the 99.8*th* percentile is mapped to 1. The results from the normalization were then clipped to the [0, 1] intensity range. The median normalized *α*-synuclein intensity per cell was then calculated by sequentially subsetting the normalized *α*-synuclein channel using the segmentation masks obtained from Xenium and aggregating accordingly.

### Cell Annotation

To annotate the Xenium data at the single cell level, we used the scRNA-seq data published in [88] as a reference for label transfer. We downloaded all the given single cell references (v2 and v3) and then concatenated them to obtain a single *h5ad* data file. During the concatenation process, we randomly subsampled the data by a factor of 4 (*i.e*., we keep 25% of the single cells) to alleviate posterior computational workload. We then ran *Tangram* [14] for each Xenium dataset using the pre-processed single cell data as reference. We ran *Tangram* in its default settings except for the run mode, for which we used *mode=“clusters”*. This mode runs the mapping at the cluster level and substantially reduces computational complexity. We annotated the Xenium data using 34 clusters.

## Data availability

MERFISH Whole Mouse Brain Atlas (ABC Atlas) for Animal 1 is available at https://alleninstitute.github.io/abc_atlas_access/descriptions/Zhuang-ABCA-1.html and for Animal 2 is available at https://alleninstitute.github.io/abc_atlas_access/descriptions/Zhuang-ABCA-2.html. MERFISH Mouse Primary Motor Cortex Atlas is available at the Brain Image Library: https://doi.brainimagelibrary.org/doi/10.35077/g.21. scRNA-seq Mouse Central Nervous System Atlas is available at the Single Cell Portal: https://singlecell.broadinstitute.org/single_cell/study/SCP1830. Slide-tags Datasets for all the tissues are available at the Broad Institute Single Cell Portal under the following accession numbers: SCP2170 (mouse E14), SCP2171 (human melanoma), SCP2169 (human tonsil) and SCP2167 (human brain). Xenium data on WT and 3MPI sections generated in this work is available at https://doi.org/10.6084/m9.figshare.33017210 [89].

## Code availability

LUNA was written in Python 3.9 using the PyTorch library. The source code is available on Github at https://github.com/mlbio-epfl/LUNA and archived on Zenodo at https://doi.org/10.5281/zenodo.21203881 [90] under the MIT license. The project website with links to data and code can be accessed at https://brbiclab.epfl.ch/projects/LUNA/.

