## Supplementary for "Tissue Reconstruction with Generative AI"

### Supplementary Note 1 Datasets and preprocessing

In LUNA, we considered the following datasets:

- **MERFISH Whole Mouse Brain Atlas (ABC Atlas)** [1]: We utilized raw gene expression data from Animal 1 (2.85 million cells) and Animal 2 (1.23 million cells) from the ABC atlas dataset. The data was log2-transformed, and $1,122$ genes were selected as input features. All slices from Animal 1 were used for training, and all slices from Animal 2 were used for testing. In the zero-shot setting, we trained LUNA on all slices from Animal 1 while excluding specific cell classes, and then applied LUNA to predict tissue structures in Animal 2, which included unseen cell types not present during training. Zero-shot experiments were conducted for each cell class with more than 1000 cells in the Animal 2 atlas, excluding HY Gnrh1 Glut (25 cells), MB Dopa (663 cells), and Pineal Glut (108 cells). All slices from Animal 1 were used for training, and all slices from Animal 2 were used for testing. The dataset provides hierarchical cell annotations at three levels: 4 classes: based on neurotransmitter identity (*e.g.*, glutamatergic, GABAergic); 34 classes: defined by brain region and neuronal vs. non-neuronal identity; 338 subclasses: fine-grained subtypes derived from unsupervised clustering within each major class.

- **MERFISH Mouse Primary Motor Cortex Atlas** [2]: This dataset consisted of raw gene expression data from 33 slices ($158,379$ cells) from one animal and 31 slices ($118,036$ cells) from another animal. After applying a log2 transformation, 254 genes were used as input features. We trained LUNA using all 33 slices from the first animal and tested it using all 31 slices from the second animal.

- **scRNA-seq Mouse Central Nervous System Atlas** [3]: Shi *et al.* [4] constructed the Mouse Central Nervous System (CNS) atlas by integrating STARmap PLUS measurements [4] with a published single-cell RNA-sequencing atlas [3]. This integration resulted in the imputation of expression profiles for $11,844$ genes and the estimation of spatial locations. We utilized this publicly available dataset from Shi *et al.* [4], which includes imputed transcriptomes and estimated spatial coordinates. By intersecting the gene panels from the ABC atlas and the CNS dataset, we identified 704 common genes. The CNS dataset comprises 13 coronal slices containing 1.08 million cells. Before integrating, both datasets were log2-transformed. For running LUNA, we integrated the gene expression matrices for these 704 common genes from both datasets—ABC Atlas (Animal 1) with 2.85 million cells and the CNS dataset with 1.08 million cells—using the Harmony method [5]. This integration, performed via the scanpy.external.pp.harmony_integrate function from the Scanpy library [6], resulted in a 600-dimensional latent space. The ABC dataset served as the training set, while the CNS dataset was utilized for testing. The effectiveness of the model was assessed by comparing the results to the estimated ground truth locations provided in the work of Shi *et al.* [4].

- **Slide-tags Datasets** [7]: For each cell class, the top 50 highly differential genes were selected, and we take the union of these highly differentially expressed genes resulting in the gene panel of 690 genes for mouse E14 tissue (4,623 spatially mapped nuclei, 4,414 spatially unmapped nuclei), 494 genes for the human melanoma tissue (4804 spatially mapped nuclei, 1662 spatially unmapped nuclei), 324 genes for the human cortex tissue (5778 spatially mapped nuclei, 3582 spatially unmapped nuclei) and 496 genes for the human tonsil tissue (4065 spatially mapped nuclei, 10100 spatially unmapped nuclei). A log2 transformation was applied for consistency with previous public datasets. Since the spatially unmapped nuclei lacked cell class labels, we trained a Linear Support Vector Classifier with the one-vs-rest strategy for multi-class classification (using "sklearn.svm.LinearSVC" with default parameters [8]) on the spatially mapped nuclei. We choose LinearSVC classifier because it achieves the highest classification accuracy across 5-fold cross-validation experiments over other alternatives such as KNN and logistic regression (Supplementary Figure 29). For each tissue, we randomly split the spatially mapped nuclei into a 90/10 train/test set with seed 42, using 90% of the mapped nuclei for training and 10% for validation. This classifier was then used to predict labels for the spatially unmapped nuclei, and these labels were used for visualization in the paper. When evaluating LUNA's performance on a held-out subset, we randomly split the spatially mapped nuclei into a 90/10 train/test set with seed 42. For generating locations for spatially unmapped nuclei, we used the same model trained on the spatially mapped nuclei and tested it on both mapped and unmapped nuclei.

- **Xenium dataset of Parkinson's disease model**: For experiments with fine-tuning, we used a LUNA model pretrained on the MERFISH ABC Atlas with 1,122 genes, utilizing all 147 slices from Animal 1 (2.85 million cells), and this model was then fine-tuned on 8 wild-type (WT) sections (494k cells) from our newly generated Xenium dataset. We applied the fine-tuned model to infer spatial coordinates for 4 diseased 3MPI sections (273k cells) that were held out during training. Xenium was performed on four male mice (two control and two 3MPI); one 3MPI animal was excluded at quality control (see Methods), leaving three animals analysed (two control and one 3MPI). For each animal, two coronal sections were collected, targeting the amygdala and the striatum. Each coronal section was divided along the midline into its left and right hemispheres, which were processed as separate sections; each animal therefore contributed four sections (two regions $\times$ two hemispheres). The 8 WT sections derive from the two control animals and the 4 3MPI sections from the single analysed 3MPI animal. For clarity, we visualize four representative sections in the paper, one hemisphere per region. During fine-tuning, we used the intersection of gene panels between the ABC Atlas and Xenium datasets, resulting in a shared set of 133 genes. For experiments without fine-tuning, the model was trained only on the MERFISH ABC Atlas data using the 133 intersecting genes between the MERFISH ABC Atlas and Xenium datasets, and then applied directly to the Xenium pathological samples without any fine-tuning. To account for differing sequencing depths between MERFISH and Xenium technologies, we started from the raw gene counts of both datasets, applied `scanpy.pp.normalize_total` to normalize per-cell library sizes to 500, followed by a `log1p` transformation [6]. We note that the Xenium gene panel was carefully curated to balance the cellular heterogeneity required for spatial reconstruction with pathological relevance. Moreover, this curated Xenium gene panel consistently outperforms randomly selected gene panels, confirming that it retains essential spatial information (Supplementary Fig. Supplementary Figure 39).

**Cell Segmentation.** For the MERFISH mouse cortex atlas, all MERFISH image analysis was performed using MERlin (available at https://github.com/ZhuangLab/MERlin). Cell boundaries in each field of view (FOV) were segmented using a seeded watershed approach [9], where DAPI images were used as seeds and polyA signals were used to identify segmentation boundaries. For the ABC Atlas, all MERFISH image analysis was also performed using MERlin. Cell segmentation was carried out using the DAPI and total polyA-mRNA signals and a deep learning-based cell segmentation algorithm, Cellpose 2.0 [10]. For the STARmap PLUS CNS Atlas, the ClusterMap method [11] was applied to segment cells based on amplicons (mRNA spots), with quality control for gene spots and pre- and post-processing. For the slide-tags datasets, cells were segmented from the images using watershed segmentation in MATLAB (release 2021b), and the centroid of each segment was calculated.

All spatial locations were normalized to a range of $-0.5$ to $0.5$ and used as the ground truth for cell spatial coordinates.

### Supplementary Note 2 Baselines

We compared LUNA with seven existing tissue spatial reconstruction methods—novoSpaRc [12], Tangram [13], CytoSPACE [14], CeLEry [15], STALocator [16], scSPACE [17] and STEM [18] on MERFISH Mouse Primary Cortex Atlas dataset [2] with in total 254 genes. Notably, CeLEry is the only method capable of processing multiple slices simultaneously. Consequently, we trained CeLEry using all 33 slices from the first animal and tested it on all 31 slices from the second animal. For each method, we inferred locations for all slices from the test set, which includes 31 slices, under two scenarios:

1. For each slice in the test set, we randomly select one of the 33 slices from the training set as a reference using a fixed random seed.

2. For each slice in the test set, we calculate the cosine similarity between its cell class distribution and those of the 33 training set slices in LUNA. We then matched each test slice with the training slice that showed the highest cosine similarity which should be the best match as a reference slice.

We conducted extensive hyperparameter tuning for all methods. For novoSpaRc [12],we use the cell locations of the reference slice as the target mapping space. We conducted 50 trials of hyperparameter tuning to optimize the model's performance. During each trial, we explored different values for two key hyperparameters: $\alpha_{\text{linear}}$ and $\epsilon$. The parameter $\alpha_{\text{linear}}$, which controls the trade-off between prior spatial information and gene expression similarity, was tested across values ranging from $0.1$ to $1$ in increments of $0.1$. The $\epsilon$ parameter, which governs the convergence of the reconstruction algorithm, was varied across values $5 \times 10^{-5}, 1 \times 10^{-4}, 5 \times 10^{-4}, 1 \times 10^{-3}, 5 \times 10^{-3}, 1 \times 10^{-2}, 5 \times 10^{-2}, 1 \times 10^{-1},$ and 1. For Tangram [13], we run it at the cell level with "uniform" density prior for MERFISH data by calling "tg.map_cells_to_space" and explored different values for the learning rate. The learning rate was tested across $5\times10^{-5}, 1\times10^{-4}, 5\times10^{-4}, 1\times10^{-3}, 5\times10^{-3}, 1\times 10^{-2}, 5 \times 10^{-2}, 1 \times 10^{-1},$ and 1. For CytoSPACE [14], we run it under default settings of single-cell mode since it has no hyperparameters to tune. For CeLEry [15], we performed 50 trials for hyperparameter optimization. During each trial, we explored different values for the learning rate, the dimension of the latent embedding, and the number of layers. The learning rate was tested across $5 \times 10^{-5}, 1 \times 10^{-4}, 5 \times 10^{-4}, 1 \times 10^{-3}, 5 \times 10^{-3}, 1 \times 10^{-2}, 5 \times 10^{-2}, 1 \times 10^{-1},$ and 1. The dimension of the latent embedding was explored with values of $32, 64, 128,$ and $256$, while the number of layers was varied across $3, 4, 5, 7, 8, 9, 10$. For STALocator [16], we performed 50 trials for hyperparameter optimization. During each trial, we explored different values for the radius cutoff factor $(0.4, 0.5, 0.55, 0.6)$, the cosine loss weight $\lambda_{\text{cos}}$ $(1, 3, 5, 7, 9, 10)$, the sliced Wasserstein distance loss weight $\lambda_{\text{SWD}}$ $(1, 3, 5, 7, 9, 10)$, the latent loss weight $\lambda_{\text{lat}}$ $(1, 3, 5, 7, 9, 10)$, and the reconstruction loss weight $\lambda_{\text{rec}}$ $(0.001, 0.01, 0.05, 0.1)$. The number of training epochs was fixed at $10,000$. For scSPACE [17], we performed $50$ trials for hyperparameter optimization. During each trial, we explored different values for the number of hidden features $(128, 256, 512, 1024)$, batch size $(32, 64, 128, 256)$, activation function (`ReLU` or `sigmoid`), learning rate $(0.01, 0.001, 0.0001)$, and number of training epochs $(500, 1000, 2000)$. For STEM [18], we performed $50$ trials for hyperparameter optimization. During each trial, we explored different values for the learning rate $(0.01, 0.005, 0.002, 0.001)$, the Gaussian kernel bandwidth parameter $\sigma$ $(0.01, 0.1, 1, 10, 100)$ and the spatial smoothing parameter $\alpha$ $(0.5, 0.6, 0.7, 0.8, 0.9)$.

In the paper, we report the results from the hyperparameter configuration that achieved the best performance for each of the baseline models. Specifically, for novoSpaRc [12], the final selected configuration was $\alpha_{\text{linear}} = 0.8$ and $\epsilon = 5 \times 10^{-3}$. For Tangram [13], the final selected learning rate was $10^{-2}$ using the "uniform" density prior. For CytoSPACE [14], we used the default single-cell-mode settings throughout. For CeLEry [15], the final selected configuration was a learning rate of $10^{-2}$, a latent embedding dimension of $128$, and $5$ hidden layers. For STALocator [16], the final selected configuration was a radius cutoff factor of $0.6$, $\lambda_{\text{cos}} = 5$, $\lambda_{\text{SWD}} = 5$, $\lambda_{\text{lat}} = 10$, and $\lambda_{\text{rec}} = 0.1$. For scSPACE [17], the final selected configuration used the `sigmoid` activation, batch size $128$, learning rate $10^{-3}$, and $1{,}000$ training epochs with feature normalization enabled. For STEM [18], the final selected configuration was a learning rate of $10^{-3}$, $\sigma = 10$, and $\alpha = 0.7$, trained for $800$ epochs.

We further benchmark alternative methods on MERFISH ABC Atlas [1]. For each benchmarked method, we used the best-performing hyperparameter sets determined from experiments on the MERFISH mouse cortex atlas. Specifically, we used a slice from Animal 1 as the reference, and for each slice from Animal 2. We identified the best-matching reference slice from Animal 1 by computing the cosine similarity of cell-type distributions. For the zero-shot experiments, we report results for the top three baseline methods based on Spearman's rank correlation on MERFISH ABC Atlas: scSpace, CeLEry, and STALocator. For each baseline, cells from one unseen cell class were excluded from the reference section, and spatial locations were inferred for test sections containing also that unseen cell class. We also identified the best-matching reference slice from Animal 1 for the test slice by computing the cosine similarity of cell-type distributions.

### Supplementary Note 3 Evaluation

There are various ways to assess the quality of tissue spatial reconstruction results. A high-quality reconstruction should preserve the local neighborhood composition of each cell, accurately capture the spatial distribution across different cell types, reflect biologically meaningful spatial gradients for various genes, and be robust to artificial effects.

We evaluate the quality of the cell location predictions across these dimensions using several metrics:

**Spearman's Rank Correlation (SRC).** SRC quantifies the strength and direction of association between two ranked variables. In our study, SRC is computed by comparing the ranks of pairwise distances between predicted and actual spatial coordinates for a group of cells. Specifically, for each cell, we calculate the rank correlation between its predicted pairwise distances to all other cells and the actual (ground truth) pairwise distances:

$$\rho = 1 - \frac{6 \sum d_i^2}{n(n^2 - 1)}$$

where $d_i$ represents the difference between the ranks of each predicted and actual pairwise distance, and $n$ is the total number of pairwise comparisons (cell pairs) in the group.

For each slice, we compute the Spearman Rank Correlation (SRC) for all the cells. We then aggregate these correlations across all cells in the slice by calculating the median.

**Moran's I.** Moran's I is a measure of spatial autocorrelation used to assess whether the spatial distribution of gene expression is clustered, dispersed, or random. It quantifies the degree of similarity between gene expression values at nearby cells based on a weighted spatial graph. The formula for Moran's I is:

$$I = \frac{N \sum_{i,j} w_{i,j} z_i z_j}{S_0 \sum_i z_i^2}$$

where $N$ is the total number of cells, $w_{i,j}$ represents the spatial weight (distance) between cells $i$ and $j$, $z_i$ and $z_j$ are the deviations of the gene expression values from the mean for cells $i$ and $j$, respectively, and $S_0$ is the sum of all spatial weights. In our case, $w_{i,j}$ corresponds to the distance between cell $i$ and cell $j$, while $z_i$ and $z_j$ denote the expression levels of a specific gene in these cells. A high Moran's I value indicates spatial clustering of gene expression, whereas a low value suggests spatial dispersion.

**Precision.** The precision is for evaluating the performance of our model in predicting contacts between cells. In the contact prediction task, we classify pairs of cells as being in contact or not based on the distances between them. To define what constitutes a contact, we use a threshold determined by a specific percentile of the distance distribution, referred to as the **percentile**. For example, using the 10% percentile means we consider the top 10% closest cell pairs as contacts.

Once the threshold is established, we label cell pairs with distances less than the threshold as positive (contacts) and those with distances greater than or equal to the threshold as negative (non-contacts). With these labels, we calculate the **Precision** ($P$) measures the proportion of predicted contacts that are actual contacts and is defined as:

$$P = \frac{\text{True Positives}}{\text{True Positives} + \text{False Positives}}.$$

**Root Sum Square Deviation (RSSD).** The Root Sum Square Deviation (RSSD) is a metric used to quantify the total deviation between predicted and true cell positions in spatial data after optimal alignment. It provides an overall measure of how accurately the model predicts cell locations by comparing the predicted positions to the true ones.

Given two sets of two-dimensional points, the true positions $\mathbf{V} = \{v_1, v_2, \ldots, v_n\}$ and the predicted positions $\mathbf{W} = \{w_1, w_2, \ldots, w_n\}$, where both sets contain $n$ points (cells), the RSSD is calculated after aligning $\mathbf{W}$ to $\mathbf{V}$ using the Kabsch algorithm [19]. This algorithm finds the optimal rotation (and optionally translation) that minimizes the deviation between the two sets.

RSSD quantifies the total deviation between predicted and true cell positions after optimal alignment, thereby measuring how well the spatial relationships among cells are preserved. Formally, we define RSSD as:

$$\text{RSSD} = \sqrt{\sum_{i=1}^{n} \|v_i - w_i'\|^2},$$

where $v_i$ denotes the true position of cell $i$, and $w_i'$ is its predicted position after alignment. A lower RSSD indicates that the predicted cell locations closely match the true spatial configuration. To capture performance at a finer granularity, we additionally compute *Per-Class RSSD* for each cell type $c \in \mathcal{C}$. For class $c$, let $\mathbf{V}_c$ and $\mathbf{W}_c$ represent the true and predicted positions of all cells of that type. The Per-Class RSSD is defined as:

$$\text{RSSD}_c = \sqrt{\sum_{i=1}^{n_c} \|v_i^c - w_i^{c\prime}\|^2},$$

where $n_c$ is the number of cells in class $c$, and $w_i^{c\prime}$ denotes the aligned predicted positions. To summarize performance across all cell types, we report the overall **RSSD** as the sum of the Per-Class RSSDs:

$$\text{RSSD} = \sum_{c \in \mathcal{C}} \text{RSSD}_c.$$

We adopt RSSD because it directly reflects spatial reconstruction fidelity by accounting local variations across cell types.

### Supplementary Note 4 Hyperparameters and Model Selection

**Hyperparameters.** In LUNA, we use a consistent learning rate of $5 \times 10^{-4}$ across all experiments, with a maximum diffusion time of $1000$ steps and a cosine noise schedule characterized by $\nu = 2$, which dictates the noise addition rate to the coordinates. The model consistently employs a latent dimension of $64$ for position encoding and utilizes $16$ attention heads in every experiment.

For large-scale atlas experiments such as the MERFISH Whole Mouse Brain Atlas, MERFISH Mouse Primary Motor Cortex Atlas, and scRNA-seq Mouse Central Nervous System Atlas, we implement $8$ transformer layers. In contrast, for smaller-scale studies like the Slide-tags experiments that focus on a single slice, we reduce the transformer layers to $2$.

The latent dimension of node features is set at $384$ for the MERFISH Whole Mouse Brain Atlas and the scRNA-seq Mouse Central Nervous System Atlas. For the smaller MERFISH Mouse Primary Motor Cortex Atlas and Slide-tags experiments, we adjust the latent dimension to $256$.

**Model Selection.** For large-scale datasets, including the MERFISH Whole Mouse Brain Atlas and the scRNA-seq Mouse Central Nervous System Atlas, we adopt a model trained for up to $3500$ epochs. This extensive training period accommodates the complexity and size of these datasets. Conversely, for smaller datasets such as the MERFISH Mouse Primary Motor Cortex Atlas and Slide-tags experiments, the model is sufficiently trained after $1000$ epochs, optimizing performance without overfitting to limited data.

**Hyperparameter Tuning Analysis.** To study the hyperparameter sensitivity of LUNA, we conducted hyperparameter tuning on the number of diffusion steps (250, 500, 1000), The noise scheduler exponent (1, 1.5, 2), hidden dimensions of the MLPs used for diffusion time embedding (64, 128, 256) and for position embedding (64, 128 ,256) (Supplementary Figure 37 and Supplementary Figure 38). Specifically, we divided all slices from one animal into training and validation sets by randomly splitting the slices in a 90/10 ratio: $90\%$ of the slices were used for training while $10\%$ of the slices were used for validation during hyperparameter tuning. For all the slices from the other animal, they serve as an independent test set. During grid search, we trained the model on slices from the train set and monitored performance on the validation slices. Once the optimal hyperparameters were selected, we fixed these parameters for all subsequent experiments across different datasets to ensure consistency and avoid overfitting to any specific data split.

As expected, increasing the number of diffusion steps leads to higher-quality sample generation. Adjusting the noise scheduler exponent affects the results subtly by encouraging the model to focus more on denoising either earlier or later in the diffusion process, depending on whether the scheduler is smoother or sharper. Also, we found that increasing the time step embedding size to match or exceed the other components leads to modest improvements in spatial accuracy while $64$ dimension is the best embedding size for spatial information embedding among the choice of $128$

and $256$. In our model, we fix diffusion steps as $1000$, noise scheduler componenet as $2$, the MLPs used for diffusion time embedding as $256$ and for position embedding as $64$ for all the experiments.

### Supplementary Note 5 Spatially Variable Gene Detection

We identified spatially variable genes (SVGs) for datasets in the paper using either SpatialDE or Moran's I statistics. The choice of method for different datasets was driven primarily by computational considerations and data preprocessing limitations. SpatialDE is known to be more accurate and statistically robust for identifying spatially variable genes, particularly in smaller or moderately sized datasets. In the Slide-tags datasets which have smaller number of cells, we used SpatialDE to obtain better precision in the SVG detection. However, SpatialDE is more computationally intensive and does not scale well to large atlas-scale datasets, such as those used in our MERFISH and ABC Atlas experiments. For these larger datasets, we used Moran's I statistic as a more scalable and efficient alternative.

**scRNA-seq CNS atlas.** We identified SVGs in the reconstructed tissue using the LUNA model from the scRNA-seq CNS atlas, employing Moran's I values for analysis. Notably, unlike SVG identification for Slide-seq data, SpatialDE was not applicable for this scRNA-seq CNS atlas. SpatialDE typically requires processing from raw gene counts; however, the scRNA-seq atlas only provided aggregated and z-scored gene expression data after integration with STARmap Plus.

For the reconstructed tissue slice analyzed with LUNA, we initially grouped cells by their classes, filtering out any class containing fewer than $50$ cells within the slice. This resulted in $25$ viable cell classes from an initial total of 27 for the slice shown in Figure 4. We further excluded the $704$ genes used in training from the scRNA-seq atlas gene library, leaving $11,140$ genes from an initial set of $11,844$ for SVG analysis using Moran's I values. For each cell class, we identified and listed the top five genes with the highest Moran's I values (see Supplementary Table 2).

**Slide-tags human melanoma sample.** We identified spatially variable genes (SVGs) in a Slide-tags human melanoma sample using SpatialDE [20]. SpatialDE employs a Gaussian Process (GP) framework to explore spatial dependencies in gene expression data, analyzing expression values at spatial locations.

Using LUNA, the number of identified nuclei in tumor cells increased from $899$ spatially mapped nuclei to $1,250$, including $263$ nuclei from tumor_1 cell type and $88$ from tumor_2 cell type. Out of a total of $36,601$ genes, we first excluded genes expressed in fewer than $10\%$ of all tumor cells ($125$ out of $1,250$), resulting in $9,413$ genes that were expressed in at least $10\%$ of tumor cells. We applied SpatialDE to analyze these genes in both the spatially mapped nuclei and the nuclei mapped with LUNA.

Given that SpatialDE assumes normally distributed noise, we adhered to the preprocessing pipeline recommended in the SpatialDE tutorial. We initially transformed the gene count data to approximate normally distributed noise using Anscombe's transform via NaiveDE.stabilize(), followed by linear regression with NaiveDE.regress_out() to adjust for biases caused by library size or sequencing depth before performing the spatial tests. We then ran SpatialDE.run() using default settings on the preprocessed gene expression and the locations of nuclei. The locations of nuclei are always normalized between $-0.5$ and $0.5$.

We assessed statistical significance using the $q$ value, which, unlike the $p$ value that measures the false positive rate, evaluates the false discovery rate [21]. We set the threshold for statistical significance at a $q$ value of less than $0.01$. Using this criterion, we detected $478$ significant genes using gene counts and nuclei locations from spatially mapped nuclei, and $833$ genes from data processed with LUNA.

**Fraction of Spatial Variance.** In SpatialDE, significant spatial variation is quantified using different covariance functions to test alternative hypotheses of expression patterns. This approach includes calculating the Fraction of Spatial Variance (FSV), which represents the proportion of total variance attributed to spatial factors, as outlined in SPATIALDE SUPPLEMENTARY NOTE 1 [20]. The FSV, therefore, quantifies the extent to which spatial patterns contribute to the total variance observed in gene expression data. Typically, the FSV varies across different technologies, tissues, and cell types. For instance, structured data obtained from spot-level spatial transcriptomics generally exhibit higher FSV values compared to unstructured data derived from continuous physical spaces, such as those from Slide-tags technologies.

### Supplementary Note 6 Estimation of High Density Regions

Using the LUNA model, we sampled $150$ times with different random seeds and performed kernel density estimation on the predicted locations to estimate high-density regions. Please note that thanks to LUNA's linear time and memory complexity during inference, this is very cheap. For each individual cell, LUNA can estimate high-density regions (HDR) and thus model uncertainty in predictions (Fig. 4h).

To estimate the HDR level of an individual cell, given the set of its predicted locations

$$\mathbf{P} = \{\mathbf{p}_1, \mathbf{p}_2, \ldots, \mathbf{p}_n\},$$

we estimate the spatial density $f(\mathbf{p})$ of its predicted locations using Gaussian kernel density estimation (Gaussian KDE). The HDR at quantile $q \in (0, 1)$ is defined as the minimal area $A_q$ such that

$$\int_{A_q} f(\mathbf{p})\, d\mathbf{p} = q.$$

In practice, this is approximated by evaluating $f(\mathbf{p})$ over a dense spatial grid, sorting the values, and identifying the threshold density corresponding to the quantile $q$ such that the cumulative probability mass above this threshold reaches $q$. The HDR area is then computed as

$$\text{HDR}_q = \sum_{\mathbf{p}_i \in \text{grid}} \mathbb{I}\big(f(\mathbf{p}_i) \geq \tau_q\big) \cdot \Delta x \cdot \Delta y,$$

where $\mathbb{I}(\cdot)$ is the indicator function, $\tau_q$ is the density threshold corresponding to quantile $q$, and $\Delta x, \Delta y$ are the grid resolutions. In our implementation, we set the quantile $q = 0.8$, meaning the HDR contains 80% of the predicted locations. The grid size is set to 250,000, corresponding to a $500 \times 500$ spatial domain subdivided into a $500 \times 500$ grid, i.e., 250,000 total grid cells.

Therefore, a smaller HDR region implies lower uncertainty of a cell's location. In such a way, the model can identify regions in which locations of cells are interchangeable.

### Supplementary Note 7 Locally Distinct Cell Identification

We aimed to identify cells exhibiting locally distinct gene expression profiles between wild-type (CTR) and 3MPI samples in the Xenium dataset. To enable direct comparison, we jointly processed slices from both CTR and 3MPI samples during the inference stage of LUNA. This allowed LUNA to predict cells from both conditions into a shared physical coordinate space learned during training, facilitating consistent spatial interpretation across samples.

To detect global expression shifts, we computed the Jensen–Shannon Divergence (JSD) for each of the $133$ genes between CTR and 3MPI samples. Genes exhibiting statistically significant distributional differences were ranked by their JSD scores, and we selected the top $10$ genes for further analysis.

Next, we assessed local expression differences for each of the $10$ divergent genes at the single-cell level. For local analysis, we identified $k$-nearest neighboring cells ($k$=200) of each cell within either the CTR or 3MPI sample and averaged their gene expression. The local divergence for each cell was quantified as the difference between these two local averages. To enhance spatial coherence and reduce local noise, we applied a Gaussian smoothing kernel to the resulting local divergence scores. Specifically, we computed a weighted average of local divergence values from spatially proximal cells, using a Gaussian kernel with a standard deviation $\sigma = 0.005$.

To identify cells most affected by local transcriptomic shifts, we selected the top $25\%$ of cells with the highest local divergence for each of the $10$ genes. We then performed a majority vote across genes, retaining cells that were among the top $25\%$ for more than half of the divergent genes, *i.e.*, at least $6$ out of $10$. These highlighted cells represent regions of pronounced local gene expression changes and may correspond to spatially localized pathological alterations associated with the disease state.

### Supplementary Note 8 Model Architecture Ablation Study

To support the use of multi-head self-attention and the pairwise distance-based loss function, we conduct a comprehensive ablation where we retrain the LUNA model using two different model variants:

1. Replacing multi-head self-attention with a feedforward network (FCN) of comparable depth and width.
2. Replacing the pairwise MSE loss with standard MSE on coordinates.

We found that each component of LUNA contributes to overall performance (Supplementary Figure 36). For the MERFISH mouse cortex atlas, which contains samples with arbitrary rotations and translations, the rotation-invariant loss provides the most significant benefit. However, because the spatial distribution of tissues in this dataset is relatively simple, using an MLP for coordinate decoding does not result in a substantial performance drop. In contrast, for the ABC Atlas, where the spatial tissue architecture is more complex, replacing the attention-based decoder with an MLP leads to a dramatic decline in performance. Since all sections in the ABC Atlas are aligned in canonical orientation, the benefit of using a rotation-invariant loss over standard MSE is less emphasized. These findings confirm that LUNA's attention-based architecture is crucial for capturing complex spatial dependencies between cells, while the pairwise loss function plays a key role in maintaining relational consistency.

### Supplementary Note 9 Efficient Attention

LUNA learns cell representations that capture cellular interactions globally and locally across entire tissue slice. To ensure scalability, we implemented Efficient Attention [22]. Different from traditional dot-product attention mechanism, Efficient Attention is an approximation of attention computation with linear complexity.

**Dot-product Attention.** Dot-product attention [23], the traditional method for implementing attention mechanisms, captures pairwise dependencies between input elements by transforming each input feature vector $x_i \in \mathbb{R}^d$ into three distinct vectors: a query $q_i \in \mathbb{R}^{d_k}$, a key $k_i \in \mathbb{R}^{d_k}$, and a value $v_i \in \mathbb{R}^{d_v}$. These transformations are performed through linear layers, which apply learned weight matrices to the input feature vectors. The similarities between the query and key vectors are computed by taking their dot products, capturing the interactions between different input features.

In matrix form, the set of queries, keys, and values are represented as $Q \in \mathbb{R}^{n \times d_k}$, $K \in \mathbb{R}^{n \times d_k}$, and $V \in \mathbb{R}^{n \times d_v}$, respectively, where $n$ represents the number of input feature vectors (in our case, the number of cells). The core operation of dot-product attention is expressed as:

$$D(Q, K, V) = \rho\left(QK^{\mathsf{T}}\right)V$$

Here, $\rho(Y)$ is a normalization function, typically a row-wise softmax, denoted as $\sigma_{\text{row}}(Y)$, which normalizes the raw dot-product scores across each row of the matrix $Y = QK^{\mathsf{T}}$. This ensures that the attention scores are positive and sum to 1 across each row, allowing the model to focus on the most relevant keys for each query while still considering all input features. The row-wise softmax function $\sigma_{\text{row}}(Y)$ for a given row $i$ is defined as:

$$\sigma_{\text{row}}(Y)_i = \frac{\exp(Y_{ij})}{\sum_{j'} \exp(Y_{ij'})}$$

where $Y_{ij}$ represents the element in the $i$-th row and $j$-th column of the matrix $Y$. This transformation converts the raw similarity scores into normalized attention weights. The resulting attention weights are then used to compute a weighted sum of the value vectors $V$, where the weight assigned to each value vector is determined by its corresponding attention score.

Thus, the final output of dot-product attention becomes:

$$D(Q, K, V) = \sigma_{\text{row}}\left(QK^{\mathsf{T}}\right)V$$

This mechanism enables the model to dynamically adjust its focus on different parts of the input sequence, ensuring that it captures the most important interactions between elements while maintaining a global view of the data through the attention weights.

Dot-product attention is effective but requires calculating all pairwise similarities, which leads to high memory and computational demands. Specifically, the memory complexity is $O(n^2)$, and the computational complexity is $O(d_k n^2)$, where $n$ represents the number of input feature vectors. These complexities grow quadratically with $n$, making it challenging to scale to large inputs.

**Efficient Attention.** Efficient attention [22] addresses the scaling issues of dot-product attention by reinterpreting the keys as global feature maps, eliminating the need to compute similarities for each individual pair of queries and keys. Instead, it uses the keys to summarize the input globally, and this summary is then applied to the queries. The output of efficient attention is computed as:

$$E(Q, K, V) = \rho(Q)\left(\rho(K)^{\mathsf{T}} V\right)$$

Efficient attention reduces memory and computation by avoiding the computation of pairwise interactions for each query-key pair. Instead, it aggregates global context vectors based on the keys. This leads to the fact that efficient attention scales linearly with $n$, making it a more practical solution for large-scale data without sacrificing performance. When using softmax normalization, the overall memory complexity becomes $O(dn + d^2)$ and the computational complexity becomes $O(d^2 n)$, assuming that $d_v = d_k = \frac{d}{2}$. In practice, $d \ll n$, therefore the complexity of Efficient Attention becomes linear to the number of cells $n$. Thus, making efficient attention a better option for processing large-scale data, from a computational standpoint.

| Method | Memory Complexity | Computational Complexity |
| --- | --- | --- |
| Dot-Product Attention | $O(n^2)$ | $O(d_k n^2)$ |
| Efficient Attention | $O(dn + d^2)$ | $O(d^2 n)$ |

*Table 1*: Comparison of the memory and computational complexities of dot-product attention and efficient attention.

### Supplementary Note 10 Model Scalability

To evaluate the scalability of our model, we performed kernel-level profiling (Supplementary Table Supplementary Table 3). We trained the model on the MERFISH mouse cortex atlas using a single NVIDIA GeForce RTX 3090 GPU with a batch size of 6. Profiling was conducted with `torch.profiler`, monitoring `torch.profiler.ProfilerActivity.CUDA` during the model's forward pass.

The analysis shows that fully connected layers (FCNs), including `aten::linear` and `aten::addmm`, dominate runtime and memory usage, accounting for over 80,000 $\mu$s of CUDA time and 7.5 GB of memory across diffusion steps. Attention layers, based on `aten::einsum` and `aten::bmm`, are significantly lighter, consuming only approximately 26,000 $\mu$s compute time and under 1 GB of memory. This validates the efficiency of our linear attention backbone in handling large-scale point clouds. While FCNs remain the most computationally intensive component, our design ensures their use is minimal and critical, with no redundant geometric computation.

This efficient separation allows our model to maintain competitive runtime even on large point sets (*e.g.*, tens of thousands of points). Unlike existing equivariant models applied to point cloud generation, we avoid operators with quadratic computational complexity during inference by shifting the geometric computations to the loss function. This design keeps the inference efficient, and the loss remains lightweight, with minimal impact on overall training time.

**Supplementary Table 1:** Top 30 genes that significantly influence certain cell class, with the percentage contribution of each gene to the specific cell class.

| Gene Name | Cell Class | Impact (%) |
|---|---|---|
| Igfbpl1 | OB-IMN GABA | 5.83 |
| Cldn11 | OPC-Oligo | 5.60 |
| Nr2f1 | CTX-CGE GABA | 5.04 |
| Vgf | CTX-MGE GABA | 5.01 |
| Nefh | CTX-MGE GABA | 5.00 |
| Arx | OB-IMN GABA | 4.83 |
| Tmem176a | MH-LH Glut | 4.52 |
| Gad2 | CTX-CGE GABA | 4.44 |
| Nr2e1 | MH-LH Glut | 4.37 |
| Arhgap28 | MH-LH Glut | 4.37 |
| Chrnb3 | MH-LH Glut | 4.34 |
| Slc32a1 | CTX-CGE GABA | 4.32 |
| Vgf | CTX-CGE GABA | 4.23 |
| Penk | CTX-CGE GABA | 4.20 |
| Cdkn1a | MH-LH Glut | 4.17 |
| Sall3 | MH-LH Glut | 4.13 |
| Mcm6 | MH-LH Glut | 4.11 |
| Pdgfc | MH-LH Glut | 4.11 |
| Chrna2 | MH-LH Glut | 4.10 |
| Lcp1 | MH-LH Glut | 4.08 |
| Sla | MH-LH Glut | 4.07 |
| Ptprc | MH-LH Glut | 4.06 |
| Tbx3 | MH-LH Glut | 4.05 |
| Alox8 | MH-LH Glut | 4.03 |
| Wnt7b | MH-LH Glut | 4.03 |
| Slc38a1 | CTX-MGE GABA | 4.02 |
| Slc5a7 | MH-LH Glut | 4.01 |
| Lmod2 | MH-LH Glut | 4.01 |
| Ung | MH-LH Glut | 3.99 |
| Ctsc | MH-LH Glut | 3.99 |

**Supplementary Table 2:** Top 5 cell-class specific spatially variable genes in slice well06 from the scRNA-seq CNS atlas using LUNA's prediction.

| Cell Class | Spatially Variable Genes |
|---|---|
| Astrocytes | *CSPG5, PANTR1, TSPAN7, LRIG1, GNAO1* |
| Cholinergic and Monoaminergic Neurons | *CDR1, COX6A2, NDUFB6, ACYP1, COX6B1* |
| Choroid Plexus Epithelial Cells | *SPTBN1, EIF4G2, APP, MRPS34, ARL6IP1* |
| Dentate Gyrus Granule Neurons | *ATP2B4, CAMK2D, NNMT, COBL, ZBTB20* |
| Di- and Mesencephalon Excitatory Neurons | *GABRD, BOK, PLEKHG1, TIAM1, SLC9A3R1* |
| Di- and Mesencephalon Inhibitory Neurons | *NEFM, ANKRD34B, CTSB, MED10, GM27199* |
| Ependymal Cells | *RB1, PALB2, WRAP73, GM42595, ARMC7* |
| Glutamatergic Neuroblasts | *COX18, FAM150B, FGD4, D230017M19RIK, SBK1* |
| Hindbrain Neurons/Spinal Cord Neurons | *SLC18A3, MED10, KCNC1, BOK, ARPP19* |
| Microglia | *FOXJ1, CCDC108, 1500015O10RIK, TCTEX1D4, CFAP77* |
| Non-glutamatergic Neuroblasts | *GALM, GNS, CCDC148, CNTN1, PLEKHB1* |
| Olfactory Inhibitory Neurons | *PROX1OS, RWDD3, NEUROD1, ZBTB18, UCHL3* |
| Oligodendrocyte Precursor Cells | *NID1, SELPLG, NDUFA4L2, SPARC, CMTM7* |
| Oligodendrocytes | *MFNG, CYBA, FAM216B, UNC45B, ADGRL4* |
| Peptidergic Neurons | *SUB1, PLP1, HINT1, MOBP, WFDC18* |
| Pericytes | *4932443I19RIK, CDKL4, SLC14A1, CFAP61, KIF6* |
| Perivascular Macrophages | *TMEM56, HIST3H2BA, FBXO44, ANKRD39, APOE* |
| Subcommissural Organ Hypendymal Cells | *CADM1, FAM73A, CABP1, NDUFC2, AI987944* |
| Telencephalon Inhibitory Interneurons | *PLXNA2, GFRA2, EPHX4, GRIA3, NRSN2* |
| Telencephalon Projecting Excitatory Neurons | *RCN3, ZBTB20, IL16, GM2115, LOXL1* |
| Telencephalon Projecting Inhibitory Neurons | *NMB, WFDC18, UBASH3B, DOCK10, PDE10A* |
| Unannotated | *MCM2, LSM8, ACTL6A, RPA3, SAP30* |
| Vascular and Leptomeningeal Cells | *SLC4A10, A230005M16RIK, COL25A1, IL17RE, SMAD9* |
| Vascular Endothelial Cells | *ADAMTSL2, CSRP2, PLXND1, RSRP1, CD82* |
| Vascular Smooth Muscle Cells | *GADD45B, GADD45G, NTRK1, KCNH2, DKK3* |

**Supplementary Table 3:** CUDA operation profiling summary, including execution time, memory usage, and call count for selected operations. We train the model on MERFISH mouse cortex dataset on a single NVIDIA GeForce RTX 3090 GPU with batch size as 6.

| **Operation** | **CUDA Time ($\mu$s)** | **CUDA Memory Usage** | **Count** |
|---|---:|---:|---:|
| aten::linear | 43,028 | 4.20 GB | 138 |
| aten::addmm | 37,592 | 3.31 GB | 114 |
| aten::einsum | 13,401 | 599 MB | 16 |
| aten::bmm | 12,637 | 300 MB | 16 |

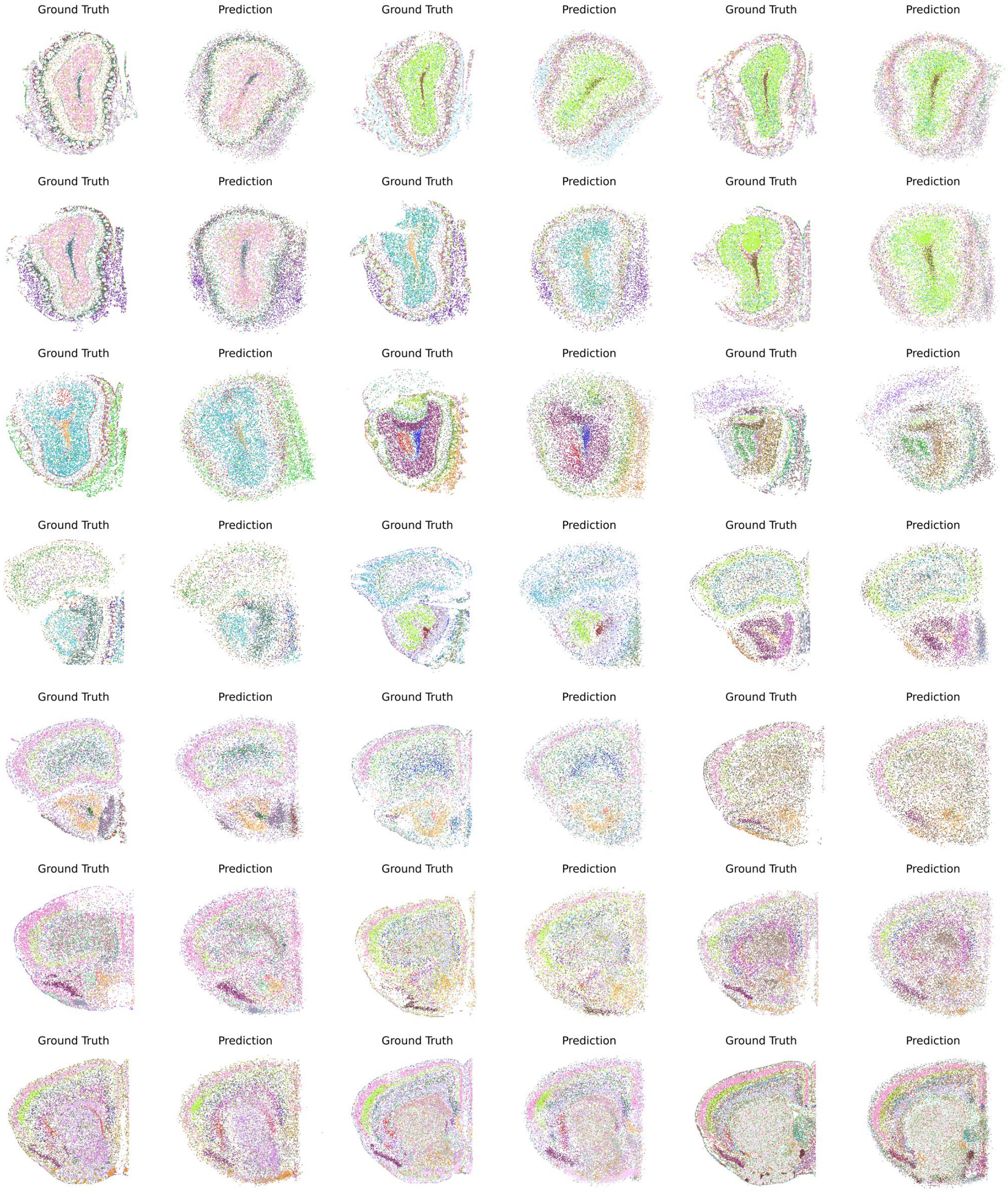

**Supplementary Figure 1:** Full tissue reconstruction results using LUNA from distinct major brain regions of the MERFISH whole mouse brain dataset (Allen Brain Cell [ABC] atlas). Cells are coloured according to their cell type annotation with 338 distinct cell subclasses.

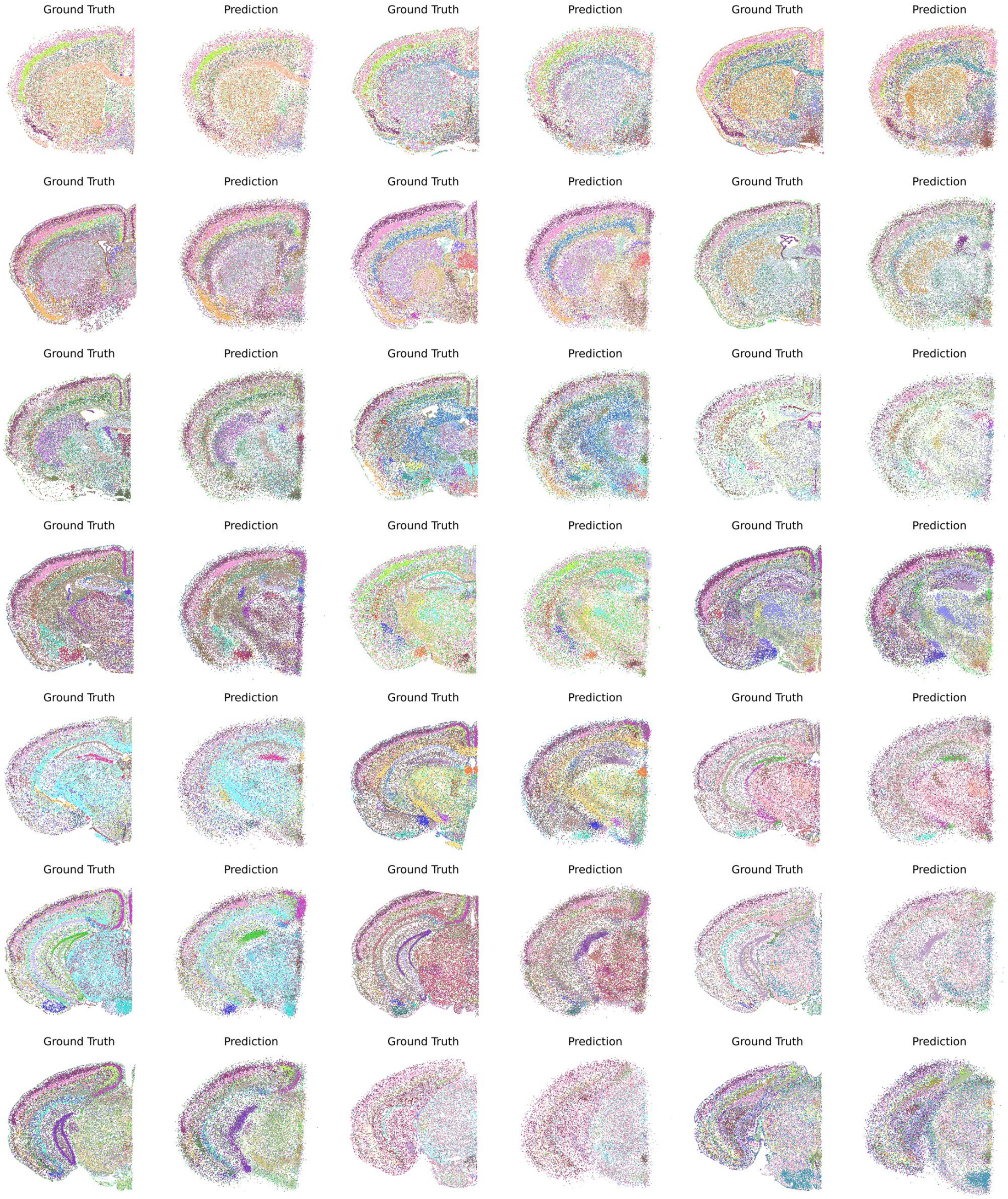

**Supplementary Figure 2:** Full tissue reconstruction results using LUNA from distinct major brain regions of the MERFISH whole mouse brain dataset (Allen Brain Cell [ABC] atlas). Cells are coloured according to their cell type annotation with 338 distinct cell subclasses.

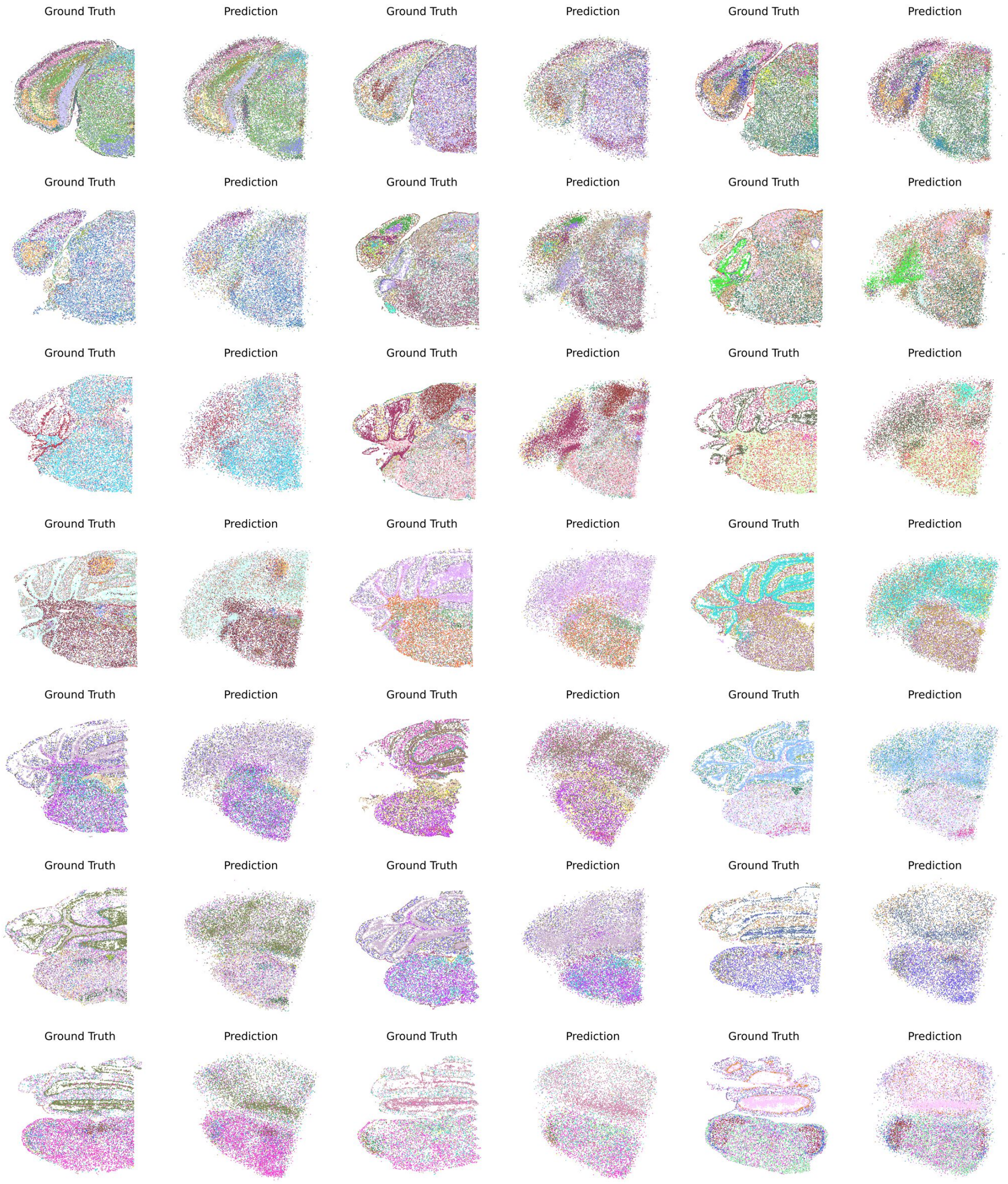

**Supplementary Figure 3:** Full tissue reconstruction results using LUNA from distinct major brain regions of the MERFISH whole mouse brain dataset (Allen Brain Cell [ABC] atlas). Cells are coloured according to their cell type annotation with 338 distinct cell subclasses.

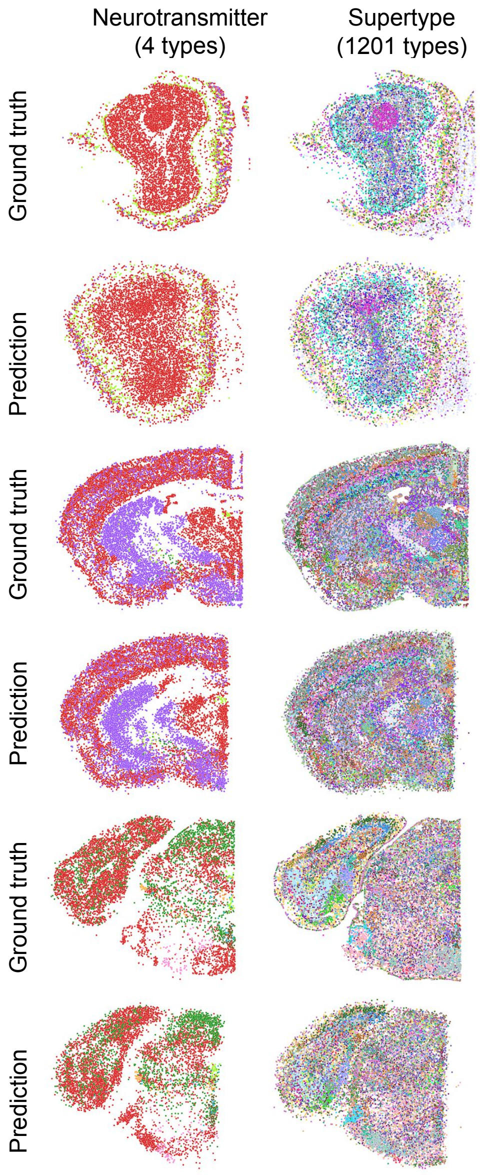

**Supplementary Figure 4:** LUNA's predictions for three example slices containing $12,610$ cells, $30,931$ cells, $22,010$ cells respectively, categorized by $4$ neurotransmitter types and $1201$ supertypes. Ground truth cell locations are shown in the top plot, while the predictions made by LUNA are displayed in the bottom plot.

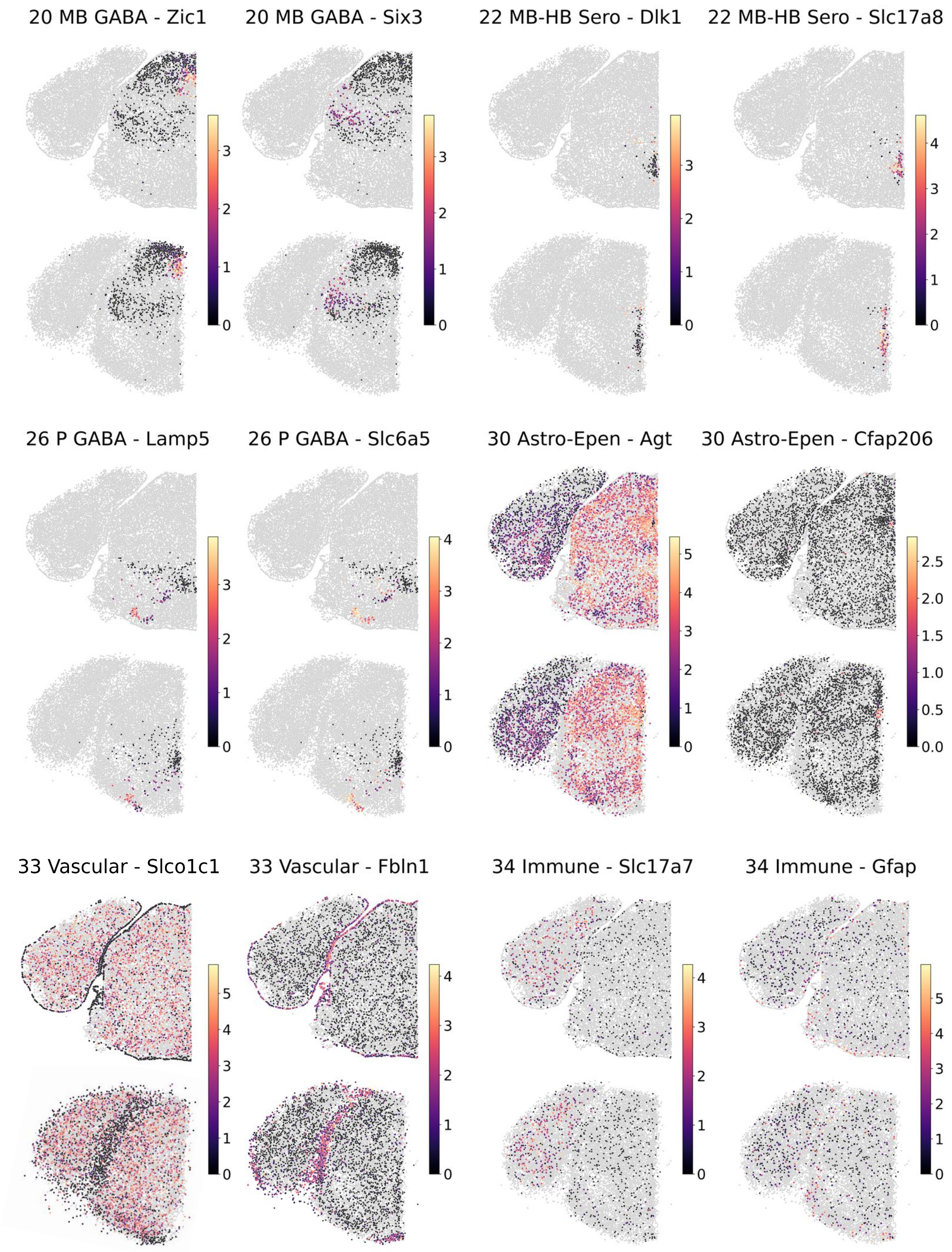

**Supplementary Figure 5:** Spatial expression patterns of different cell classes, including MB GABA cells, MB-HB Sero cells, P GABA cells, Astro-Epen cells, Vascular cells and Immune cells. For each cell class, we analyzed the spatial variability of $1,122$ genes by calculating Moran's I values, based solely on the ground truth locations of cells from the respective cell class. From this analysis, we identified the top two genes exhibiting the highest Moran's I values for each cell class and visualized those genes..

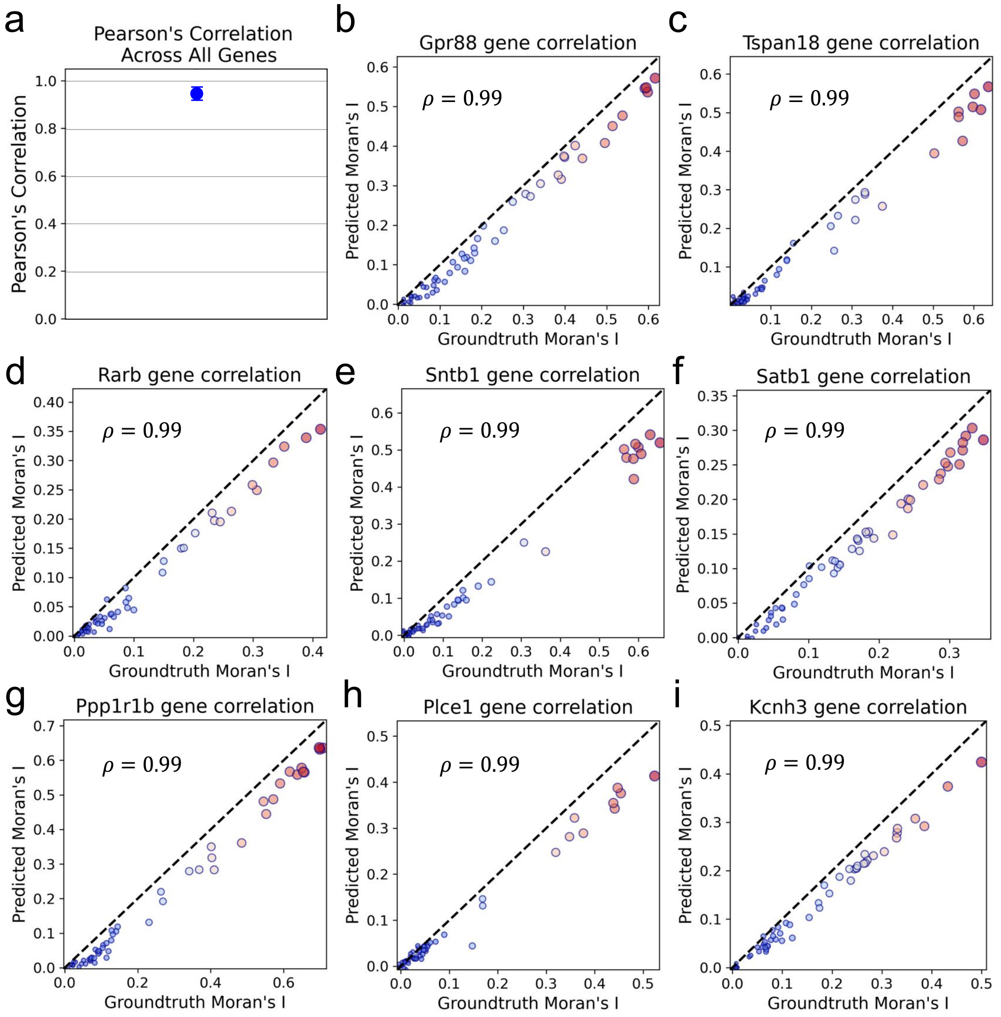

**Supplementary Figure 6: (a)** We computed Pearson's correlation for all 1122 genes across the 66 slices. For each slice, the Pearson's correlation was calculated for all 1122 genes, and the average correlation across the 66 slices was used as the central point. The error bars represent the standard deviation across slices. **(b-i)** The spatial autocorrelations for the top 8 genes ranked by the highest Pearson's correlation across all the slices are also listed. Each point represents a single slice out of the 66 total slices, with slices colored on a gradient from blue (low) to red (high) based on the ground truth Moran's I value. Points closer to the diagonal suggest better preservation of spatial patterns by LUNA. The Pearson's correlation coefficient ($\rho$) is displayed in the top left corner.

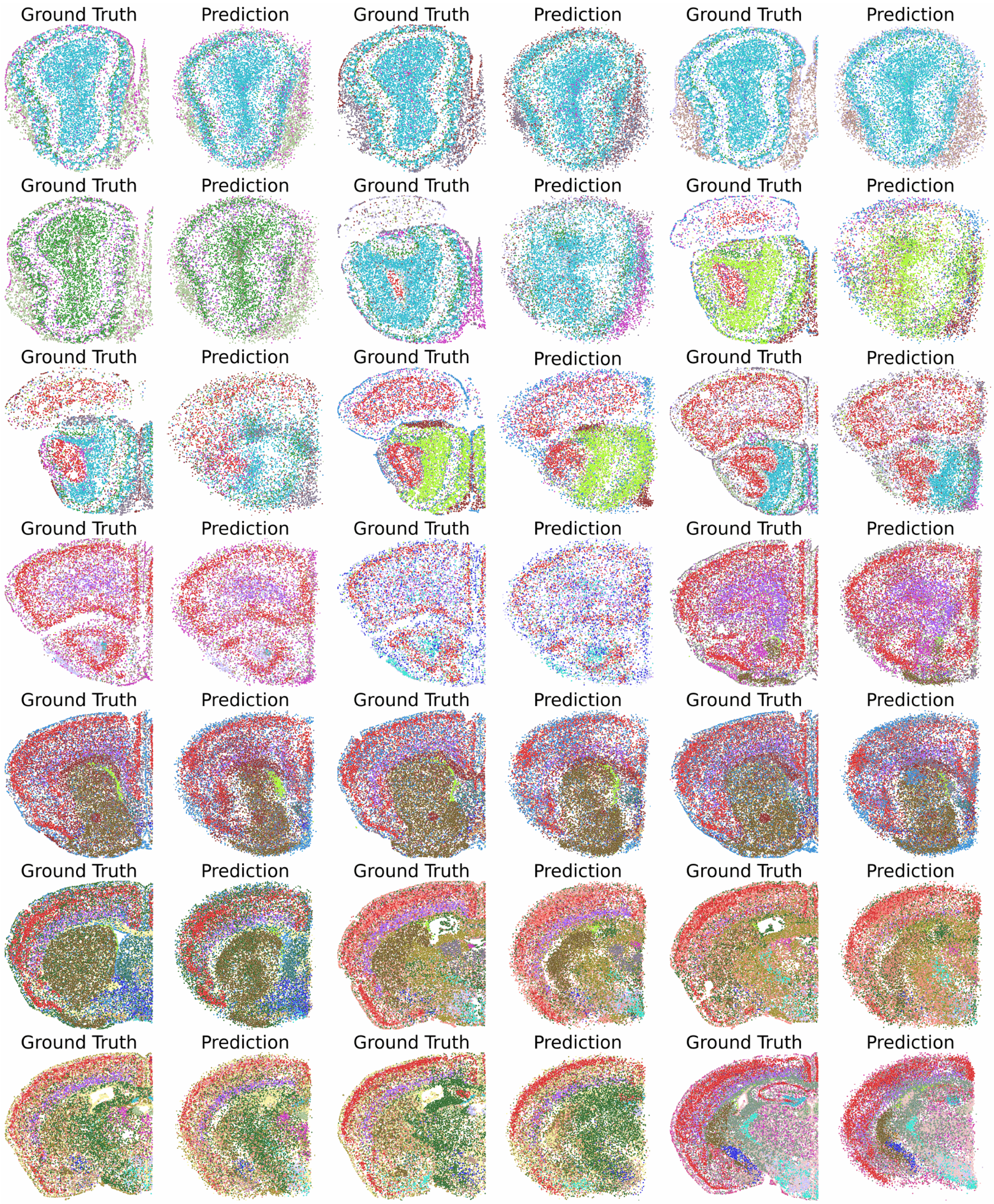

**Supplementary Figure 7:** Tissue reconstruction results on cross slice generalization using LUNA from distinct major brain regions of the MERFISH whole mouse brain dataset (Allen Brain Cell atlas). Cells are coloured according to their cell type annotation with 338 distinct cell subclasses. We train the model with 70% of randomly selected slices from Animal 1 and test on the remaining 30% of slices (41 slices).

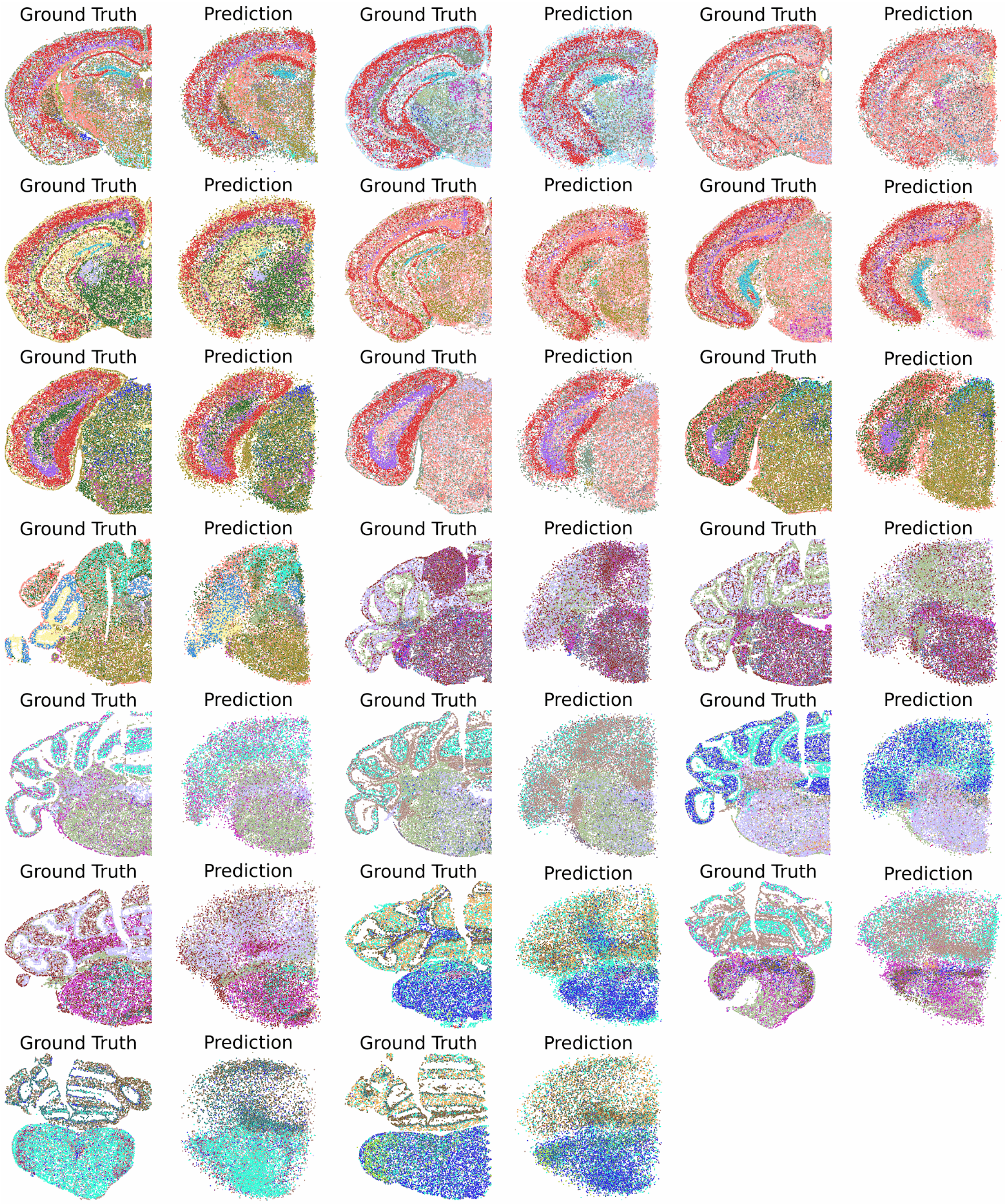

**Supplementary Figure 8:** Tissue reconstruction results on cross slice generalization using LUNA from distinct major brain regions of the MERFISH whole mouse brain dataset (Allen Brain Cell atlas). Cells are coloured according to their cell type annotation with 338 distinct cell subclasses. We train the model with 70% of randomly selected slices from Animal 1 and test on the remaining 30% of slices (41 slices).

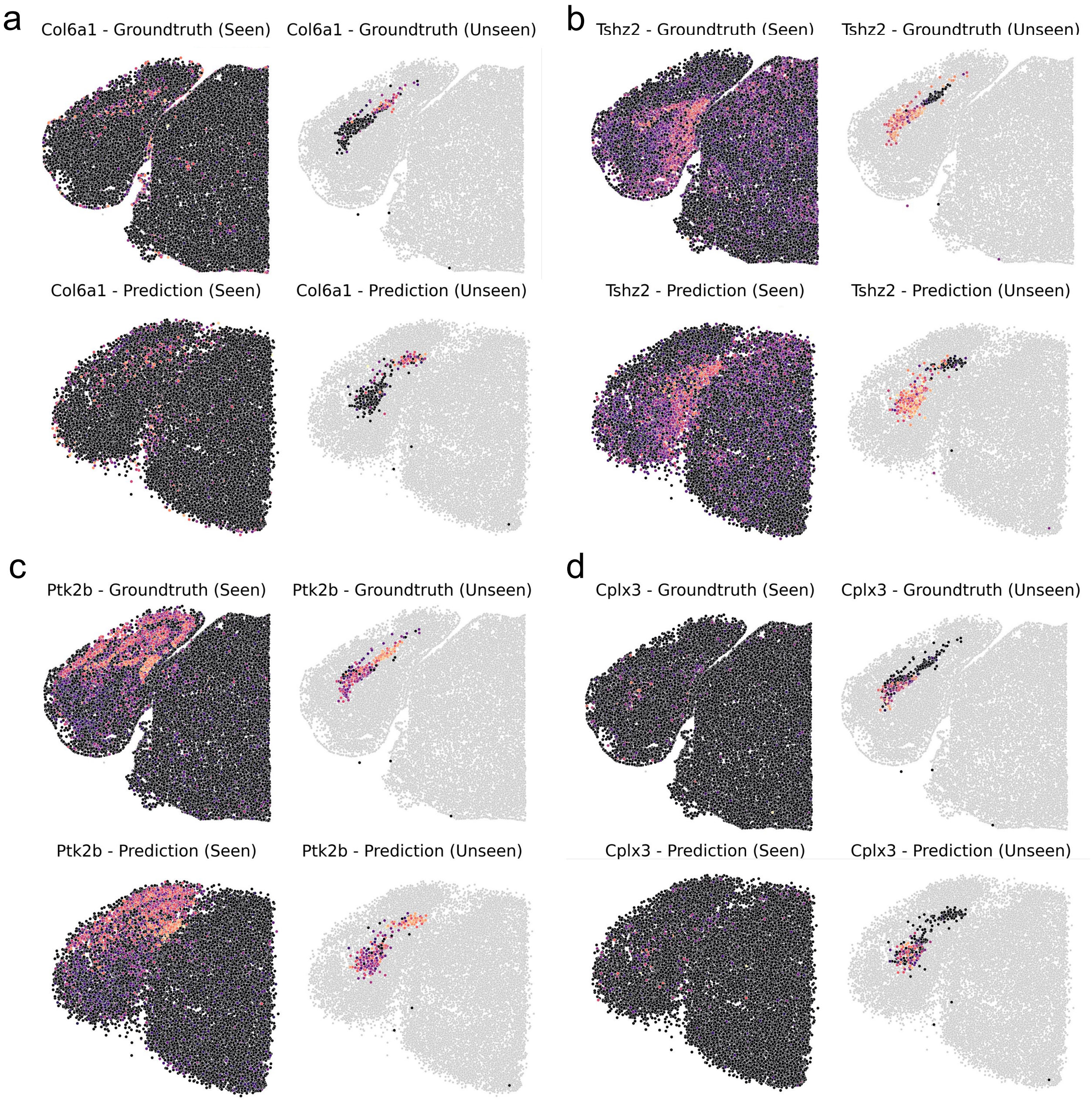

**Supplementary Figure 9:** Gene expression gradients in spatial structure for both seen cell classes and an unseen cell class (NP-CT-L6b Glut). Spatial expression patterns of **(a)** the *Col6a1* gene, **(b)** the *Tshz2* gene, **(c)** the *Ptk2b* gene, and **(d)** the *Cplx3* gene, which are shared across seen and unseen classes. For each gene, we visualize expression in seen cell classes (left) and the unseen NP-CTL6b Glut class (right), comparing ground truth spatial expression (top row) with LUNA's predicted expression maps (bottom row).

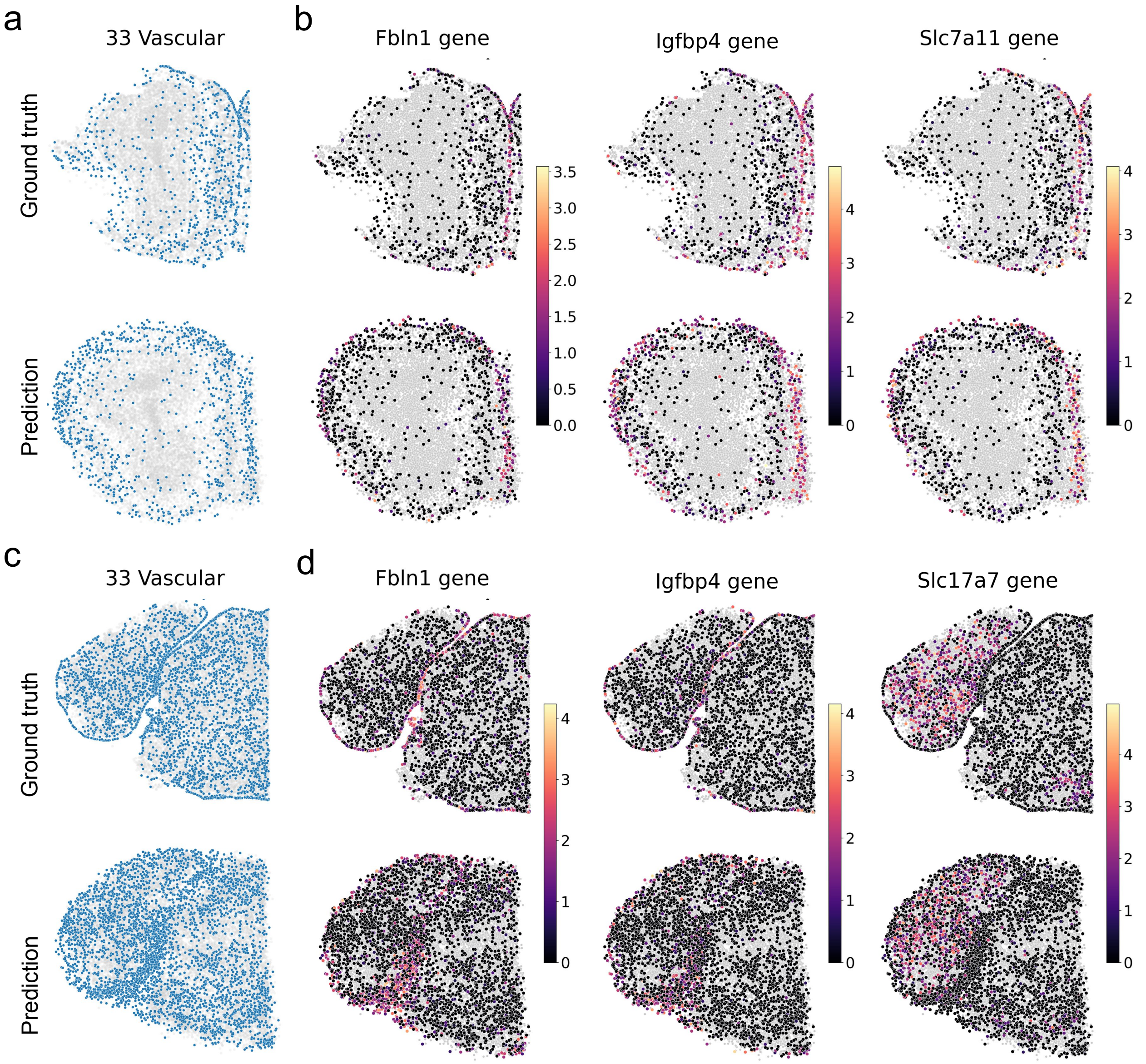

**Supplementary Figure 10:** **(a, c)** Tissue reconstruction results with LUNA on two slices for the excluded Vascular class, comprising $447,941$ cells from $34$ classes. LUNA was trained on $2.41$ million cells from Animal 1 in the ABC atlas and tested on all classes, including Vascular, in Animal 2 ($1.23$ million cells), which was not seen during training. **(a, c)** Displays show the spatial distribution of $998$ and $5,676$ cells in two slices, with ground truth at the top and LUNA's predictions below. **(b, d)** Displays show spatial expression of a gene in the Vascular class from the same slices, selected for its high Moran's I values from $1,122$ genes, notably *Fbln1*, *Igfbp4*, *Slc7a11*, and *Slc17a7*. Other cell classes are in gray.

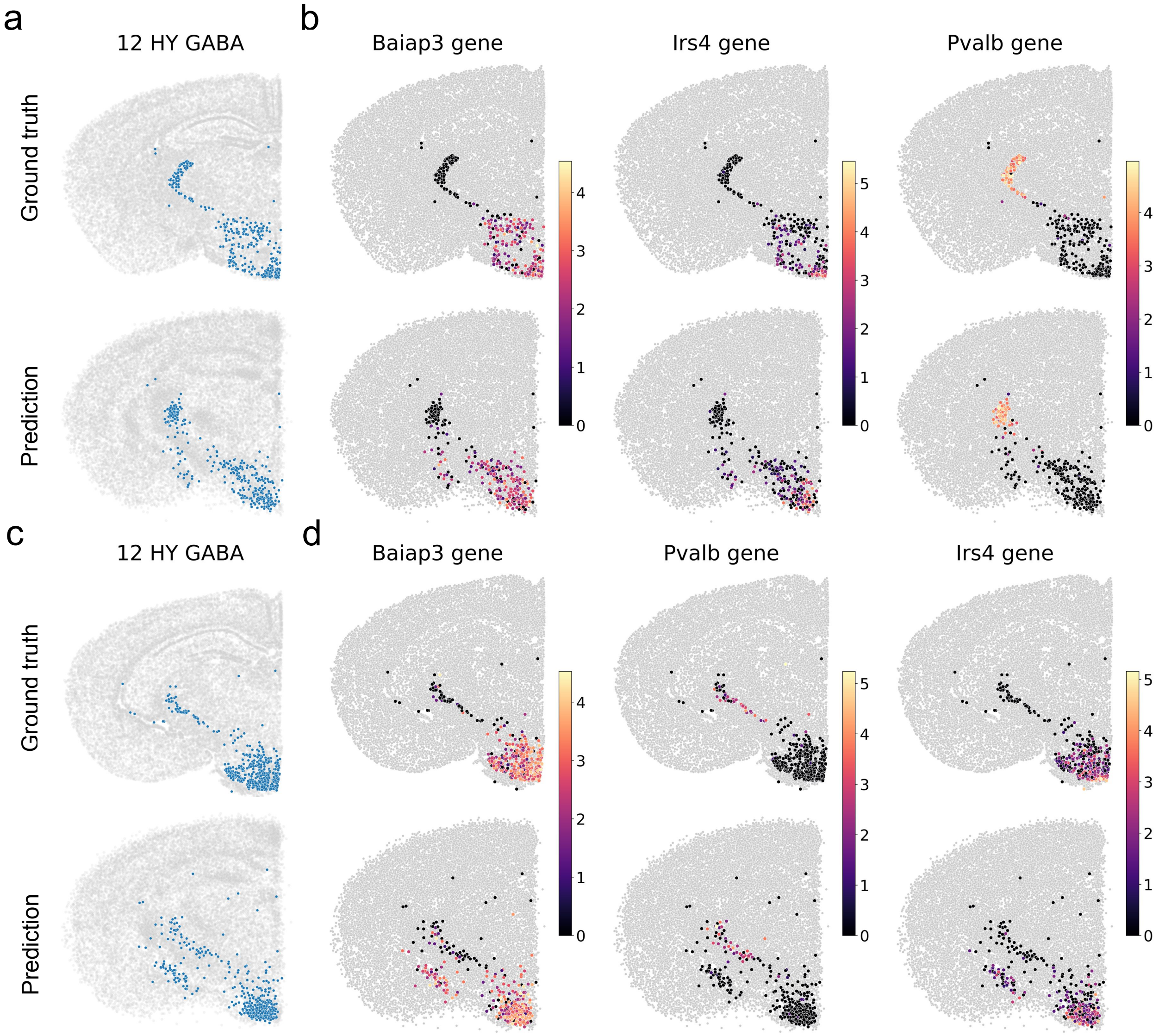

**Supplementary Figure 11:** **(a, c)** Tissue reconstruction results with LUNA on two slices for the unseen HY GABA class, which was excluded during training from $34$ classes totaling $15,055$ cells. LUNA trained on the remaining classes using $2.6$ million cells from Animal $1$ in the ABC atlas and applied to all classes in untrained Animal $2$ ($1.23$ million cells), including HY GABA. Visualizations show the spatial distribution of $333$ and $495$ cells in selected slices, respectively, with ground truth at the top and LUNA's predictions below. **(b, d)** Displays spatial expression of a gene in HY GABA cells from the same slices, selected for its high Moran's I values across $1,122$ genes, notably *Baiap3*, *Irs4*, *Pvalb*. Other cell classes are shown in gray.

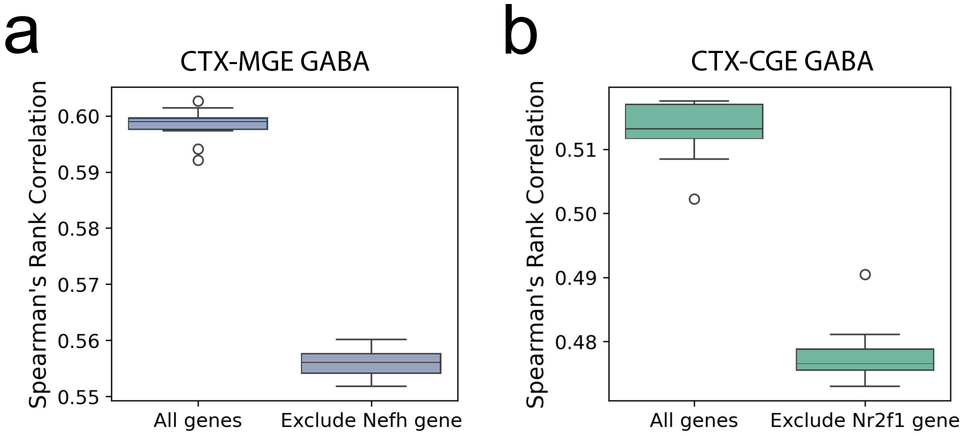

**Supplementary Figure 12:** The effect of different initializations of the initial noise during inference. We conduct $10$ experiments with different random seeds, focusing on genes that significantly influence specific cell classes. We compare SRC when using all genes with scenarios excluding **(a)** *Nefh* in CTX-MGE GABA and **(b)** *Nr2f1* in CTX-CGE GABA. Box plots show the performance across seeds, with the median (center), $25$th–$75$th percentiles (box bounds), and whiskers extend to the most extreme value within $1.5\times$ the IQR of the box edges, for each group.

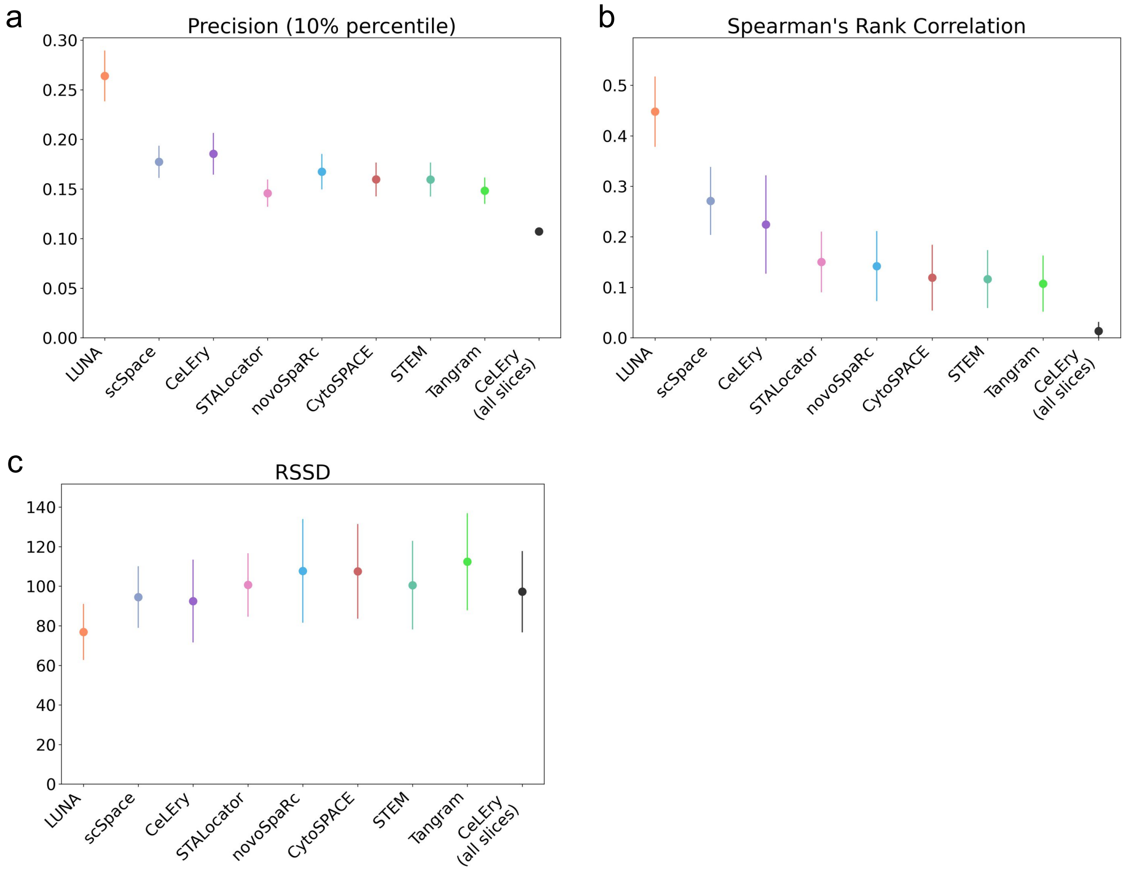

**Supplementary Figure 13:** Performance comparison on the cross-animal generalization task between LUNA and alternative baselines on the MERFISH mouse primary motor cortex atlas [2]. Models were trained on slices from one mouse and evaluated on all slices from an unseen mouse (n=$31$ slices, $118,036$ cells). We evaluated performance based on **(a)** precision at the $10$-th percentile threshold for predicting the closest cell pairs as contacts, **(b)** Spearman's rank correlation (SRC) across all cells within the same slice, and **(c)** Root Sum Square Deviation (RSSD) between the ground truth and predicted spatial coordinates. Error bars represent standard deviation across $31$ slices (n=$31$ slices). For precision and SRC higher values indicate better performance. For RSSD, lower values indicate better performance.

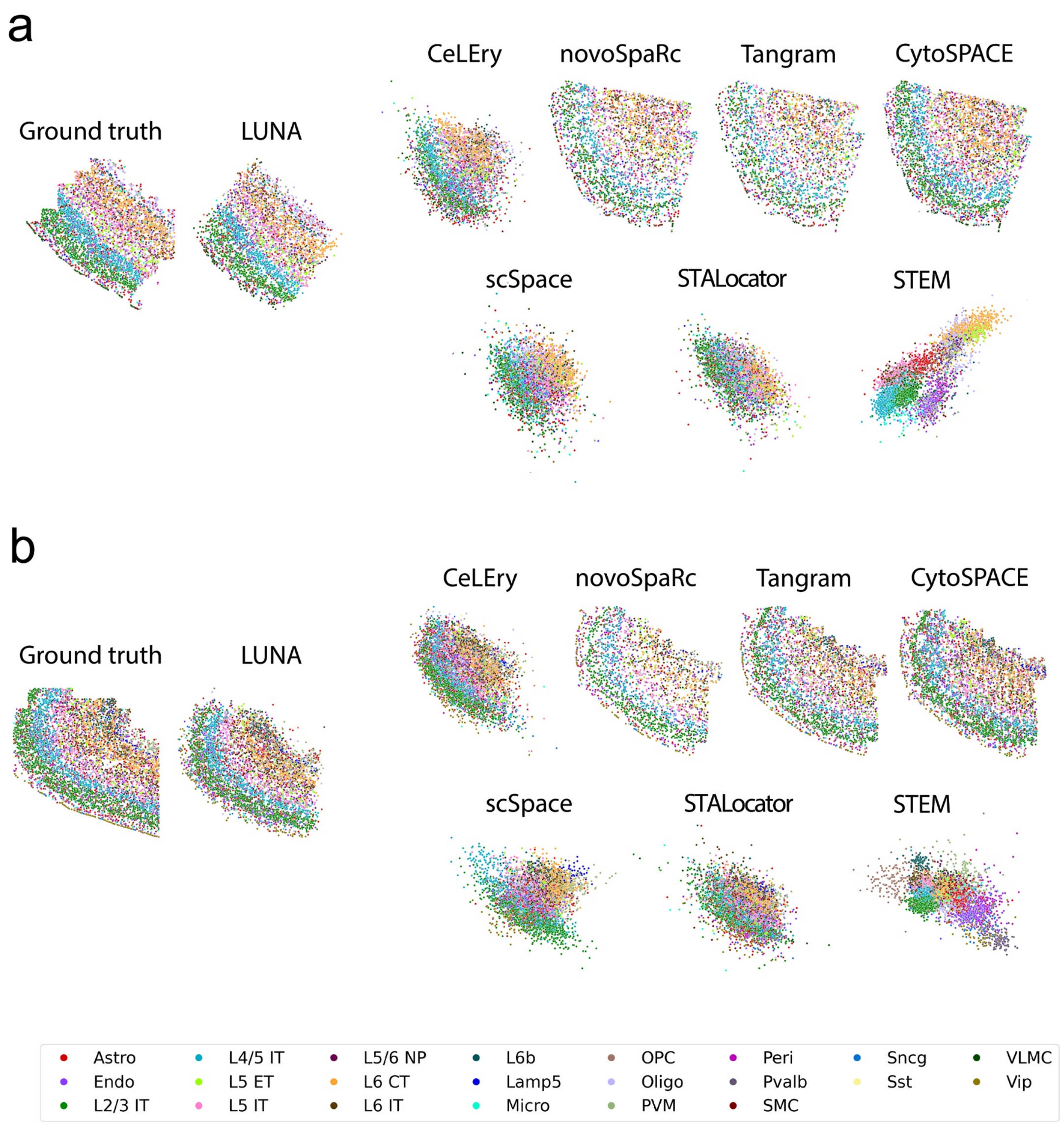

**Supplementary Figure 14:** Visualization of ground-truth locations and predictions using LUNA and alternative baselines for **(a)** one example slice ($5,235$ cells) using randomly selected reference slice and **(b)** one example slices ($5,024$ cells) using the best matched slice as a reference for alternative baselines. Colors denote cell class labels.

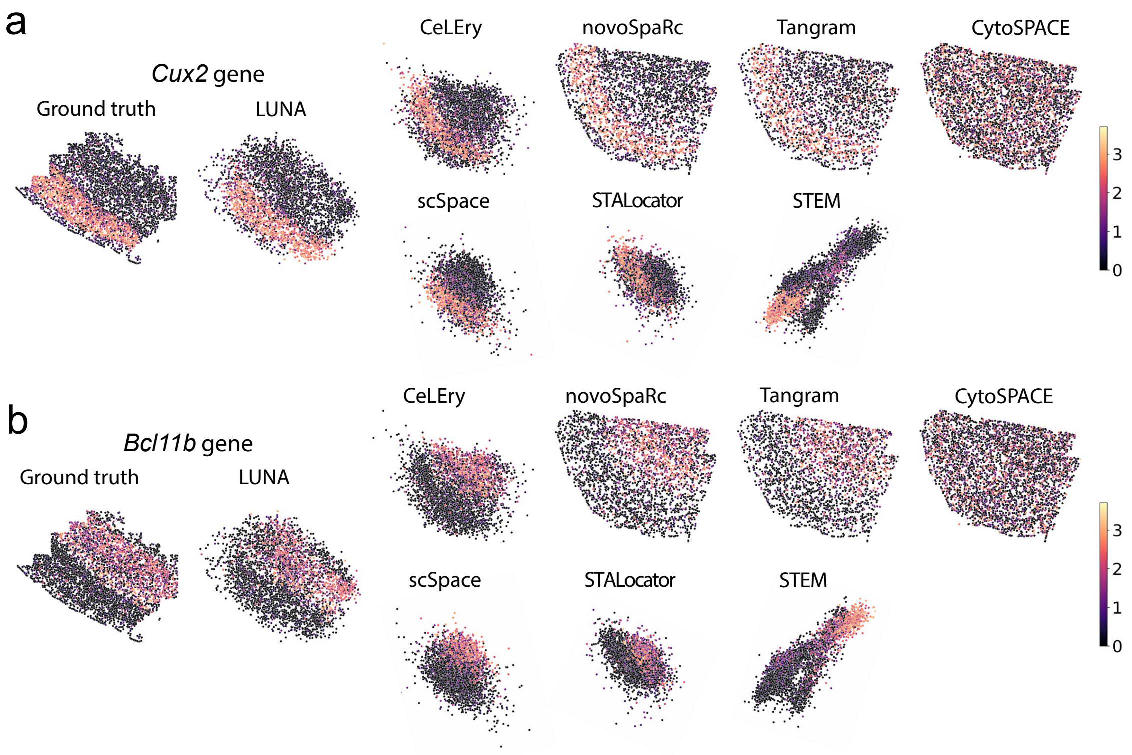

**Supplementary Figure 15:** Comparison of spatial expression patterns of the genes **(a)** *Cux2* (a marker for layer 2/3 intratelencephalic neurons, L2/3 IT) and **(b)** *Bcl11b* (a marker for layer 6 corticothalamic neurons, L6 CT). For each gene, the ground truth cell locations are shown in the leftmost plot, followed by the predictions made by LUNA and alternative baselines.

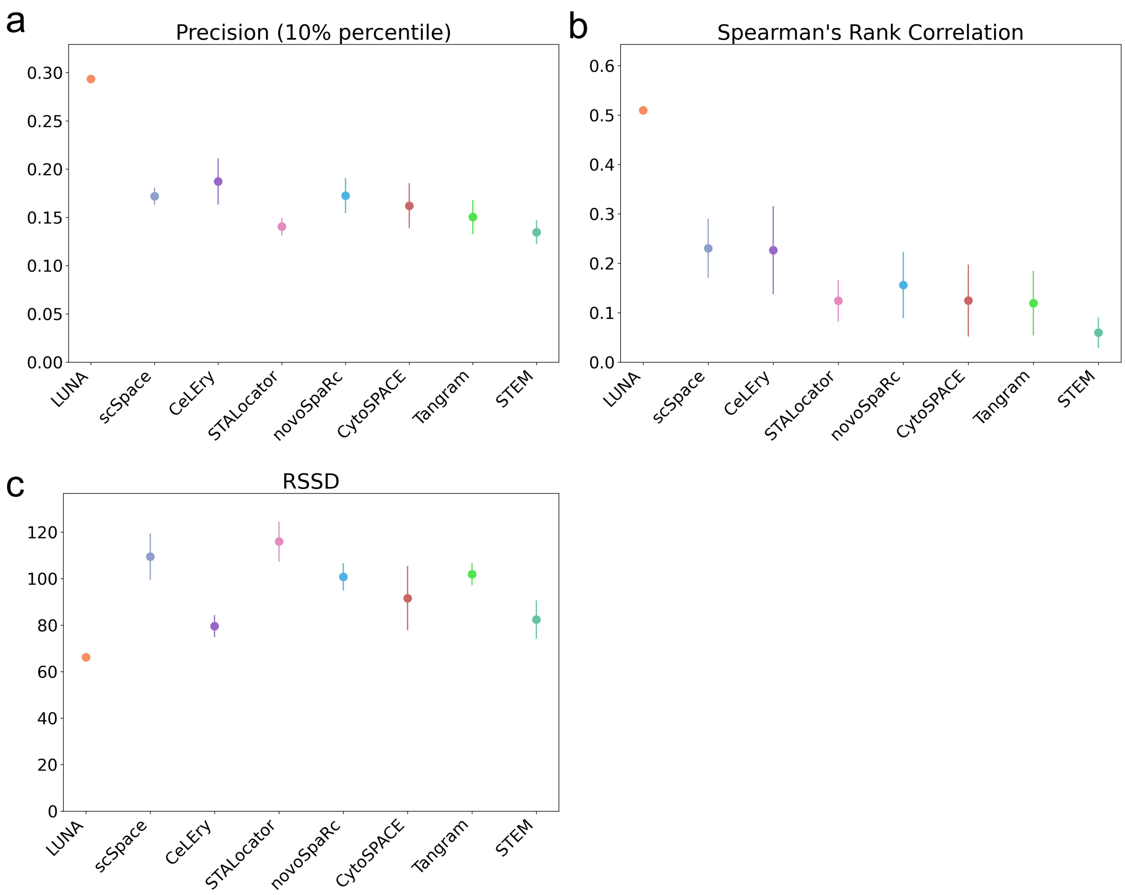

**Supplementary Figure 16:** Evaluation of the sensitivity of alternative methods to the choice of reference slice. We selected 5 different slices as a reference training slice and evaluated performance on a fixed test slice (n=5 slices). We computed the average performance of these methods across different reference slices. We evaluated performance based on **(a)** precision at the 10% percentile threshold for predicting the closest cell pairs as contacts, **(b)** Spearman's rank correlation (SRC) across all cells within the same slice, and **(c)** Root Sum Square Deviation (RSSD) between the ground truth and predicted spatial coordinates. Error bars represent standard deviation across 5 slices. For precision and SRC higher values indicate better performance. For RSSD, lower values indicate better performance.

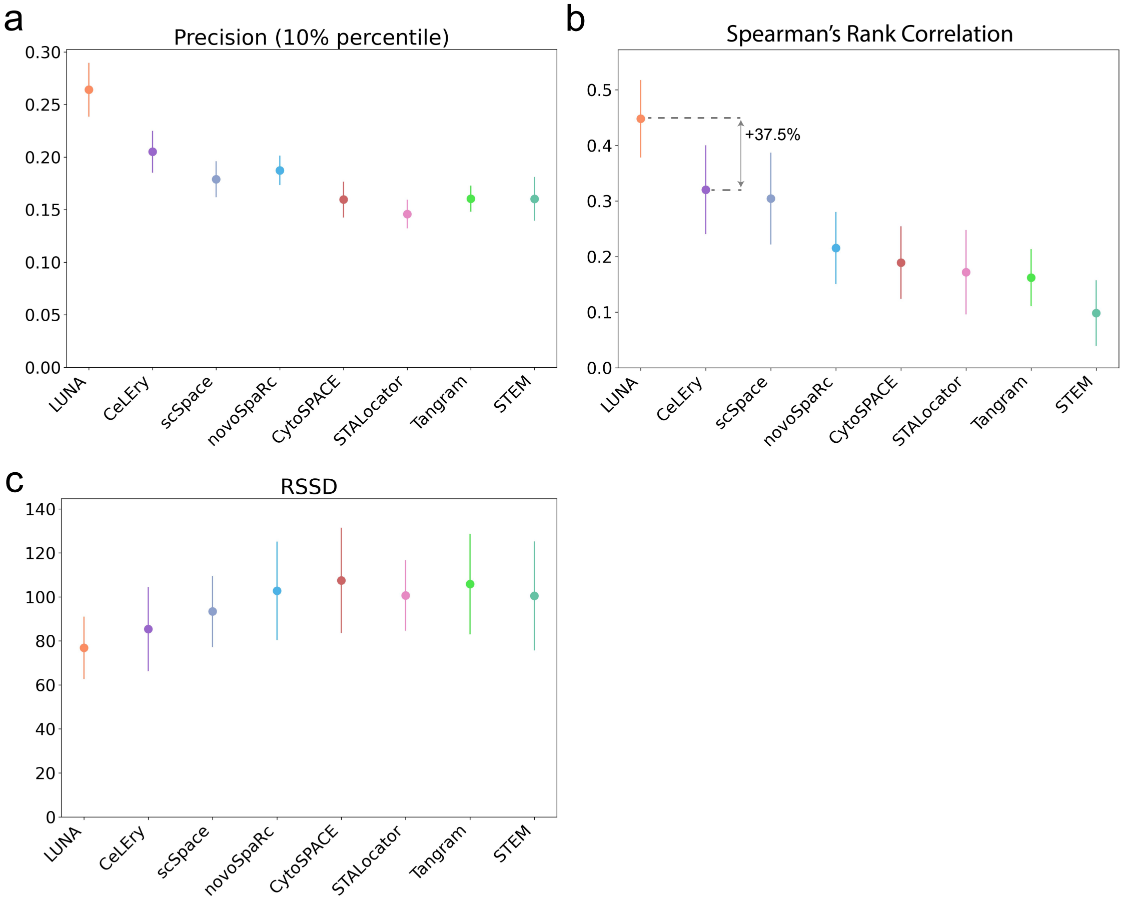

**Supplementary Figure 17:** Evaluation of alternative methods using the best match of the reference slice. For each fixed test slice, the most similar slice from the train set (including 33 slices) was selected by computing the cosine similarity of the cell type distributions. The average performance of these methods was computed across all slices from an unseen mouse, comprising 31 slices and 118,036 cells (n=31 slices). We evaluated performance based on **(a)** precision at the 10% percentile threshold for predicting the closest cell pairs as contacts, **(b)** Spearman's rank correlation (SRC) across all cells within the same slice, and **(c)** Root Sum Square Deviation (RSSD) between the ground truth and predicted spatial coordinates. Error bars represent standard deviation across 31 slices. For precision and SRC higher values indicate better performance. For RSSD, lower values indicate better performance.

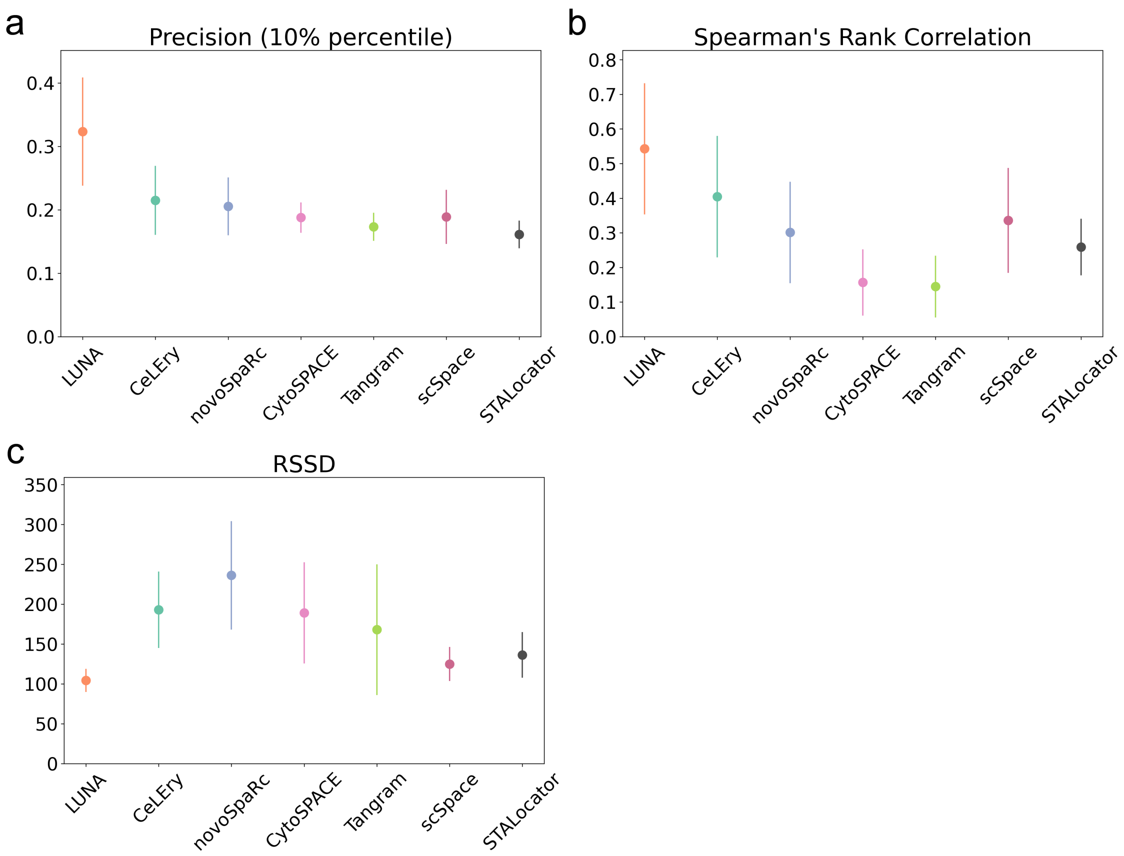

**Supplementary Figure 18:** Performance comparison on the cross-animal generalization task between LUNA and alternative baselines on the MERFISH ABC Atlas. Models were trained on slices from one mouse and evaluated on all slices from an unseen mouse. We evaluated performance based on **(a)** precision at the 10% percentile threshold for predicting the closest cell pairs as contacts, **(b)** Spearman's rank correlation (SRC) across all cells within the same slice, and **(c)** Root Sum Square Deviation (RSSD) between the ground truth and predicted spatial coordinates. Error bars represent standard deviation across 66 slices (n=66 slices). For precision and SRC higher values indicate better performance. For RSSD, lower values indicate better performance.

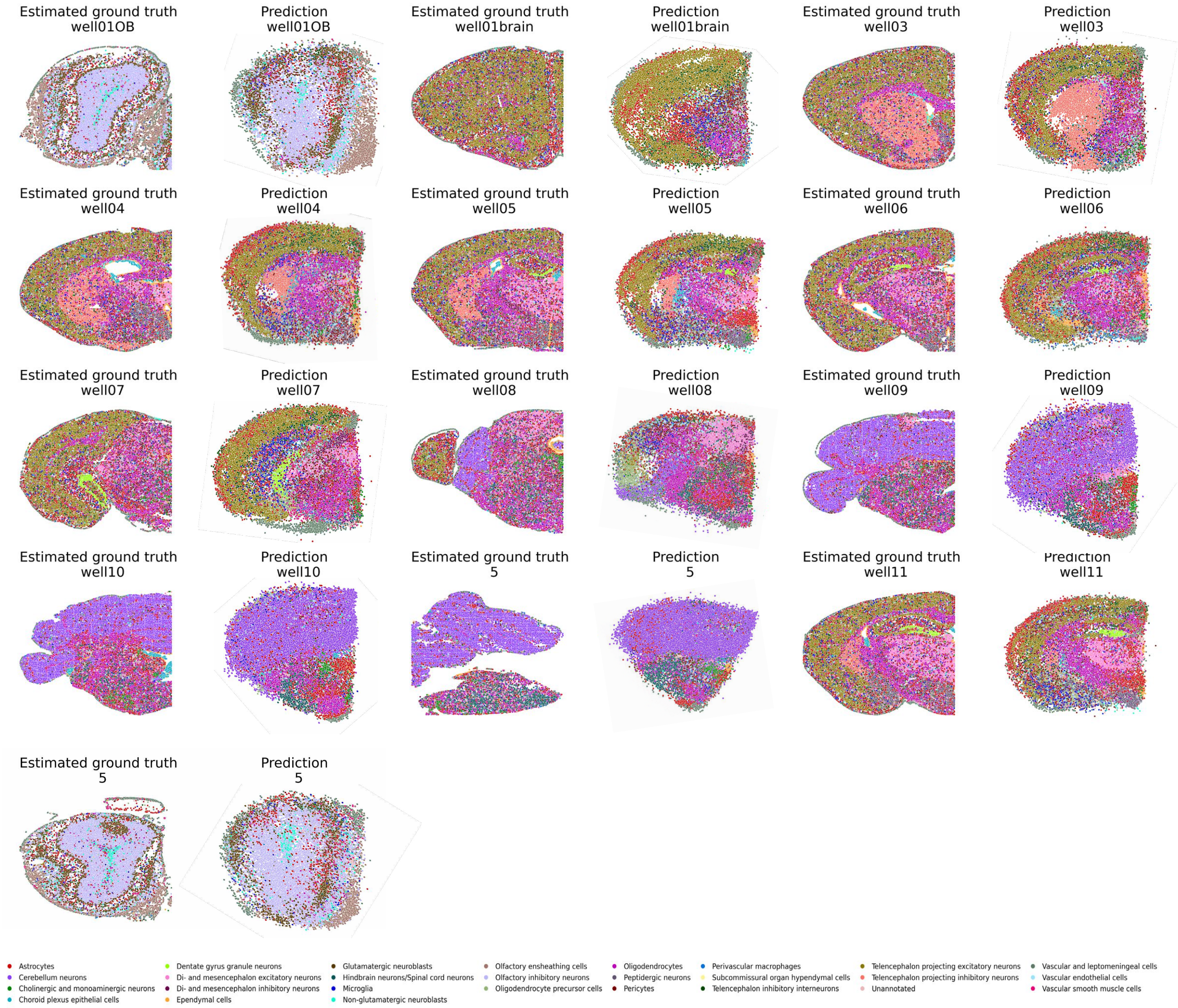

**Supplementary Figure 19:** Tissue reconstruction of the scRNA-seq atlas of the mouse central nervous system atlas [3] across different slices using LUNA. Cells are colored based on the main cell types (26 types). LUNA was trained on all the cells from Animal 1 of the ABC atlas (2.85 million cells) and applied to generate cell locations for a scRNA-seq atlas from the mouse central nervous system (1.08 million cells, 13 coronal slices). Ground truth spatial locations are estimated by integrating the CNS scRNA-seq dataset with the STARmap PLUS dataset [4].

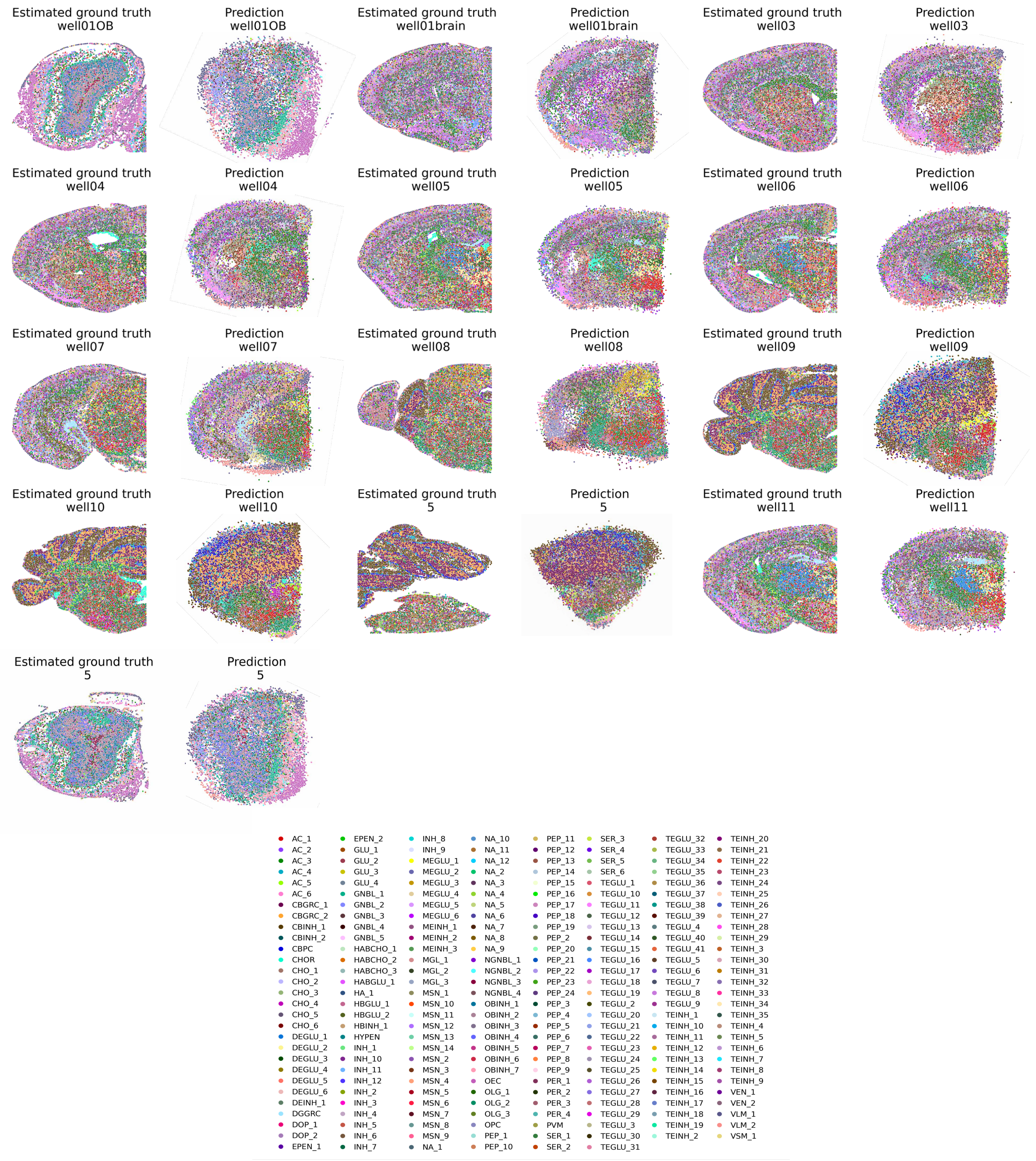

**Supplementary Figure 20:** Tissue reconstruction of the scRNA-seq atlas of the mouse central nervous system atlas [3] across different slices using LUNA. Cells are colored based on the cell sub-molecular types (216 types). LUNA was trained on all the cells from Animal 1 of the ABC atlas (2.85 million cells) and applied to generate cell locations for a scRNA-seq atlas from the mouse central nervous system (1.08 million cells, 13 coronal slices). Ground truth spatial locations are estimated by integrating the CNS scRNA-seq dataset with the STARmap PLUS dataset [4].

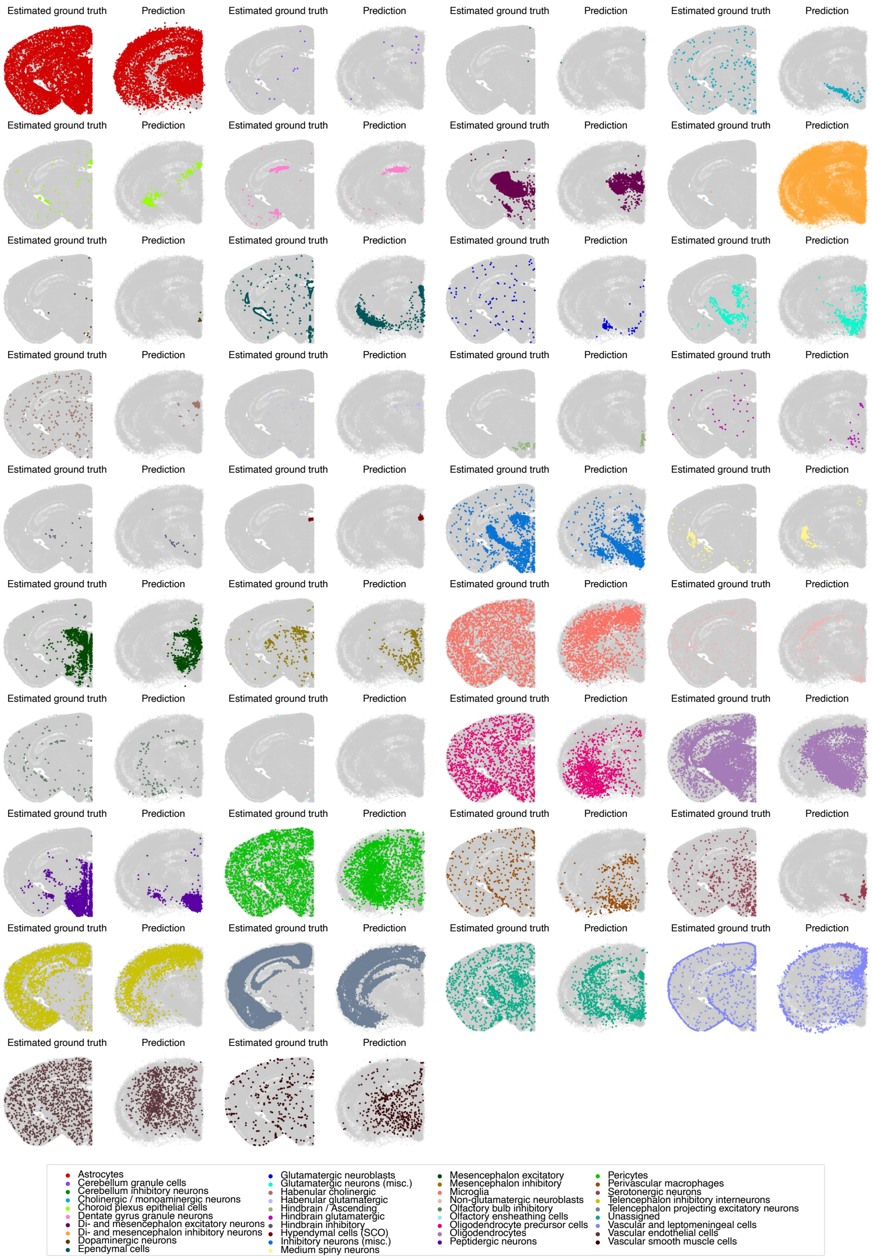

**Supplementary Figure 21:** Tissue reconstruction of a slice from the scRNA-seq mouse central nervous system atlas using LUNA. We display LUNA's performance across various main cell types. The left plot illustrates estimated ground truth locations from aligning th5e scRNA-seq and STARmap atlases, while the right plot features LUNA's predictions. Cells from other main cell types are shown in gray.

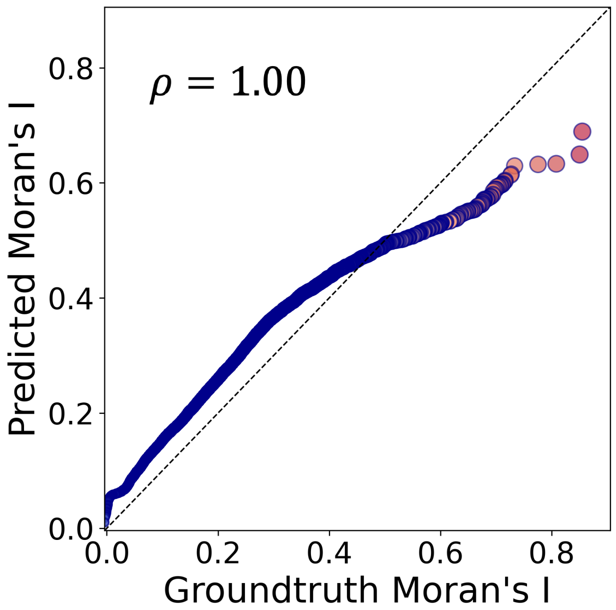

**Supplementary Figure 22:** Spatial autocorrelation computed using the Moran's I on the ground truth locations and locations predicted by LUNA on one example slice from scRNA-seq CNS Atlas. Each point represents a gene and all $11,844$ genes are visualized. Genes are colored on a gradient from blue (low) to red (high) based on their ground truth Moran's I value. Points closer to the diagonal indicate better preservation of spatial patterns by LUNA. The Pearson's correlation coefficient is reported on the top left.

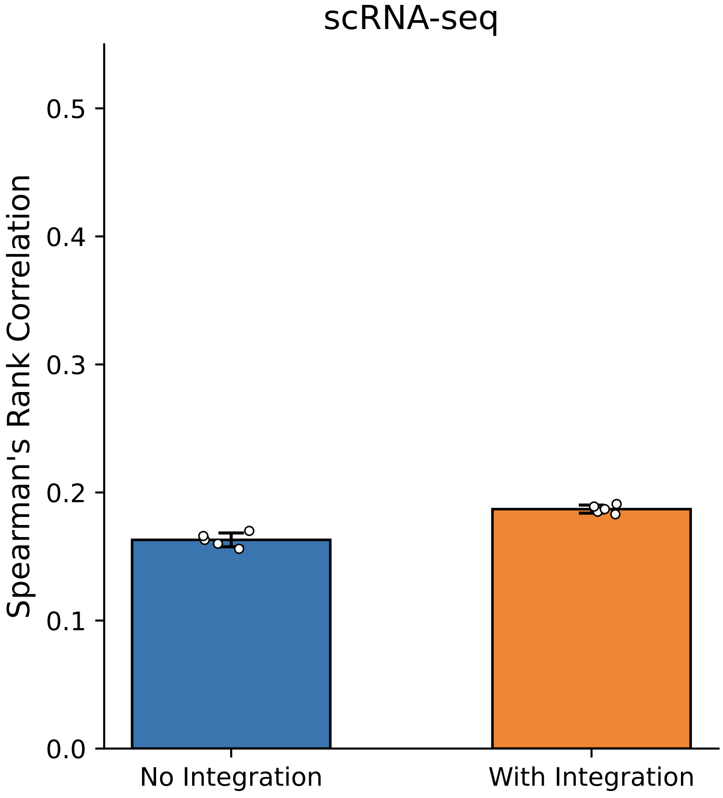

**Supplementary Figure 23:** Analysis of batch effects on LUNA's reconstruction performance. We compare the average Spearman's rank correlation between predicted and true pairwise distances with and without integration for the CNS scRNA-seq atlas. Data integration was performed using Harmony [5] to align dissociated single-cell data with a reference spatial atlas prior to training. Integration leads to a modest improvement in reconstruction accuracy for scRNA-seq atlas, highlighting the benefit of mitigating batch effects. Error bars indicate standard deviation across five runs with different random sampling seeds, while overlaid points represent the individual runs (n=5 runs).

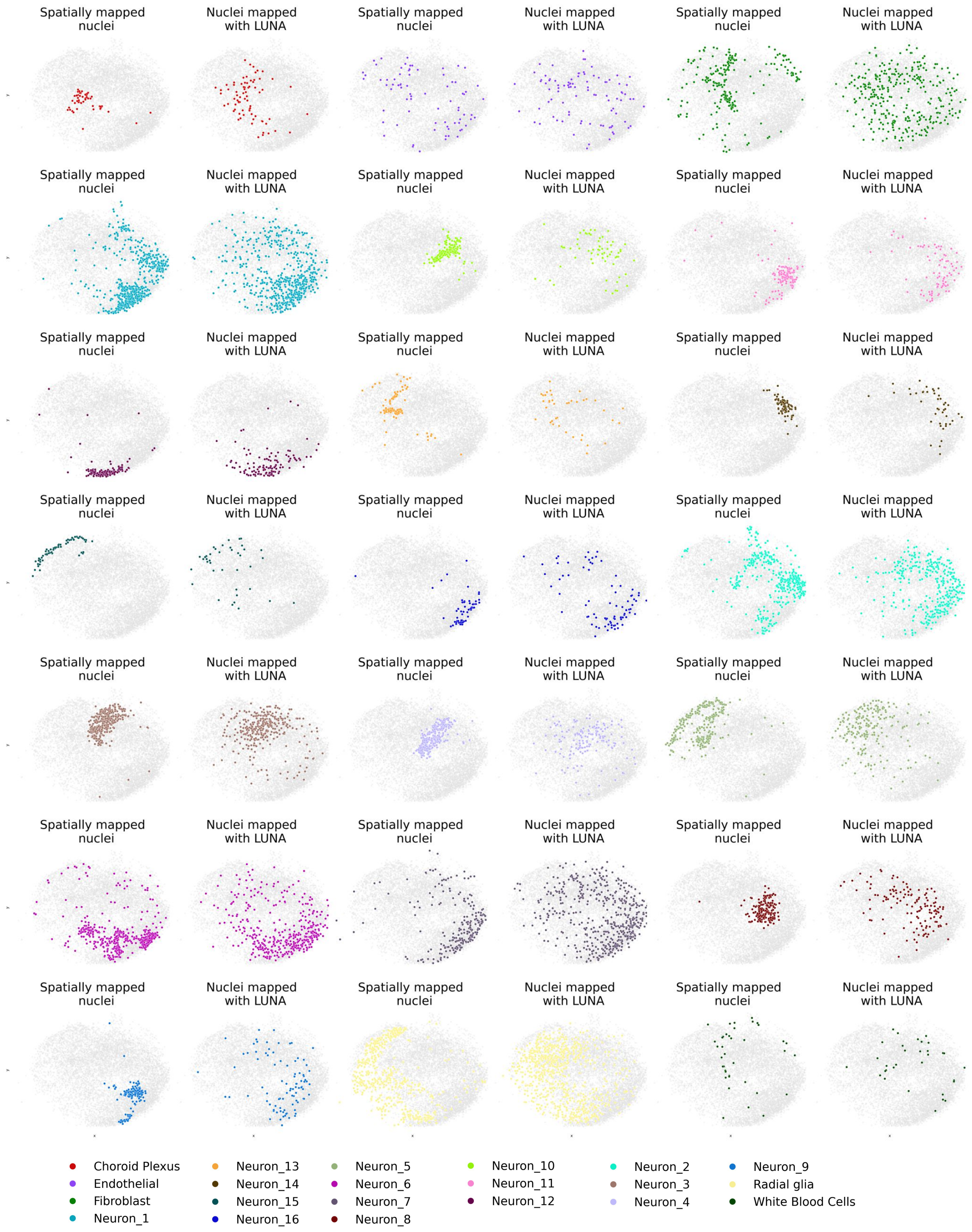

**Supplementary Figure 24:** Applying LUNA to enrich Slide-tags data by mapping spatially unmapped nuclei. Nuclei mapped by Slide-tags (left) and additional nuclei mapped by LUNA for all cell types from Slide-tags mouse E14 tissue. All other cells are shown in gray color.

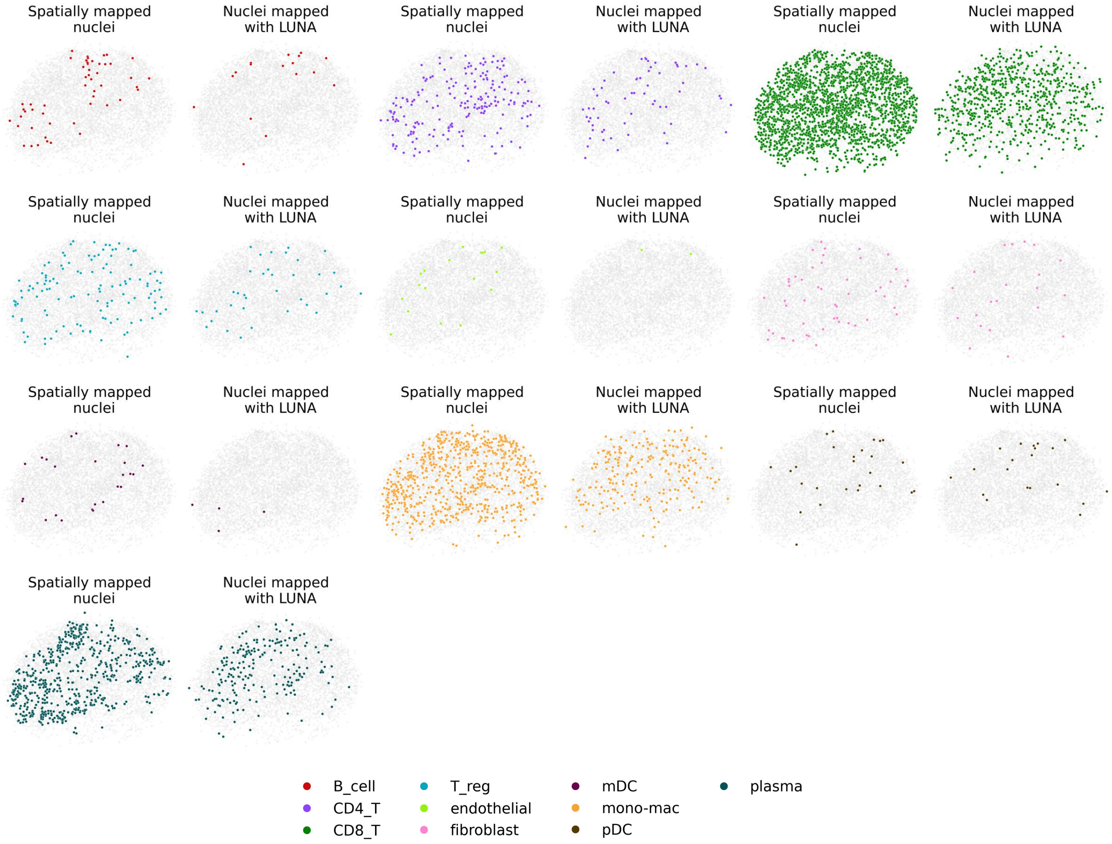

**Supplementary Figure 25:** Applying LUNA to enrich Slide-tags data by mapping spatially unmapped nuclei. Nuclei mapped by Slide-tags (left) and additional nuclei mapped by LUNA for all cell types from Slide-tags human melanoma tissue. All other cells are shown in gray color.

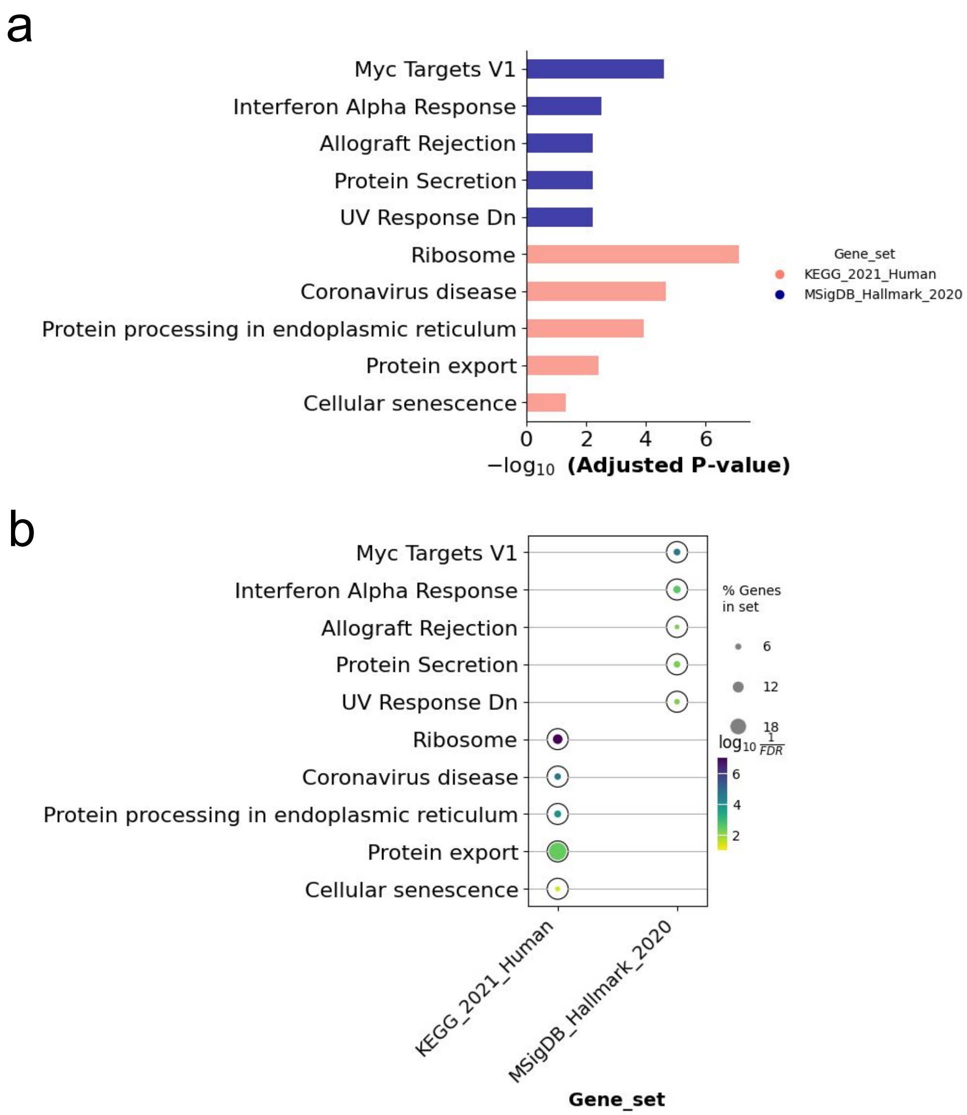

**Supplementary Figure 26:** Gene Ontology (GO) enrichment analysis was conducted on spatially variable genes uniquely identified using the LUNA-enriched melanoma tissue from Slide-tags using GSEApy package [24]. This analysis highlighted statistically significant GO terms (FDR-corrected $p$-value $< 0.05$) across the KEGG [25] and MSigDB [26] databases. **(a)** Top five significant GO terms with their FDR-corrected $p$-values. **(b)** Significant GO terms annotated by dots. The larger the dots, the greater overlap between the GO sets and the uniquely detected genes. Intensity of the dot color corresponds to the statistical significance of each GO term, with darker shades indicating higher levels of significance.

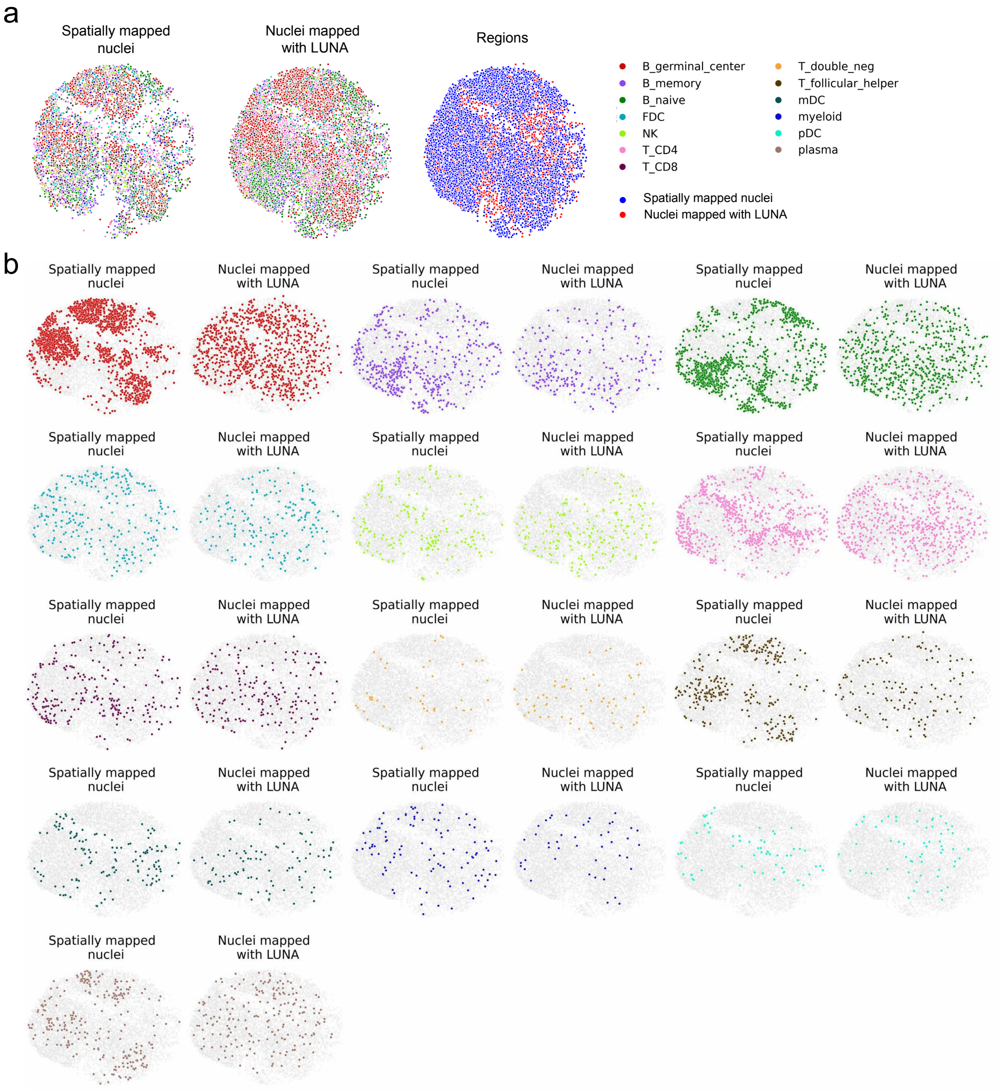

**Supplementary Figure 27:** **(a)** LUNA was applied to infer locations for the Slide-tags human tonsil tissue. The left panel shows the spatially mapped nuclei using Slide-tags, the middle panel shows the combined locations of nuclei mapped by LUNA and those spatially mapped by Slide-tags with cells colored according to their cell type annotation. The right panel differentiates the source of cell locations: nuclei mapped by Slide-tags (blue) and those mapped by LUNA (red). **(b)** Nuclei mapped by Slide-tags (left) and nuclei mapped by LUNA for all cell types from Slide-tags for human tonsil tissue. All other cells are shown in gray color.

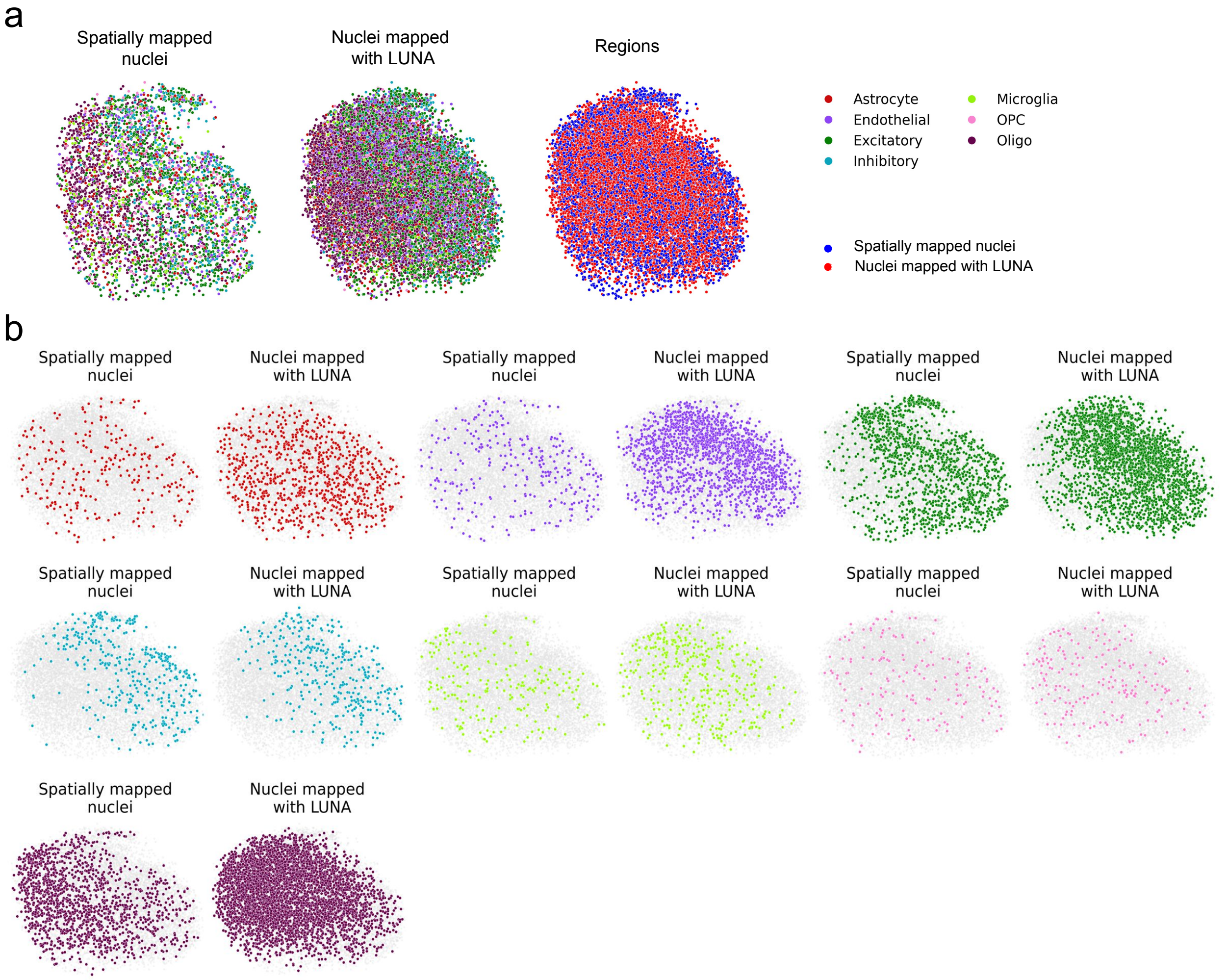

**Supplementary Figure 28:** **(a)** LUNA was applied to infer locations for the Slide-tags human cortex tissue. The left panel shows the spatially mapped nuclei using Slide-tags, middle panel shows the combined locations of nuclei mapped by LUNA and those spatially mapped by Slide-tags with cells coloured according to their cell type annotation. The right panel differentiates the source of cell locations: nuclei mapped by Slide-tags (blue) and those mapped by LUNA (red). **(b)** Nuclei mapped by Slide-tags (left) and nuclei mapped by LUNA for all cell types from Slide-tags for human cortex tissue. All other cells are shown in gray color.

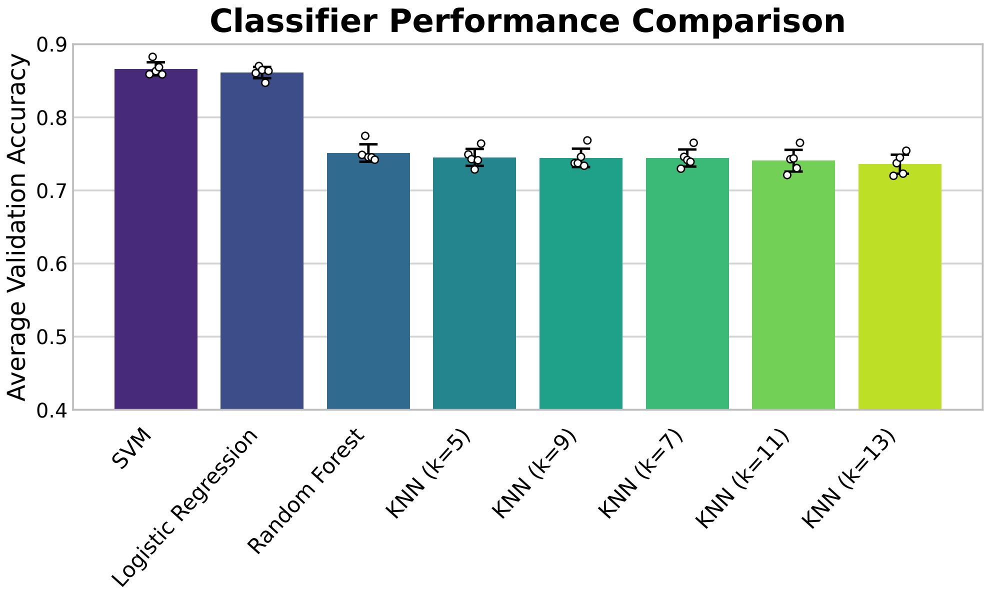

**Supplementary Figure 29:** Classifier performance comparison on the Slide-tags E14 sample. We evaluate several classifiers including SVM, logistic regression, random forest, and K-nearest neighbors with varying values of $k$ based on their average validation accuracy across 5-fold cross-validation, while overlaid points represent the individual runs. (n=$5$ runs).. Error bars indicate standard deviation across the cross-validation folds. Linear SVM achieved the highest classification accuracy across 5-fold cross-validation, with an accuracy of $86.63\%$, slightly outperforming the logistic regression model which achieves accuracy of $86.11\%$.

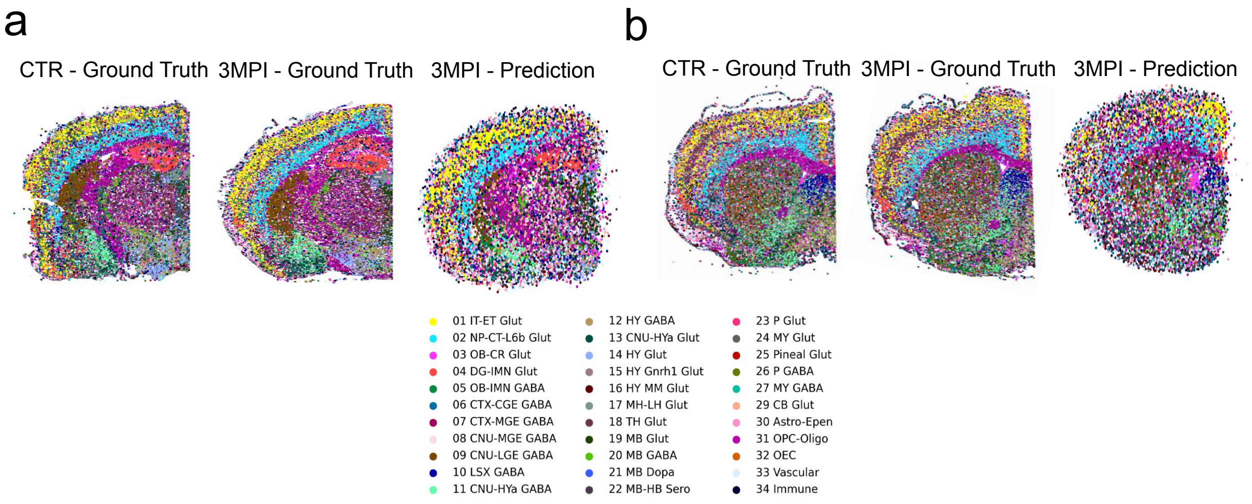

**Supplementary Figure 30:** Tissue reconstruction of the 3MPI sample using LUNA on **(a)** a posterior section targeting the amygdala ($-1.70$ mm relative to bregma) and **(b)** an anterior section targeting the striatum ($+0.6$ to $-1$ mm relative to bregma). For each section, we visualize its spatial distribution in a training sample (CTR, left), a 3MPI sample (middle), and the spatial coordinates predicted by LUNA for the 3MPI sample (right). Cells are colored by their cell class.

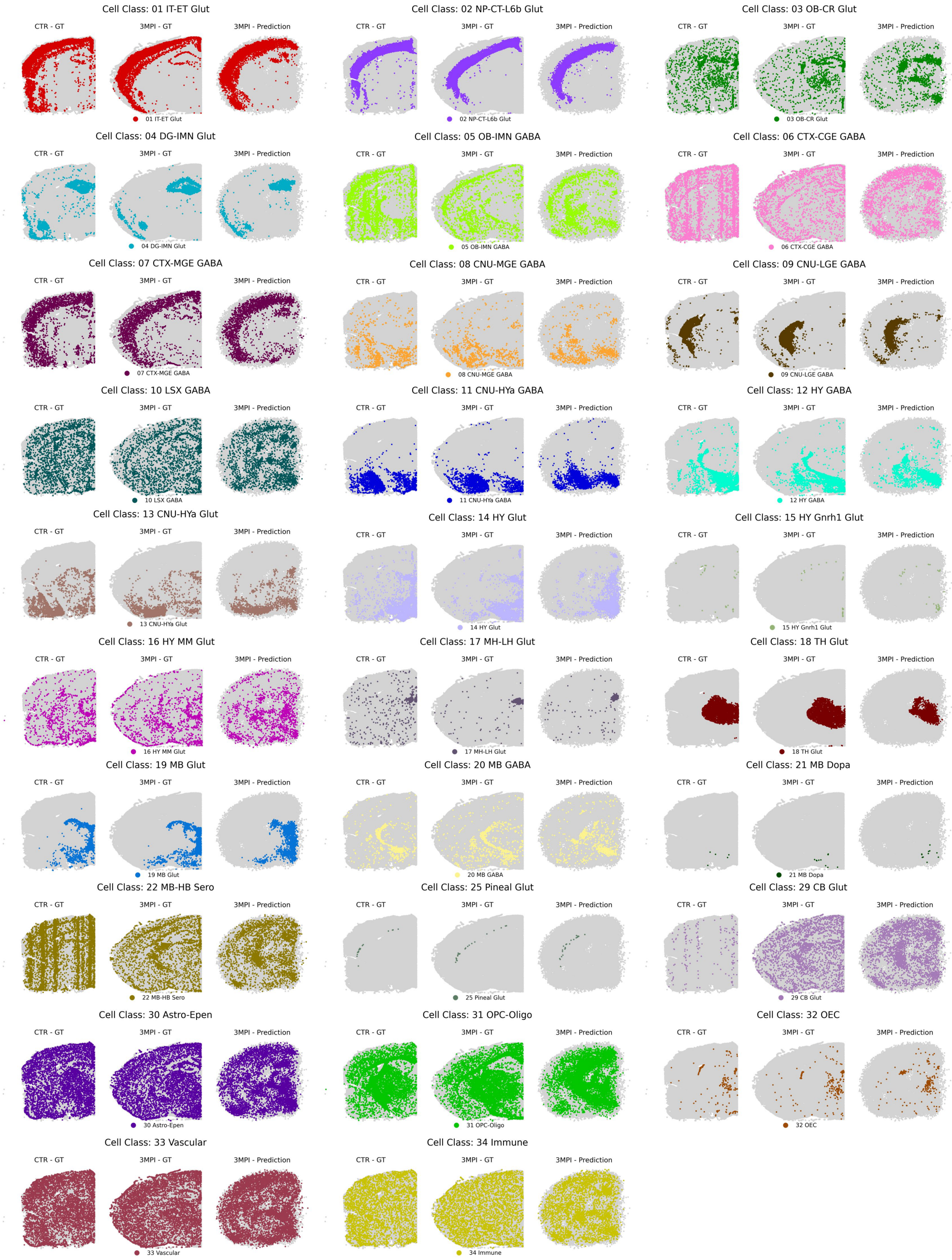

**Supplementary Figure 31:** Spatial distributions of all cell classes on the posterior section targeting the amygdala ($-1.70$ mm relative to bregma). For each cell class, we visualize its spatial distribution in a training sample (CTR, left), a 3MPI sample (middle), and the spatial coordinates predicted by LUNA for the 3MPI sample (right). Gray points indicate all other cell types for reference.

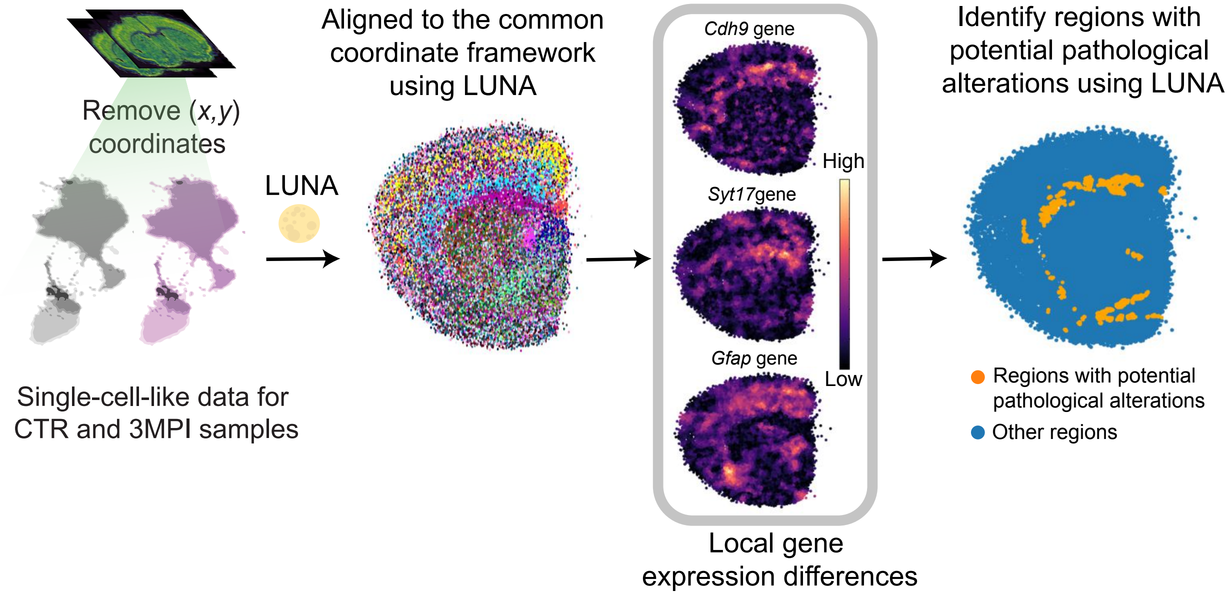

**Supplementary Figure 32:** Cells exhibiting differential transcriptomic profiles between anterior CTR and 3MPI samples targeting the striatum ($+0.6$ to $-1$ mm relative to bregma) identified from LUNA's predictions. We removed spatial coordinates from the Xenium dataset and input single-cell-like gene expression data from the CTR and 3MPI sections into LUNA. During inference, LUNA aligns these samples to a common coordinate framework by jointly embedding them. We visualized the genes *Cdh9*, *Syt17*, and *Gfap*, which were selected for their local spatial differential expression in the striatum region. Cells were colored by their local expression differences, with lighter shades indicating greater local shifts. To identify regions with potential pathological alterations, we further computed a unified score across genes with high distributional shifts between the 3MPI and CTR samples. Cells with significant local transcriptomic differences were labeled in yellow, while all others were shown in blue.

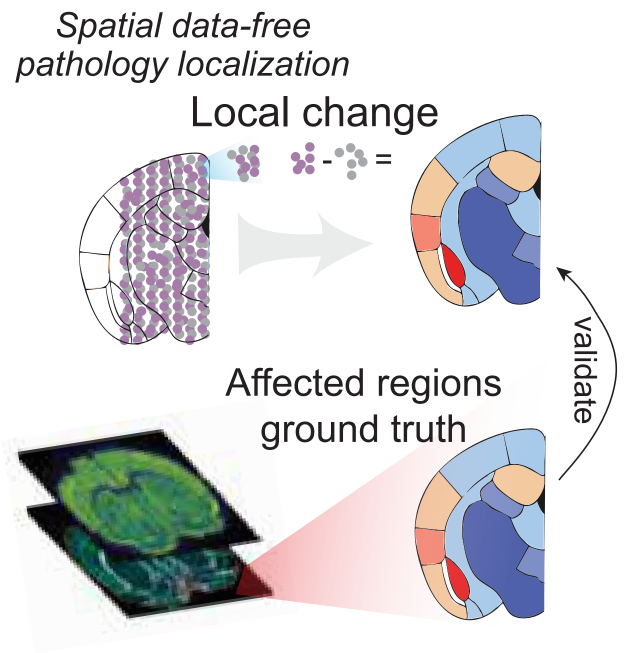

**Supplementary Figure 33:** Local gene expression shifts were quantified at the single-cell level to identify regions that potentially associate with pathological alterations in 3MPI samples, which are potentially affected by pathology. These predictions were validated on the same sections using immunostaining against $\alpha$-syn.

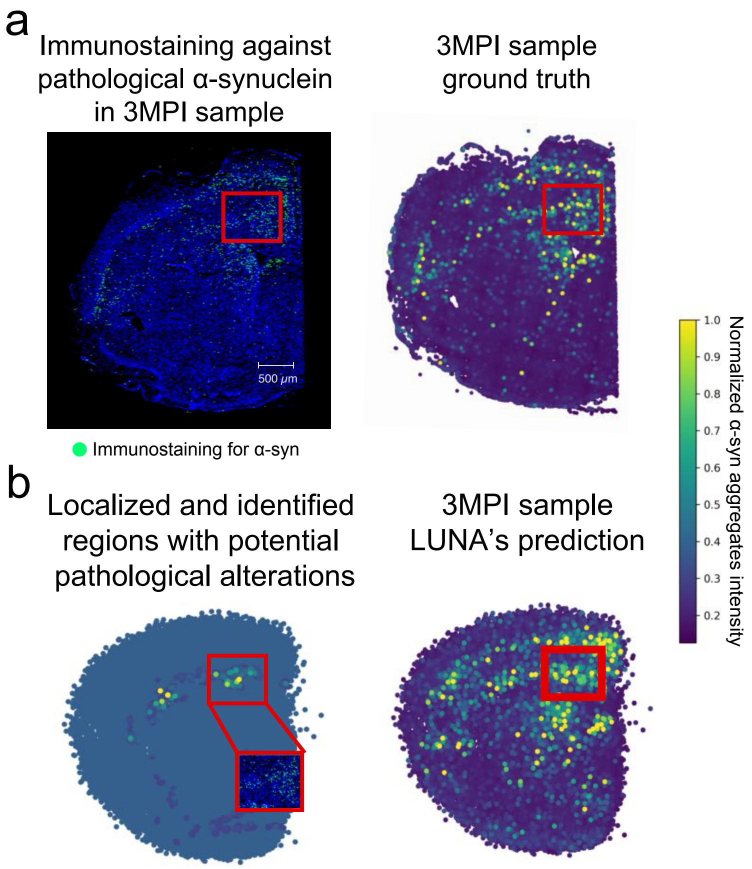

**Supplementary Figure 34:** **(a)** Immunostaining for $\alpha$-syn pathology for a 3MPI anterior section targeting the striatum ($+0.6$ to $-1$ mm relative to bregma) on the left which shows $\alpha$-syn in green, with each dot representing an individual protein aggregate. Ground truth cell locations for the same sample are shown on the right, with each cell colored by the intensity of $\alpha$-syn fibril staining. Scale bar is on the panel. **(b)** LUNA enables spatial data-free identification of brain regions with spatial gene expression dysregulation that correlate with $\alpha$-syn pathology. Cells that are identified with potential pathological alterations between 3MPI and CTR (left) and all cells (right) are positioned according to LUNA's predicted locations and colored by $\alpha$-syn fibril staining intensity. A close-up view highlights the immunostaining of the regions identified by LUNA that potentially associate with pathological alterations, overlaid above the left plot. Immunostaining was performed on two 3MPI sections with similar results.

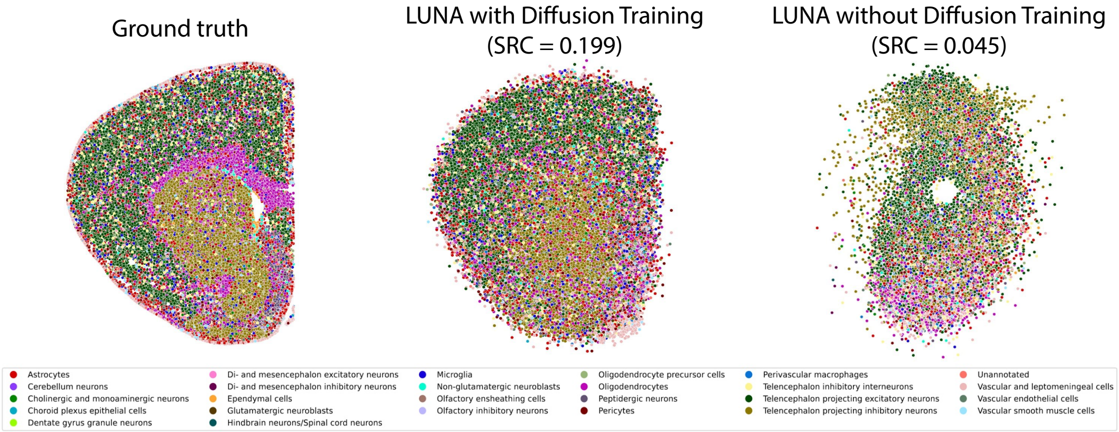

**Supplementary Figure 35:** Impact of diffusion training on spatial reconstruction performance on generalizing to the STARmap dataset. Cells are colored by cell type for the ground truth (left), LUNA with diffusion training (middle), and LUNA without diffusion training (right). LUNA with diffusion training better recovers the anatomical structure and spatial organization of the tissue, as reflected by a higher Spearman rank correlation (SRC = $0.199$) between predicted and ground truth spatial coordinates. In contrast, removing the diffusion component results in a less coherent reconstruction with substantially lower spatial correspondence (SRC = $0.045$), highlighting the importance of the diffusion-based modeling in LUNA.

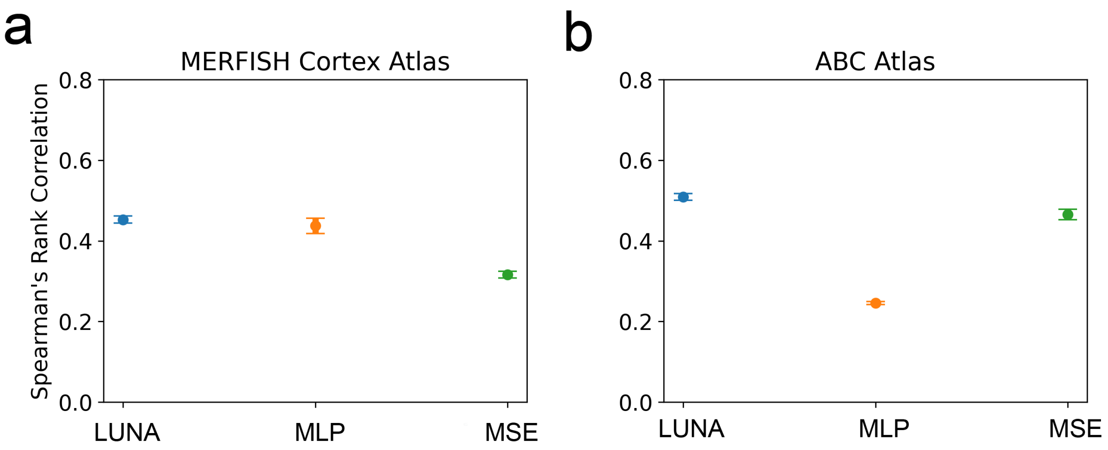

**Supplementary Figure 36:** Performance on the **(a)** MERFISH Cortex Atlas and **(b)** the ABC Atlas under the cross animal generalization scenario, measured by Spearman's rank correlation between predicted and true pairwise distances. We compare the full LUNA model (self-attention + rotation-invariant loss) against ablations that replace the attention-based decoder with a simple MLP (MLP), replace the SE($k$)-invariant loss with a mean squared error loss (MSE). Results are averaged over all test slices, with five independent samplings using different sample seeds per slice. Error bars represent the variance on the average performance across these five runs (n=5 runs).

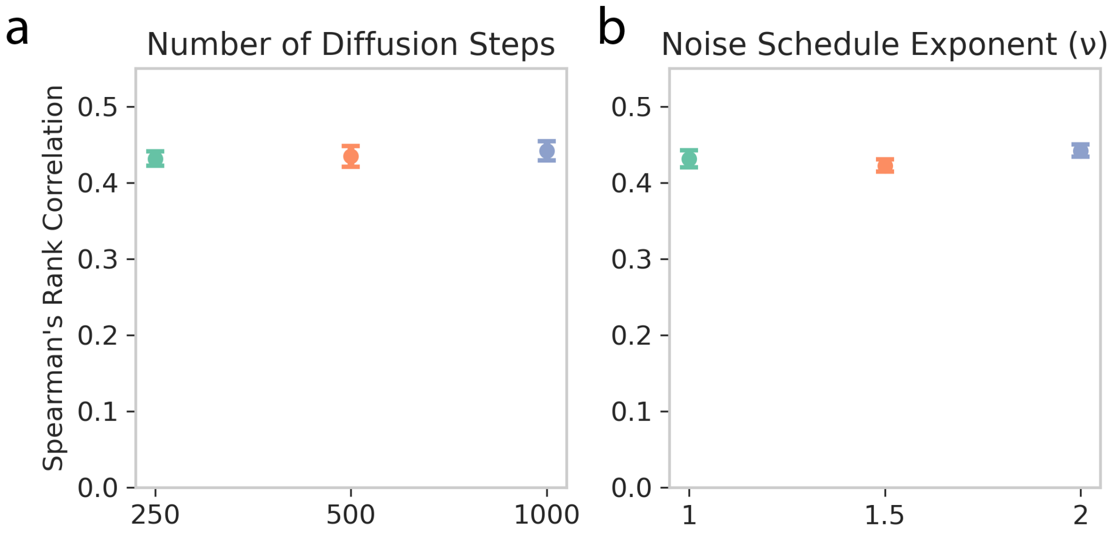

**Supplementary Figure 37:** Sensitivity analysis of key hyperparameters of diffusion model in LUNA, evaluated on the MERFISH mouse cortex dataset under the cross animal generalization scenario using Spearman's rank correlation. **(a)** Number of diffusion steps; **(b)** noise schedule exponent, which controls the shape of the noise decay across diffusion timesteps. Results are averaged over all test slices, with five independent samplings using different sample seeds per slice. Error bars represent the variance on the average performance across these five runs (n=5 runs).

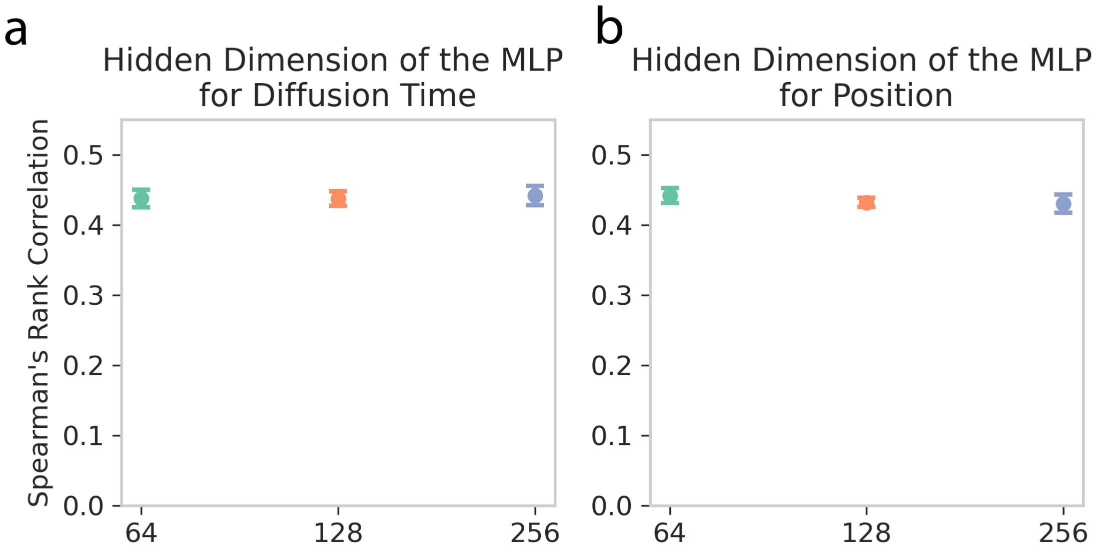

**Supplementary Figure 38:** Sensitivity analysis of key hyperparameters of diffusion model in LUNA, evaluated on the MERFISH mouse cortex dataset under the cross animal generalization scenario using Spearman's rank correlation. Hidden dimensions of the MLPs used for **(a)** diffusion time embedding and **(b)** position embedding. Results are averaged over all test slices, with five independent samplings using different sample seeds per slice. Error bars represent the variance on the average performance across these five runs (n=5 runs).

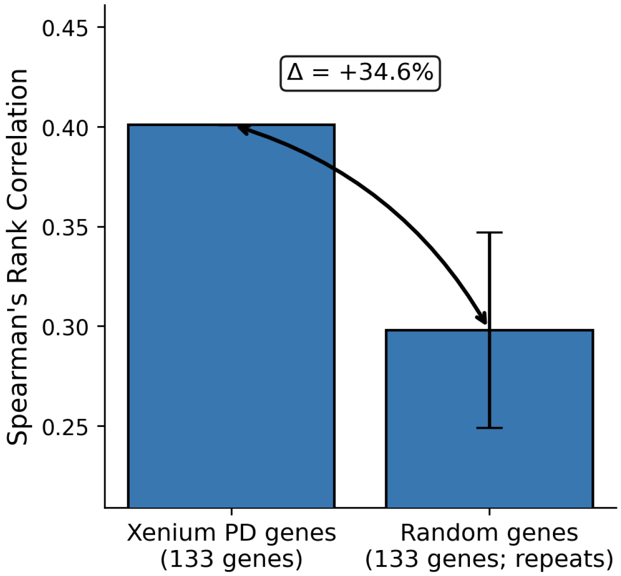

**Supplementary Figure 39:** The Xenium gene panel provides more predictive information than randomly selected gene panels. We compared models trained on the ABC Atlas using either the 133 genes from the Xenium PD panel or one of six randomly selected panels of equal size (n=6 independent random gene-panel runs). All sections from Animal 1 were used for training and Animal 2 for testing. Performance was evaluated by Spearman's rank correlation between predicted and ground truth spatial coordinates, averaged across test sections. Bar heights correspond to the mean Spearman's rank correlation across test sections and the error bar indicates standard deviation across six random runs. The Xenium PD panel (left) outperformed random panels (right) by $\Delta = 34.6\%$ relative to the mean performance of the random gene-panel runs.
